# Tree diversity and composition of cocoa agroforests in Côte d’Ivoire: a regional ecological mosaic, implications for cocoa landscapes sustainable management

**DOI:** 10.64898/2026.08.20.745955

**Authors:** Beda Innocent Adji, Assiri Assiri Alexis, Assi Maryse Evelyne, Aya Diane Larissa Houphouet, Kassin Koffi Emmanuel, Doffou Sélastique Akaffou

## Abstract

Cocoa (*Theobroma cacao* L.) is grown in Côte d’Ivoire (the world’s leading producer), almost always in association with trees, yet these cocoa-based agroforestry systems (cocoa AFS) remain characterized largely descriptively, without a robust inferential framework or modern diversity indices. We analyse a survey database covering 474 farms and 103 localities across the three main cocoa-producing zones of Côte d’Ivoire, locally known as « loops » (East/South-East, Centre/Centre-West, South/South-West; surveys 2013-2016), applying a renewed ecological and statistical framework: Hill numbers (*q* = 0, 1, 2), individual-based Hurlbert rarefaction, multiplicative *alpha/beta/gamma* diversity partitioning, ordination (*PCoA*) and *PERMANOVA* on Bray-Curtis matrices, Ward’s hierarchical clustering for a data-driven floristic typology, multivariate *MANOVA*, and linear mixed models. Across 99 adequately sampled localities (≥10 stems), mean taxonomic richness and the Shannon index differed significantly among zones (*MANOVA*: Wilks’ λ = 0.350; *F_8,180_* = 15.5; *p* < 0.001), including after standardizing sampling effort by rarefaction (*F* = 8.72; *p* < 0.001). Floristic composition differed significantly by zone (*PERMANOVA*, *pseudo-F* = 2.85; *p* = 0.001), though with uneven within-group dispersion (*PERMDISP*, *p* = 0.024) that calls for caution in interpretation. The Centre/Centre-West zone showed the highest *beta* diversity (inter-farm floristic turnover ≈ 16.4, versus 7.2 and 2.2 in the other two zones), revealing a highly heterogeneous mosaic of individual agroforestry choices rather than a homogeneous regional system. A typological classification based on floristic composition distinguishes four cocoa-AFS profiles, ranging from dense, diverse stands to impoverished systems with low densities of associated trees, partially but not fully overlapping with the zonal boundaries. At the scale of the three loops *(n* = 409 farms with both age and yield recorded), cocoa yield varied significantly by orchard age class (*Kruskal-Wallis*, *H* = 18.2; *p* < 0.001), following a bell-shaped profile peaking around 20-35 years, consistent with the intermediate-age yield optimum suggested qualitatively by the source study; this relationship, robust to log-transformation of yield and to the exclusion of extreme values, nonetheless remains of small magnitude (*R²* ≈ 1.5%) and confers no out-of-sample predictive power under cross-validation, illustrating the need to distinguish statistical significance from predictive utility. This reanalysis provides a reproducible and transferable statistical framework for characterizing West African cocoa AFS, and highlights the potential value of high *beta* diversity as a possible indicator of landscape resilience.

## 1. Introduction

Cocoa farming is the leading economic pillar of Côte d’Ivoire, which has supplied roughly 38 to 44% of the world’s cocoa bean output since 1978 (ICCO, 2025). This perennial crop is grown almost everywhere in association with fruit, forest, or exotic trees (spontaneously or deliberately) forming cocoa-based agroforestry systems (cocoa AFS). In Nair’s (1993) sense, these systems constitute an agrosilviculture combining trees and perennial crops, and they play a recognized role in diversifying farmers’ incomes, maintaining soil fertility, regulating microclimate, and conserving part of the residual forest biodiversity (Boffa, 1999; Rolando et al., 2014; Somarriba, 2007; Sonwa et al., 2003), even as cocoa farming remains the leading cause of deforestation in Côte d’Ivoire (Adji et al., 2020; Assiri et al., 2015; Kouamé et al., 2024; Kouassi et al., 2023; N’Guessan et al., 2026; Serge Cherry et al., 2011). Beyond Côte d’Ivoire, the ecological performance of cocoa agroforestry is a question of direct international relevance. An extensive body of literature spanning Africa, Latin America, and Southeast Asia has shown that shaded cocoa systems can reconcile production with the conservation of considerable tree, bird, and insect biodiversity, but that this potentially « win-win » outcome is highly sensitive to the intensity and structure of shade management, and can collapse under progressive shade removal and management intensification (Clough et al., 2011; Tscharntke et al., 2011). Rigorously characterizing the diversity, composition, and structure of the associated tree stratum is therefore not a peripheral descriptive exercise, but a prerequisite for assessing whether a given cocoa-producing region actually delivers on this claimed potential. This question has acquired renewed and immediate urgency since the adoption of the European Union’s « deforestation-free » products regulation (Union Européenne, 2023), whose due-diligence and traceability obligations for cocoa (now fully applicable to large and medium operators from 30 December 2026) increasingly require exporting countries to document the ecological state of their cocoa landscapes, including their associated tree component, at a level of statistical rigor that most existing national characterizations (aggregated and largely descriptive) do not yet offer. Recent empirical work conducted in Côte d’Ivoire further shows that farmers’ actual level of preparedness for these new obligations remains highly heterogeneous and depends strongly on their integration into collective support arrangements such as cooperatives (Moluh Njoya et al., 2025), which reinforces the value of a fine-grained ecological characterization capable of capturing this heterogeneity rather than masking it behind a national average.

It is precisely this gap between the political demand for rigorous, spatially resolved ecological evidence and the descriptive state of existing regional inventories that motivates the present reanalysis, which takes as its case study the most comprehensive characterization of Ivorian cocoa AFS to date: that of Adji et al. (2016), based on a survey of 474 cocoa farms and a floristic inventory of 103 villages and hamlets spread across the three major historical « loops » of Ivorian cocoa farming (East/South-East, Centre/Centre-West, South/South-West). This work established an inventory of more than a hundred associated taxa, described their uses, and produced a qualitative typology of cocoa AFS based on orchard age. It nonetheless presents important statistical limitations, common to many agroforestry characterization studies in West Africa: (i) diversity indices (Shannon, Simpson, Pielou’s evenness) were calculated only at the aggregated scale of the three loops, without replication or unambiguous significance testing; (ii) multivariate analyses (PCA, CA, hierarchical clustering) were conducted on only three observations (one per loop), which permits no statistically valid inference in the strict sense; (iii) no correction for sampling effort (rarefaction) was applied, even though the number of individuals recorded varied considerably from one zone to another; and (iv) floristic composition as such (beyond synthetic indices) was never analysed using community ecology methods (ordination, PERMANOVA), even though these have become standard in the international ecological literature on shaded perennial crops (Anderson, 2006; Chao et al., 2014; Clough et al., 2011; Jost, 2006). We propose here a complete ecological and statistical analysis of this original dataset, drawing on a renewed methodological framework that is now standard in community ecology and increasingly applied to cocoa and coffee agroforestry systems elsewhere in the tropics: Hill diversity numbers of order *q* = 0, 1, 2 (Chao et al., 2014; Hill, 1973; Jost, 2006), which unify richness, the Shannon index, and the Simpson index within a single abundance-weighted family of measures; individual-based Hurlbert rarefaction (Hurlbert, 1971) for fairly comparing sites sampled with unequal effort; the multiplicative partitioning of diversity into *alpha* (local), *beta* (turnover), and *gamma* (regional) components (Jost, 2007; Whittaker, 1972) ; the analysis of floristic composition through ordination of Bray-Curtis dissimilarities (*PCoA*) and a *PERMANOVA* test (Anderson, 2001) ; an agglomerative hierarchical classification based on floristic composition rather than on aggregated indices, allowing us to build a genuinely data-driven typology of cocoa AFS; and linear mixed models together with a multivariate *MANOVA* to rigorously test the effect of production zone on a vector of diversity indices, accounting for the hierarchical spatial structure of the data (locality nested within sub-zone nested within loop).

The specific objectives of this study are to: (i) statistically quantify and compare, at the locality scale (the replicated ecological unit, *n* = 99-119 sites), the *alpha*, *beta*, and *gamma* diversity of cocoa-associated trees across the three cocoa loops; (ii) test whether the floristic composition of cocoa AFS differs significantly among zones and derive a data-driven typology independent of simple administrative boundaries; (iii) quantitatively describe the vertical stratification of agroforestry stands using a subsample of plots inventoried by stratum; and (iv) re-evaluate, using formal statistical tests, the hypothesis of a relationship between cocoa orchard age and yield, as suggested descriptively in the source study. By combining these analyses, this paper aims to provide not only a substantial update of knowledge on Ivorian cocoa AFS, but also a reproducible and exportable methodological framework, directly relevant to the emerging evidentiary requirements of « deforestation-free » cocoa policies, and transferable to other cocoa-producing regions of West and Central Africa facing similar limitations in their national agroforestry inventories.

## 2. Material and methods

This study is a reanalysis of raw survey and floristic inventory data on associated species collected by Adji et al. (2016) as part of a Master’s thesis in Agroforestry, using survey sheets annexed to this document (supplementary material: S1). No new fieldwork was carried out; the original contribution of this work concerns the processing, harmonization, and statistical and ecological analysis of these data.

### 2.1. Plant material

The plant material consisted of all spontaneous and/or cultivated plant species found in the surveyed plots. It comprised mainly cocoa trees (both unselected planting stock and improved varieties) and associated species (megaphanerophytes, nanophanerophytes, mesophanerophytes and microphanerophytes) present in the cocoa farms (spontaneous, remnant, and planted).

### 2.2. Study area and original sampling design

The source study covers 16 departments representative of the three major historical loops of Ivorian cocoa farming (Fig. 1; Table 1): the first loop (East/South-East; departments of Abengourou, Adzopé, Akoupé, Agboville; 116 farms covering an area of 461.15 ha); the second loop (Centre/Centre-West; departments of Djékanou, Toumodi, Daloa, Vavoua, Issia, Zoukougbeu, Yamoussoukro, Bouaflé; 178 farms with an area of 504.75 ha); and the third loop (South/South-West; departments of Divo, Guitry, Soubré, Tiassalé; 180 farms covering an area of 630.82 ha). In total, 474 farms spread across 103 villages and hamlets were surveyed (Table 1), for a cumulative area of 1596.7 ha (see supplementary material: S16 and Adji et al. (2016) for details of the sampling design).

**Fig. 1.**
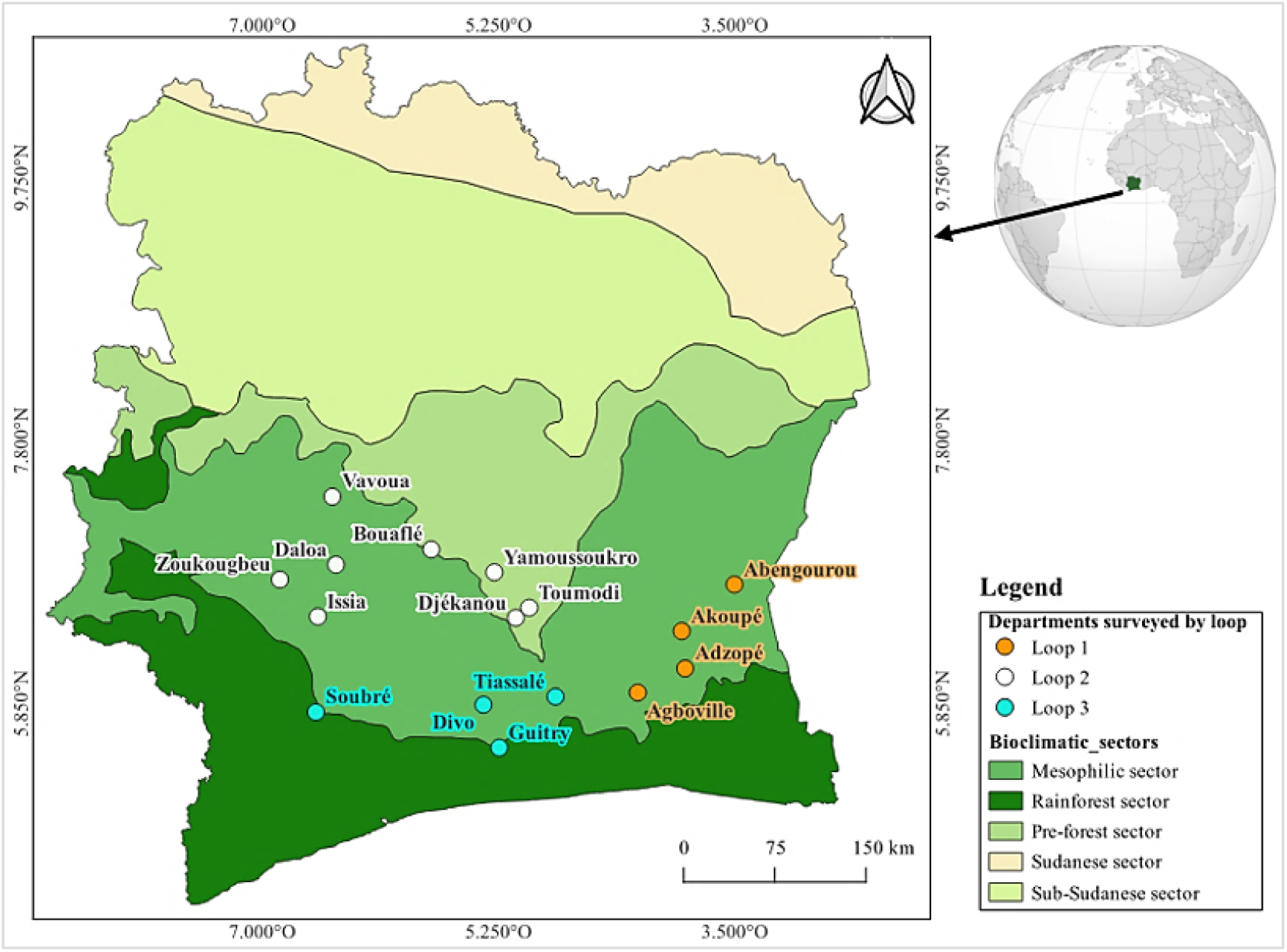
Geographical distribution map of the surveyed departments. (Source: the authors)

**Table 1.**
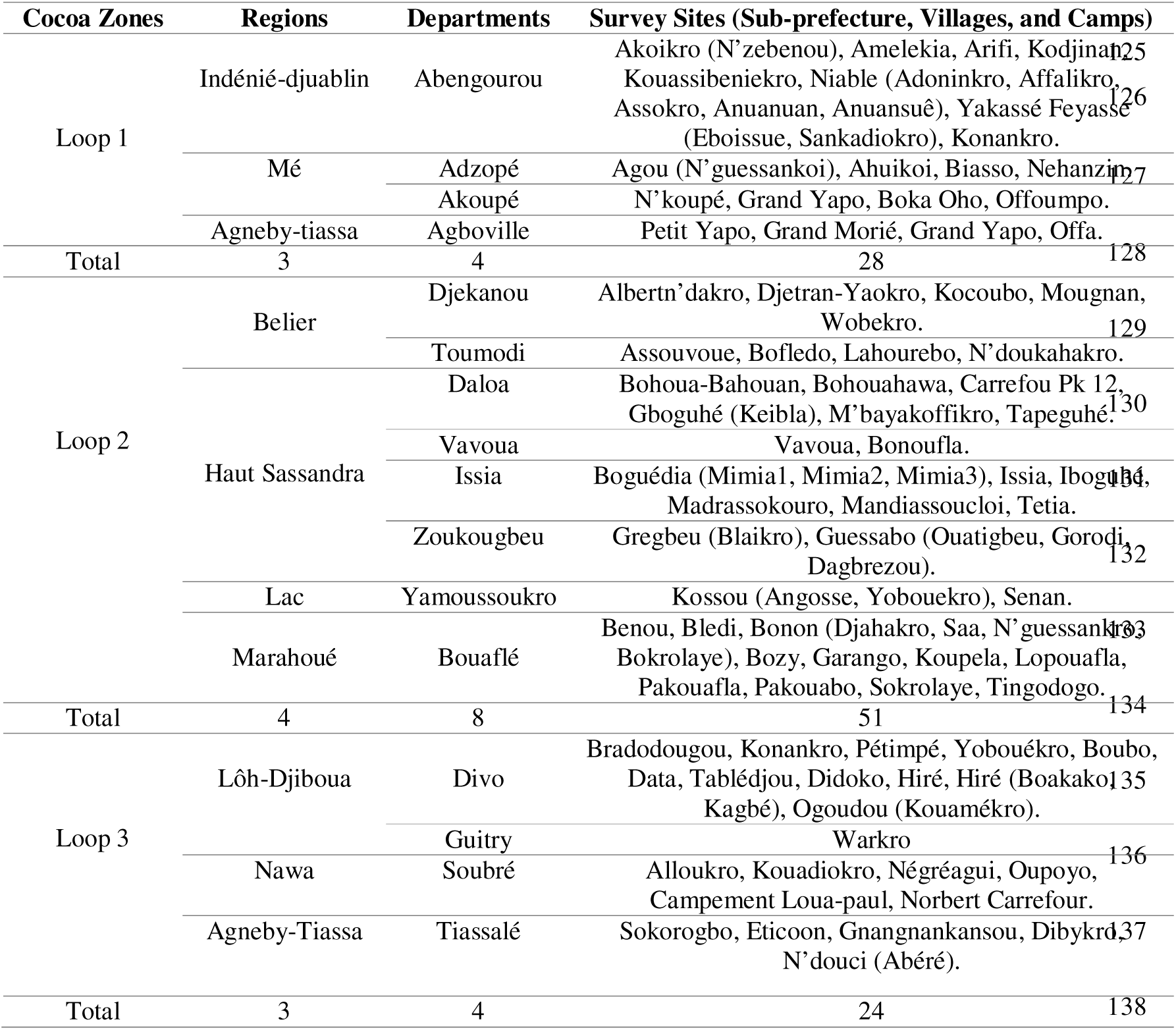
Presentation of survey sites by hamlet, village, sub-prefecture, department, region, and loop.

### 2.3. Data compilation, cleaning, and harmonization

Data were extracted from four source spreadsheets (supplementary material 1): (i) a socio-economic and agronomic survey sheet per farmer (478 raw rows, 35 coded variables); (ii) a floristic inventory of associated species by locality (3,461 raw rows: locality, vernacular name, origin, number of stems, role, part used); (iii) a master species list (277 entries, used only as a name reference); and (iv) a structural inventory per plot with vertical stratification (245 rows, 3 height strata and associated cocoa tree densities).

Programmatic cleaning (Python 3.11, pandas) normalized the spelling of place names (removing accents, harmonizing case and punctuation) and assigned each locality to one of the three loops based on the department prefixes explicit in the field labels (e.g., « ab » = Abengourou, « hs » = Haut-Sassandra, « divo » = Divo). This procedure classified 3403 of the 3404 usable floristic inventory rows (99.97%). Vernacular species names (389 raw labels, in local French and in vernacular languages « Agni, Baoulé, Dida, Bété, etc. ») were harmonized through approximate string matching (Levenshtein distance, similarity threshold 0.84), reducing the number of distinct taxa to 333. Scientific name assignment drew on two complementary sources: an identification carried out with experts from Jean Lorougnon Guédé University and the National Center for Floristics (Université Félix Houphouët-Boigny), which reliably covers the dominant and best-documented taxa; and a complementary identification sheet (183 entries, 146 with a scientific name), drawn from the associated-species census annex of the source thesis (Adji et al., 2016) and included in supplementary material (S20) of this document. These two sources converge very closely for the twenty dominant taxa, all already scientifically named. The added value of this second source concerns 26 additional, rarer vernacular taxa (outside the top 20), now scientifically identified. The remaining vernacular entries are kept as recorded and treated as folk taxa in the sense of quantitative ethnobotany, for lack of a verifiable match in either source. The taxonomic richness reported in this document should therefore be understood as a richness in vernacular taxa recognized by farmers, and not as a strict floristic richness in the Linnean taxonomic sense.

One important limitation should be noted: the sub-prefecture of Soubré (announced in the original sampling design of the third loop) does not appear in any recognizable form in the digitized floristic inventory files made available to us. Our results for the third loop therefore rest on the localities actually found in the data (Divo, Guitry, Oumé, Tiassalé). The « plantation age » field of the socio-economic survey sheet, meanwhile, is numerically recorded for 452 of the 470 farmers surveyed across the three loops (96%), which allows the agromorphological analysis by orchard age to be conducted on a sample representative of the three production « loop » zones.

### 2.4. Diversity indices and rarefaction

For each locality, a *site × taxon* matrix was built from the cumulative abundances (number of stems) recorded in the floristic inventory sheet. For each site *i*, we calculated: the observed taxonomic richness *S* (*Hill* number of order *q* = 0, *^0^D*); the *Shannon* diversity index *H’* = *-*Σ *p_k_ .ln(p_k_)* and its transformation into the order-1 *Hill* number (*^1^D* = *exp(H’)*, the « effective number of common species »); the *Simpson* concentration index and its transformation into the order-2 *Hill* number (*^2^D* = *1/*Σ*p_k_²*, the « effective number of dominant species ») (Hill, 1973; Jost, 2006); and *Piélou*’s evenness (*J* = *H’/ln S*). *Hill* numbers offer a unified interpretation in « effective species units », directly comparable to one another regardless of order *q*, unlike classical indices expressed in heterogeneous units (Chao et al., 2014).

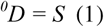

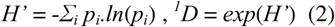

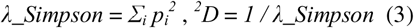

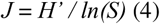

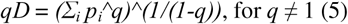

In equations (1) to (5), *S* is the number of taxa recorded at the site in question; *p_i_*is the relative abundance of taxon *i* at that site, calculated as *p_i_*= *n_i_/N* (with *n_i_* the abundance of taxon *i* and *N* the total number of individuals recorded at the site); *q* is the order of the *Hill* number, which determines the weight given to abundant taxa relative to rare ones (*q* = 0: raw richness, insensitive to abundance; *q* = 1, obtained by taking the limit of equation (5), corresponds to the exponential of the *Shannon* index *H’*; *q* = 2 corresponds to the inverse of the Simpson concentration index λ*_Simpson*); and *qD* denotes the *Hill* number of order *q*, expressed as an effective number of species (a common unit that allows the three orders to be compared directly with one another).

Because observed richness depends strongly on the number of individuals sampled (*N* ranged from 1 to 5006 individuals depending on the site), analytical individual-based rarefaction after Hurlbert (1971) was applied to estimate the expected richness at a standardized sampling effort of *n* = 15 individuals (a threshold chosen because it preserved 91 of the 99 reliable sites while limiting extrapolation). Rarefaction/accumulation curves were also calculated by zone, pooling all individuals recorded in each loop, to visualize cumulative *gamma* diversity at different sampling efforts.

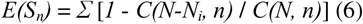

where *E(S_n_)* is the expected number of taxa in a subsample of *n* individuals drawn without replacement from the *N* individuals recorded at the site, the sum running over all *S* taxa observed at the complete site; *N_i_* is the abundance of taxon *i*, and *C(a,b)* denotes the binomial coefficient « *a* choose *b* », i.e., the number of ways of drawing *b* individuals out of *a* without regard to order. The bracketed term, equal to 1 as soon as *n* exceeds *N-N_i_*, represents the probability that taxon *i* is present in a given subsample of size *n*; the sum over all taxa thus gives the expected richness at standardized sampling effort (here *n* = 15 individuals). For example, with *N*=10 individuals in total, *N_i_*=2 of taxon *i*, and a subsample of size *n*=9 (*n* exceeds *N-N_i_*=8), we are therefore certain to draw at least one individual of taxon *i*, since there are only 8 « other » individuals. The probability of presence is therefore 1 (certain). The calculation confirms: C(8,9) = 0, so the bracket = 1-0 = 1.

Diversity was partitioned according to the multiplicative framework of Whittaker (1972) updated by Jost (2007): *gamma* (regional) diversity of each loop was defined as the total number of distinct taxa recorded across all its sites; mean *alpha* diversity as the mean of the site richnesses (*^0^D*); and *beta* diversity as the mean ratio *gamma/alpha*, interpretable as the effective number of floristically distinct communities making up the region.

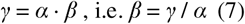

where γ is the regional *gamma* diversity of a loop (total number of distinct taxa recorded across all its sites); α is here the arithmetic mean of the site richnesses *^0^D* (equation 1) across all sites of the loop, and not the richness of a single site; and β, obtained as the simple ratio γ*/*α, is interpreted as the effective number of floristically distinct communities making up the region (Jost, 2007; Whittaker, 1972).

### 2.5. Floristic composition, ordination, and typology

The floristic composition of the 99 sites with at least 10 individuals recorded (a threshold chosen to limit sampling noise in the multivariate analyses) was compared using a Bray-Curtis dissimilarity matrix (Bray and Curtis, 1957) calculated on relative abundances (scikit-bio 0.6). A principal coordinates ordination (*PCoA*) was performed on this matrix. The effect of zone (loop) on floristic composition was tested by *PERMANOVA* (999 permutations; Anderson, 2001), and the homogeneity of within-group multivariate dispersion was checked with a *PERMDISP* test (Anderson, 2006), a condition necessary for an unambiguous interpretation of the *PERMANOVA*. A data-driven typology was then built by agglomerative hierarchical clustering (Ward’s method) applied to the first ten *PCoA* axes (68.9% of cumulative variance explained), the number of groups retained (*k* = 4) resulting from a compromise between parsimony and balanced group sizes. To avoid any ambiguity: the units grouped here are the 99 sites (farms/localities) themselves, grouped according to the similarity of their associated tree stands, and not taxa grouped into botanical categories. Each « floristic group » obtained should therefore be read as a recurring composition profile, or type, of cocoa AFS (data-driven rather than following the administrative division by loop, built directly from the taxa that co-occur and their relative abundance):

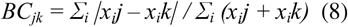

*X_i_j* and *x_i_k* are the relative abundances of taxon *i* in sites *j* and *k*, respectively; *BC_jk_* ranges from 0 (strictly identical composition) to 1 (no shared taxa), and constitutes element (*j,k*) of the dissimilarity matrix submitted to the *PCoA*, to the *PERMANOVA*, and to Ward’s classification.

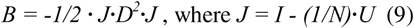

*D^2^* is the matrix of squared Bray-Curtis dissimilarities between the *N* sites; *I* is the identity matrix of dimension *N*; *U* is the square matrix of dimension *N* all of whose elements equal 1; and *J* is the resulting centering operator. The coordinates of sites on the *PCoA* are obtained by eigenvalue decomposition of the *B* matrix thus centered, with the axes corresponding to the largest eigenvalues summarizing most of the variance in floristic composition (68.9% for the first ten retained axes).

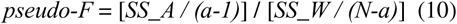

*SS_A* is the sum of squared distances (calculated from the Bray-Curtis matrix) between the centroids of the *a* compared groups and the overall centroid; *SS_W* is the sum of squared distances within each group; *N* is the total number of sites; and *a* is the number of groups compared (*a* = 3 loops). Unlike a classical *F*, the significance of *pseudo-F* is not assessed against a theoretical distribution but by permutation: zone labels are randomly reassigned to sites (999 permutations), and the proportion of permuted *pseudo-F* values equal to or greater than the observed value gives the *p* value (Anderson, 2001).

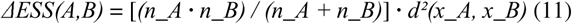

In equation (11), *A* and *B* are two candidate groups for merging at a given step of the clustering; *n_A* and *n_B* their sizes; *x_A* and *x_B* their centroids in the space of the first ten *PCoA*; and *d^2^*the squared Euclidean distance between these centroids. At each step, the Ward algorithm merges the pair of groups that minimizes the increase in Δ*ESS* in the total within-group sum of squares (error sum of squares, *ESS*), producing a hierarchy of partitions from which the *k* = 4 solution was retained.

### 2.6. Statistical models

The effect of zone on the multivariate vector of diversity indices (H’, richness S, Piélou’s evenness *J*, log total abundance *log N*) was tested using a MANOVA (*Wilks*’ test). Each index was then tested univariately by one-way analysis of variance and, given non-normal distributions, by the non-parametric *Kruskal-Wallis* test. To account for the hierarchical, spatially autocorrelated structure of the data (localities grouped into sub-zones (e.g., Haut-Sassandra, Marahoué, Bélier, Divo, Tiassalé) themselves grouped into loops), linear mixed models (random intercept by sub-zone) supplemented the fixed-effects tests. For the agromorphological analysis, yield (converted to kg/ha/year as the mean of the three annual harvest declarations recorded per farm « one per season, in bags/tonnes/kg, based on a standard weight of 75 kg/bag » divided by the declared area) was compared across the four orchard age classes (Type I: 0-5 years; Type II: 6-15 years; Type III: 16-30 years; Type IV: over 30 years, following the classification of Adji et al. (2016)) by *ANOVA/Kruskal-Wallis*, and modeled as a function of continuous age (quadratic term) by ordinary linear regression and then by a mixed model with a random intercept by village. The role of production zone (loop) in the age-yield relationship was also tested, both as a main effect and in interaction with age, to verify the consistency of this relationship across the three zones rather than within a single one. A generalized additive model (*GAM*, smoothing term on continuous age) was additionally fitted as a non-parametric check on the shape of the relationship, and 5-fold cross-validation was used to honestly assess its out-of-sample predictive power rather than its in-sample fit alone. All analyses were conducted in Python 3.11 (pandas 2, numpy, scipy 1.14, statsmodels 0.14, scikit-bio 0.6, scikit-learn, pygam). The significance threshold used is α = 0.05:

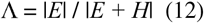

*H* is the sum-of-squares-and-cross-products matrix (*SCP*) attributable to the tested effect (production zone); *E* is the residual (within-group) *SCP* matrix, and |**·**| denotes the determinant of a matrix; Wilks’ lambda Λ ranges from 0 (perfect separation between groups across all four indices tested jointly) to 1 (no difference), and is converted into an approximate *F* statistic via *Rao*’s transformation (*F_8,180_*).

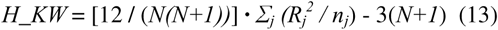

*N* is the total sample size (across all zones or all age classes, depending on the test), *n_j_* is the size of group *j*, and *R_j_* is the sum of ranks of the observations in group *j,* these ranks being computed on the complete sample rather than group by group. Under the null hypothesis of no difference between groups, *H_KW* approximately follows a χ*²* distribution with (*k-1*) degrees of freedom; with *k* the number of groups compared. This non-parametric test, insensitive to non-normality of the distributions, was used systematically to complement the classical ANOVA, both for the diversity indices and for the comparison of yield across orchard age classes.

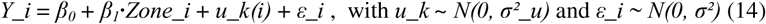

*Y_i* is the diversity index (*H’*, *S*, *J* or *log N*) observed at site *i*; *Zone_i* is the production loop to which this site belongs (fixed effect); *u_k(i)* is the random effect (intercept) of the sub-zone *k* to which site *i* belongs (accounting for the non-independence of sites within the same sub-zone rather than treating them as fully independent observations), and ε*_i* is the site-level residual. The variances σ*²_u* and σ*²* are estimated by restricted maximum likelihood (*REML*).

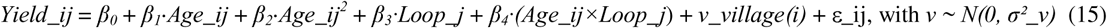

In equation (15), *Yield_ij* is the yield (kg/ha/year) of farm *i* (within its village) in loop *j*, and *Age_ij* is the corresponding orchard age (continuous variable, in years). The quadratic term Age^2^ captures a yield optimum at intermediate age (bell-shaped profile) rather than imposing a strictly monotonic relationship; *Loop_j* is a three-level categorical fixed effect, and its interaction with age tests whether the shape of this relationship differs significantly among production zones; *v_village(i)* is the random effect (intercept) of the village to which farm *i* belongs, and ε*_ij* is the farm-level residual.

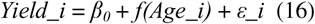

*f* is a non-parametric smoothing function (penalized regression spline) estimated directly from the data rather than constrained a priori to an imposed polynomial form. This generalized additive model (*GAM*) thus provides an independent, non-parametric check on the bell-shaped form identified by the quadratic model in equation (15).

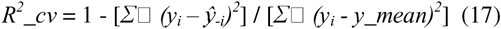

In equation (17), *y_i_*is the observed yield value for farm *i*; ŷ*_-i_* is the predicted value for that same farm from a model fitted on the other four cross-validation folds (i.e., without using observation *i* or any other observation from its fold during fitting), and *y_moy* is the overall mean yield across the full sample. Unlike the classical *R^2^*(measured in-sample), *R^2^_cv* can take negative values when the model predicts out-of-sample worse than the simple overall mean (which is precisely what is observed for the age-yield relationship, despite its in-sample statistical significance).

## 3. Results

### 3.1. Sample overview

After cleaning and harmonization, the usable floristic inventory covers 119 sites (localities), for 25,090 individuals recorded belonging to 333 distinct vernacular taxa. Of these, 99 sites met the minimum statistical reliability threshold (≥10 individuals recorded) and were retained for the comparative and multivariate analyses: 3 sites in loop 1 (department-level resolution, due to the structure of the data), 69 in loop 2, and 27 in loop 3 (Table 2).

**Table 2.** Sampling effort and diversity by production zone (« cocoa loop »)

| Zone (loop) | Sites analyzed (n) | Individuals recorded (N) | Cumulative richness $\gamma$ (taxa) | Mean richness $\alpha$ ($^0D$) | Diversity $\beta$ ($\gamma/\alpha$) |
| --- | --- | --- | --- | --- | --- |
| 1 « East/Southeast » | 3 | 6742 | 142 | 64.0 | 2.22 |
| 2 « Centre/Centre-West » | 69 out of 88 total | 8821 | 169 | 10.3 | 16.40 |
| 3 « South/Southwest » | 27 out of 28 total | 9528 | 131 | 18.3 | 7.17 |

Loop 1 is represented by only 3 units (department-level resolution), which strongly limits the precision of the $\beta$ estimate for this zone; the $N$ and $\gamma$ columns cover all available sites (before the reliability filter).

### 3.2. Alpha diversity by production zone

Alpha diversity differs strongly and significantly among the three zones (*MANOVA* on the vector [*H’*, *S*, *J*, *log N*]: *Wilks* λ = 0.350; *F_8,180_*= 15.54; *p* < 0.0001; Fig. 2). Taken individually, each univariate component confirms this overall effect: the *Shannon* index (ANOVA *F_2,96_* = 9.51; *p* < 0.001; *Kruskal-Wallis H* = 15.46, *p* < 0.001), taxonomic richness (ANOVA *F_2,96_* = 54.93; *p* < 0.0001), and the logarithm of total abundance (*F_2,96_* = 13.68; *p* < 0.0001) differed significantly by zone; only evenness (the complementary *Simpson* concentration index, *F_2,96_*= 2.76; *p* = 0.069) did not differ markedly. On average, loop 1 sites show the highest diversity (*H’* = 2.93 ± 0.19; *^0^D* = 64.0 taxa), followed by loop 3 (*H’* = 1.98 ± 0.52; *^0^D* = 18.3 taxa) and then loop 2 (*H’* = 1.56 ± 0.71; *^0^D* = 10.3 taxa). The linear mixed model with a random intercept by sub-zone confirms this effect for the *Shannon* index (loop 2: coefficient = -1.34 ± 0.40, *p* = 0.001; loop 3: coefficient = -0.94 ± 0.41, *p* = 0.021, relative to loop 1), with a very low residual inter-sub-zone variance (0.012), indicating that most of the variation in diversity is captured by the zone effect rather than by unmeasured within-zone heterogeneity.

**Fig. 2.**
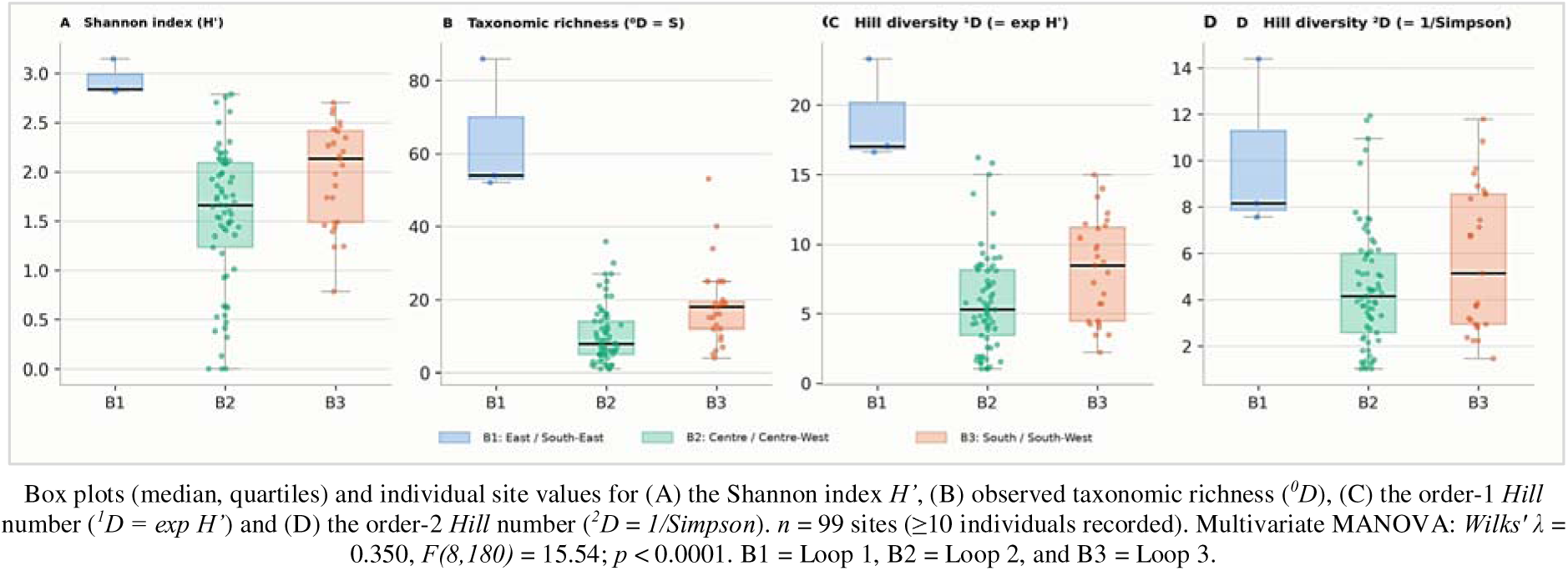
Diversity of trees associated with cocoa by production zone.

This raw result, however, may be confounded by unequal sampling effort: loop 1 sites, aggregated at the department level, accumulate on average far more recorded individuals *(N* mean = 2247) than those of loops 2 and 3, sampled at the village level (*N* mean = 100 and 340 respectively). Once richness was standardized to a common sampling effort of 15 individuals by Hurlbert rarefaction (Fig. 3), the zone effect remains significant (ANOVA *F_2,88_*= 8.72; *p* < 0.001; *Kruskal-Wallis H* = 13.79; *p* = 0.001), with an unchanged ranking (loop 1: 9.3 ± 0.9 expected taxa; loop 3: 6.9 ± 1.7; loop 2: 5.6 ± 2.1), which supports the hypothesis of a genuine ecological difference rather than a simple sampling artifact.

**Fig. 3.**
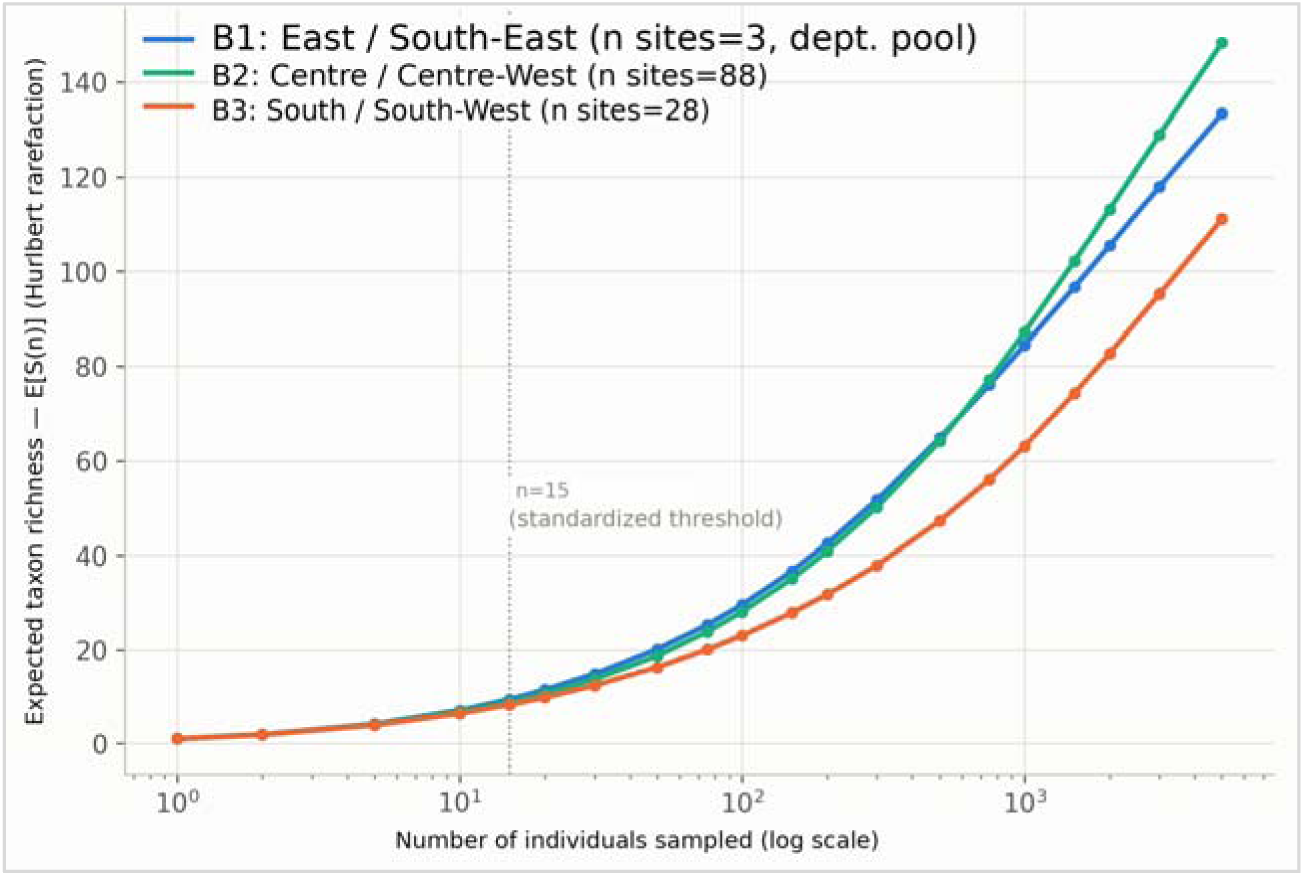
Individual-based rarefaction curves of cumulative taxonomic richness by production zone (logarithmic scale).

### 3.3. Alpha/beta/gamma partitioning of diversity

The *alpha/beta/gamma* partitioning of diversity (Table 2) reveals a major contrast invisible in a simple comparison of *alpha* indices: while loop 2 shows the lowest mean *alpha* diversity per farm, it displays the highest *beta* diversity of the three zones (β = 16.4, versus 7.2 in loop 3 and 2.2 in loop 1). This last value nonetheless remains fragile given the very small number of available units. In other words, each farm in loop 2 hosts on average few associated taxa, but the composition of these farms varies strongly from one to another, such that the cumulative regional richness (γ = 169 taxa) ultimately exceeds that of the other two zones. Cumulative rarefaction curves by zone (Fig. 3) confirm that the *gamma* richness of loop 2 catches up with and then exceeds that of loop 1 beyond about 700 cumulative individuals.

### 3.4. Floristic composition and ordination

Ordination of floristic composition by *PCoA* on Bray-Curtis dissimilarities (Fig. 4; 20.8% and 12.5% of variance explained on the first two axes) reveals a continuous gradient linking axis 1 mainly to production zone: sites in loops 1 and 3 are positioned mostly to the right of the ordination, whereas loop 2 sites spread across its entire left and central half; reflecting the greater internal floristic heterogeneity already suggested by its high *beta* diversity. The *PERMANOVA* confirms a significant effect of zone on floristic composition (*pseudo-F* = 2.85; *p* = 0.001 over 999 permutations). The *PERMDISP* test, however, indicates significant heterogeneity in within-zone multivariate dispersion (*F* = 8.07; *p* = 0.024): the significance of the *PERMANOVA* thus reflects both a shift in the floristic centroid between zones and a difference in dispersion (loop 2, being more heterogeneous, has a more scattered point cloud than loops 1 and 3). This result is consistent with the *alpha/beta* contrast described above but calls for caution in a purely “compositional” interpretation of the test used.

**Fig. 4.**
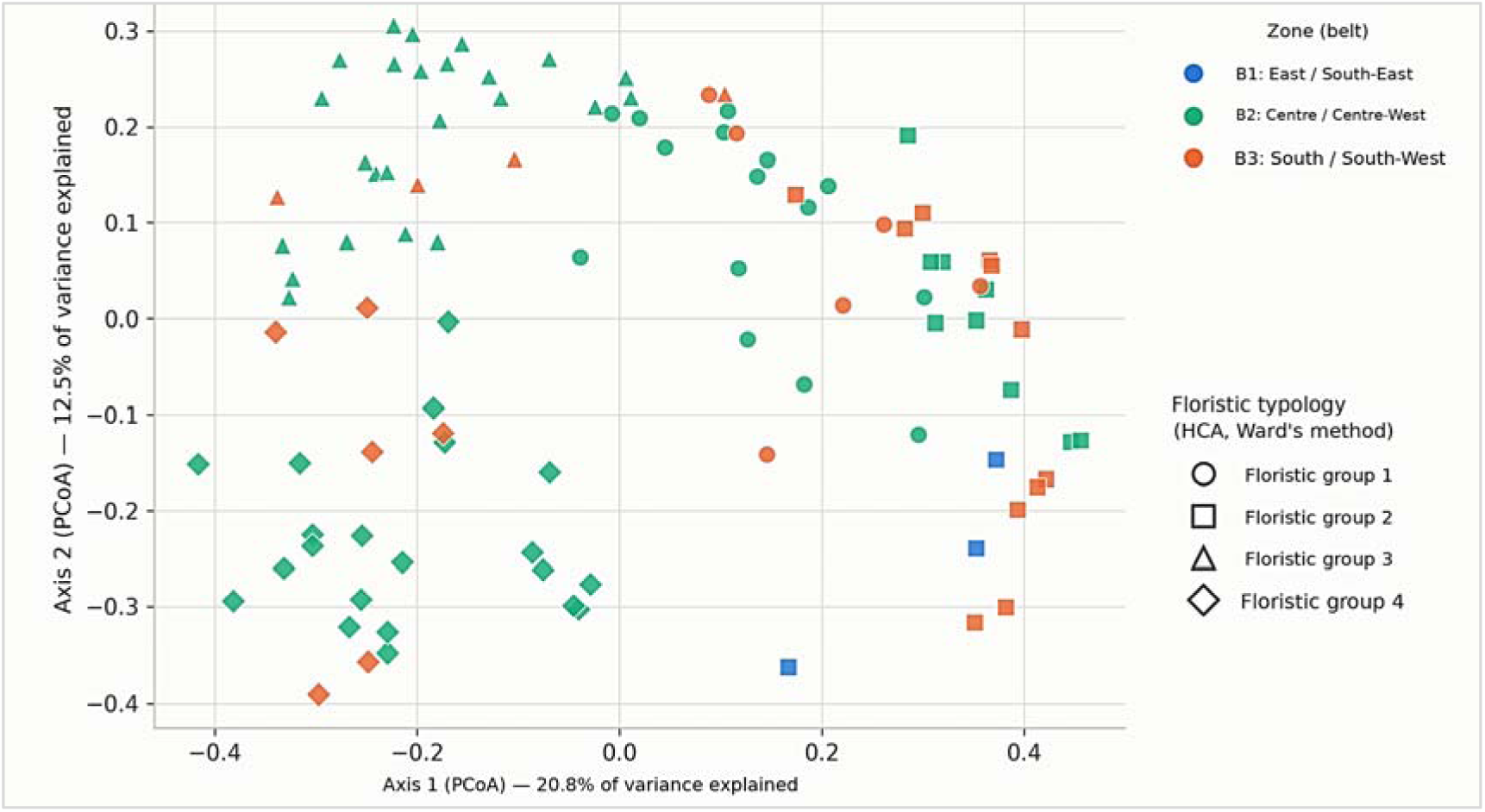
Principal coordinates ordination (*PCoA*, Bray-Curtis dissimilarity) of the floristic composition of cocoa AFS.

### 3.5. Data-driven floristic typology

Agglomerative hierarchical clustering (Ward’s method, first 10 *PCoA* axes) splits the 99 sites into four floristic groups of comparable size (21, 23, 28, and 27 sites; Fig. 5); that is, four recurring types of cocoa AFS defined by the associated taxa that co-occur and their abundance, not four groups of taxa. Their diversity profile (Table 3) traces a gradient that can be interpreted as a functional typology of cocoa AFS, independent of the simple administrative division by loop: group 2 (*n* = 23) comprises dense, diverse stands (mean abundance *N* = 740 individuals, richness *S* = 28.1 taxa, *H’* = 2.04), present in all three zones (3 loop-1 sites; 9 loop-2 sites; 11 loop-3 sites) and describable as « diversified-multi-layer » agroforestry systems; group 1 (*n* = 21) shows intermediate diversity (*S* = 17.1; *H’* = 2.11) and good evenness, dominated by loop 2 (15/21); groups 3 (*n* = 28) and 4 (*n* = 27), very largely composed of loop 2 sites (24/28 and 21/27 respectively), comprise poorer, less abundant stands (*S* = 8.3 and 5.9 respectively; *H’* = 1.58 and 1.26), corresponding to « simplified » cocoa AFS with a low density of associated trees. This data-driven typology therefore partially overlaps with the administrative zoning (loop 2 numerically dominates the poorest groups, owing to its strong representation in the sample) but above all demonstrates that contrasting agroforestry configurations coexist within a single zone, which are not captured by a typology based on orchard age alone.

**Fig. 5.**
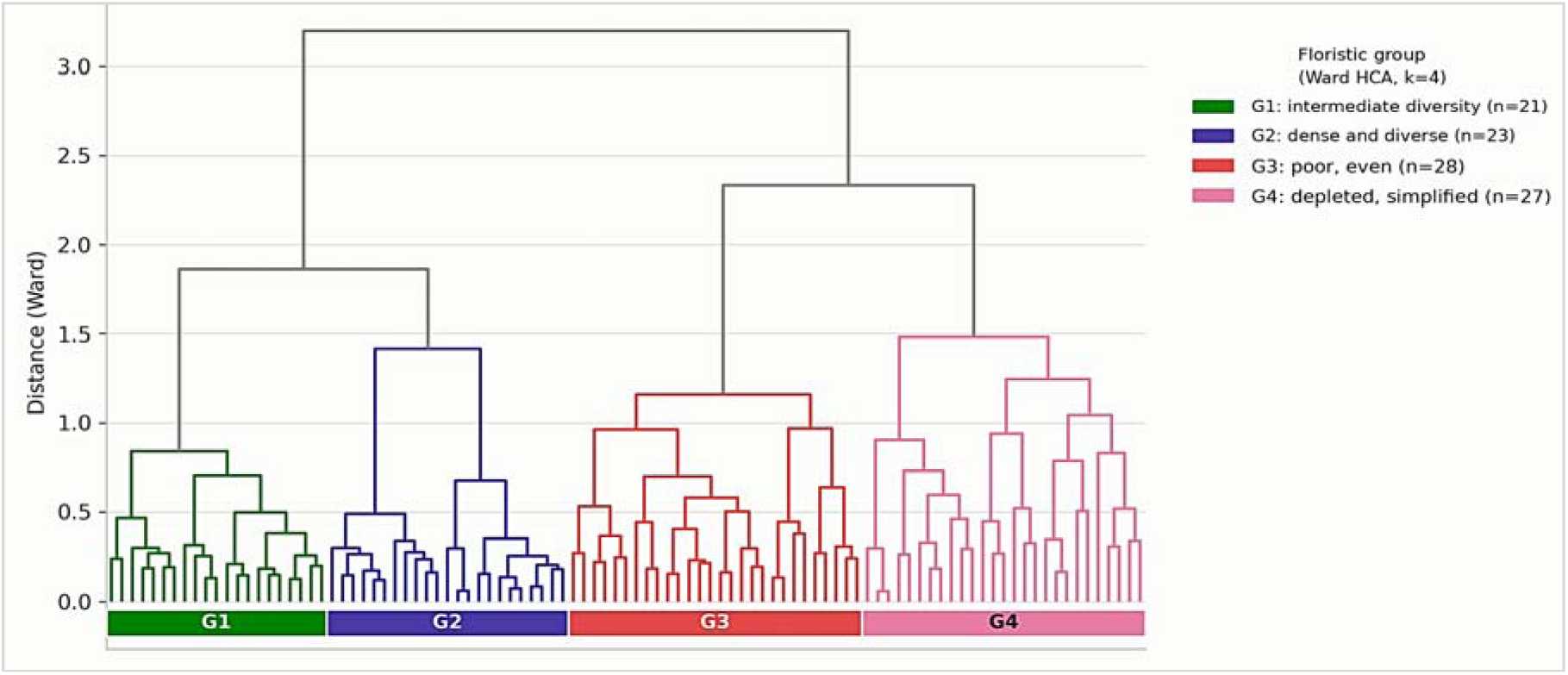
Dendrogram of the agglomerative hierarchical clustering (Ward’s method, applied to the first 10 PCoA axes, 68.9% of cumulative variance explained) of the 99 sites by floristic composition.

**Table 3.** Diversity profile of the four floristic groups (Ward’s classification, *k* = 4)

| Floristic group | *n*<br>sites | Abundance<br>*N* (mean) | Richness<br>*S*<br>(mean) | Shannon<br>*H'*<br>(mean) | Simpson<br>*1-D*<br>(mean) | Pielou<br>*J*<br>(mean) | Dominant<br>zone |
| --- | --- | --- | --- | --- | --- | --- | --- |
| 1 « diversified,<br>intermediate » | 21 | 256 | 17.1 | 2.11 | 0.81 | 0.75 | Loop 2<br>(15/21) |
| 2 « dense and<br>diversified » | 23 | 740 | 28.1 | 2.04 | 0.74 | 0.65 | Mixed (3<br>zones) |
| 3 « poor, regular » | 28 | 58 | 8.3 | 1.58 | 0.70 | 0.81 | Loop 2<br>(24/28) |
| 4 « impoverished,<br>simplified » | 27 | 36 | 5.9 | 1.26 | 0.57 | 0.76 | Loop 2<br>(21/27) |

Table 3 presents the complete diversity profile and dominant zone of each group.

### 3.6. Dominant taxa

Across the 25,090 individuals recorded, five taxa alone account for 61.9% of the stems associated with cocoa farms (Fig. 6): oil palm (*Elaeis guineensis*, 28.9%), avocado (*Persea americana*, 9.7%), mango (*Mangifera indica*, 9.6%), orange (*Citrus sinensis*, 9.2%), and kola tree (*Cola nitida*, 4.6%). This dominance imbalance is pronounced: the next 15 taxa collectively account for only 21% of the stems. Taxonomic identification of the inventory covers all twenty dominant taxa shown in Figure 6, all scientifically named, as well as 26 rarer vernacular taxa outside this top 20, out of the 389 distinct vernacular taxa recorded in the floristic inventory (Table 4); notably ilomba (*Pycnanthus angolensis*, Myristicaceae, 25th in abundance rank), lemon (*Citrus limon*, Rutaceae, 27th rank), mahogany (*Khaya grandifoliola*, Meliaceae, 41st rank), soursop (*Annona muricata*, Annonaceae, 43rd rank), rubber tree (*Hevea brasiliensis*, Euphorbiaceae, 50th rank), coffee (*Coffea canephora*, Rubiaceae, 66th rank), cola grandis (*Cola grandis*, Malvaceae, 68th rank), and sipo (*Entandrophragma utile*, Meliaceae, 83rd rank). These 26 taxa individually remain marginal in abundance (none exceeds 112 recorded stems, versus 7249 for oil palm), but their identification extends the verified taxonomic coverage of this inventory beyond the dominant core alone.

**Fig. 6.**
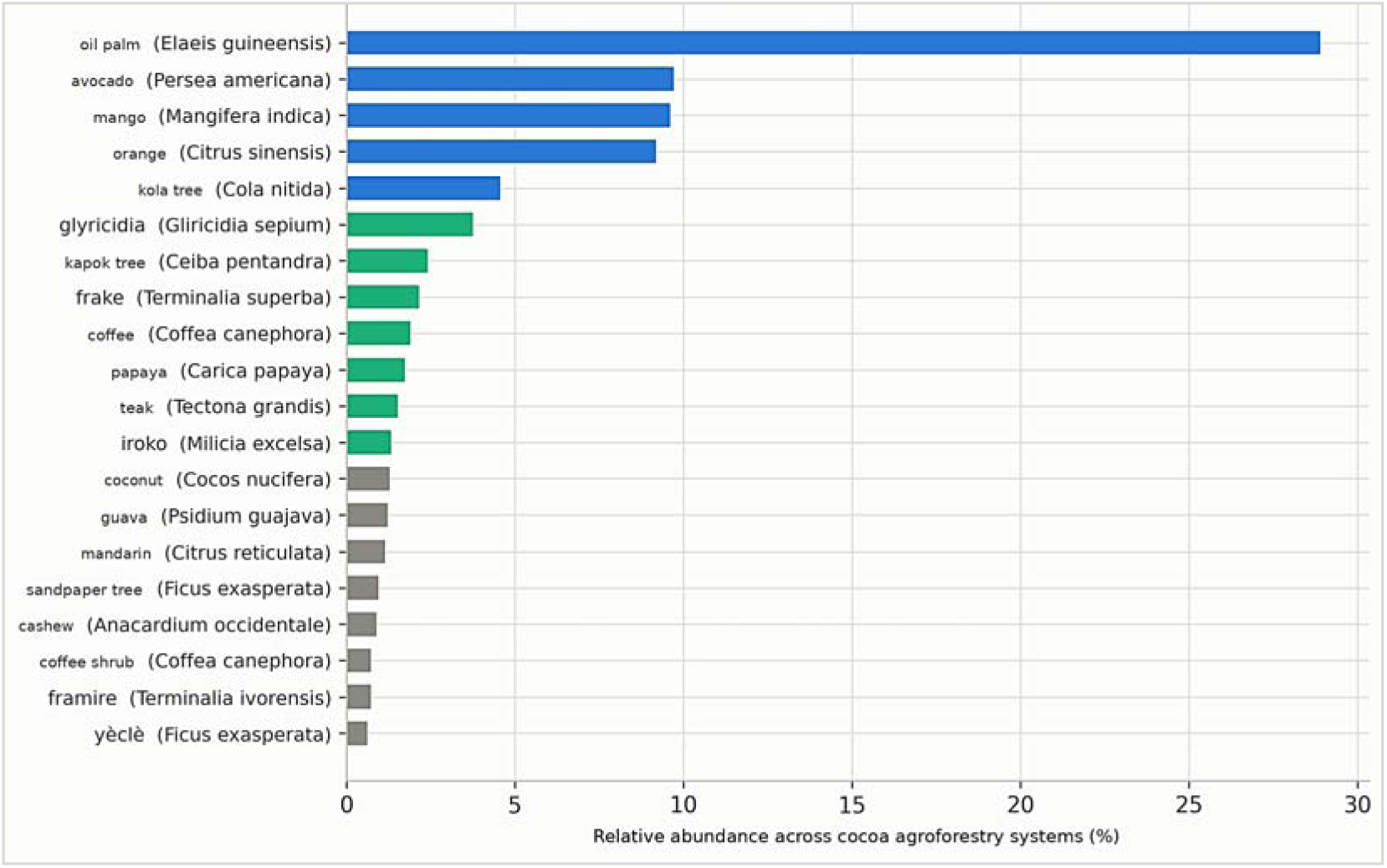
Dominant associated taxa across all the cocoa agroforestry systems studied (20 most abundant taxa, with scientific name correspondence for well-documented taxa).

**Table 4.** Vernacular name / scientific name correspondence for a subset of rare taxa newly identified through the complementary identification sheet, sorted by decreasing abundance rank.

| Vernacular name | Abundance rank | Scientific name | Family |
| --- | --- | --- | --- |
| **Ilomba** | 25 | *Pycnanthus angolensis* | Myristicaceae |
| **Lemon** | 27 | *Citrus limon* | Rutaceae |
| **Mahogany** | 41 | *Khaya grandifoliola* | Meliaceae |
| **Soursop** | 43 | *Annona muricata* | Annonaceae |
| **Rubber tree** | 50 | *Hevea brasiliensis* | Euphorbiaceae |
| **Coffee** | 66 | *Coffea canephora* | Rubiaceae |
| **Cola grandis** | 68 | *Cola grandis* | Malvaceae |
| **Sipo** | 83 | *Entandrophragma utile* | Meliaceae |

### 3.7. Vertical stratification (illustrative subsample)

The structural inventory by vertical stratum, available for an illustrative subsample of 16 individual plots, quantitatively confirms the multi-layer organization: on average, 60% of associated stems occupy the upper stratum (forest emergents, SD 18%), 23% the middle stratum (cocoa canopy height), and only 16% the lower stratum (Fig. 7). Of the 16 plots, 14 show a dominant or co-dominant upper stratum; only one is dominated by the middle stratum (plot P1, 84%, a particular orchard mainly combining intermediate-height species). The mean density of associated trees in this subsample is 33.6 stems/ha (SD 22.8), with a mean richness of 14.9 taxa per plot (SD 4.8).

**Fig. 7.**
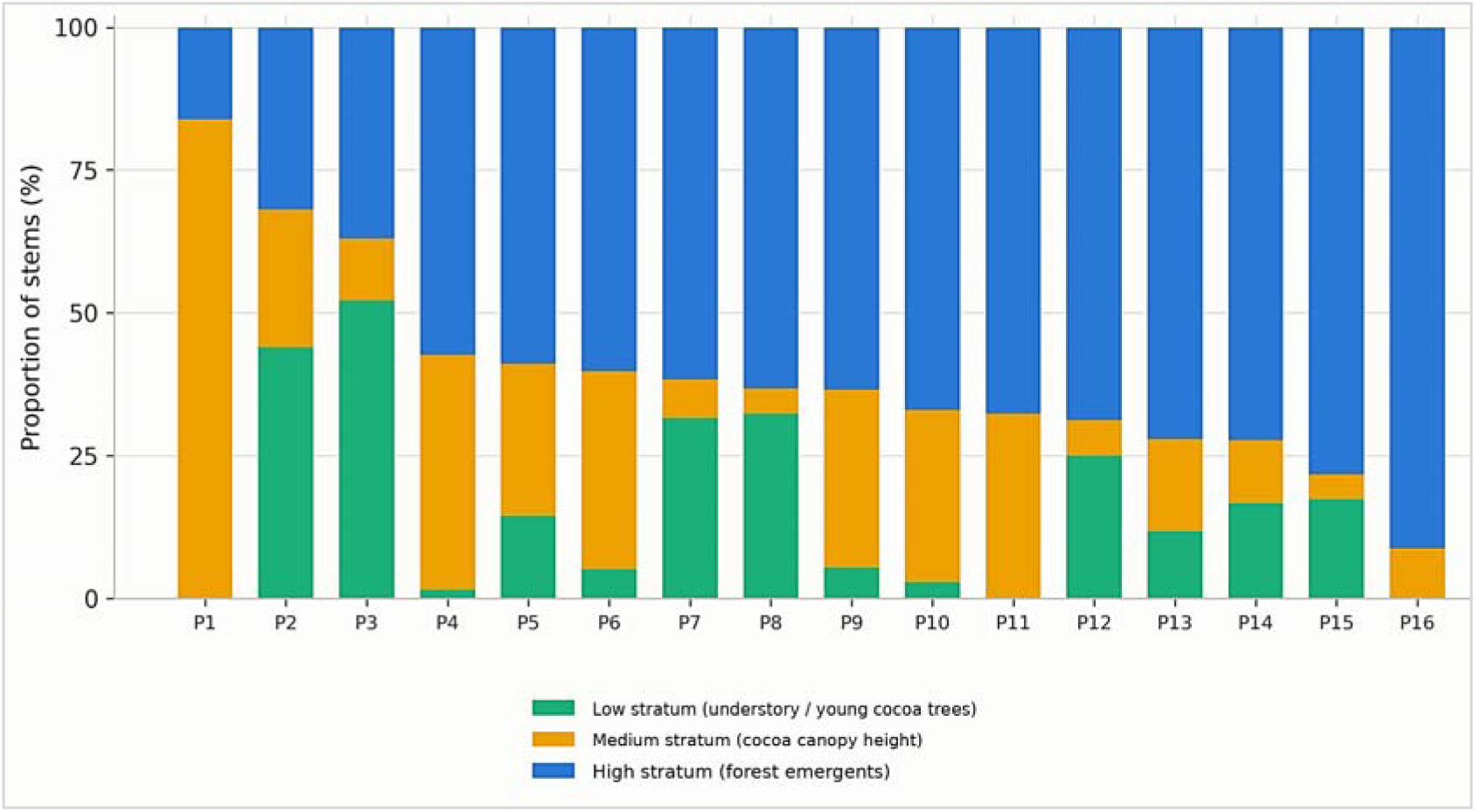
Vertical stratification of trees associated with cocoa on an illustrative subsample of 16 plots inventoried by height stratum.

### 3.8. Agromorphological typology by orchard age

The national agronomic sample (*n* = 409 farms with both age and individual yield recorded, spread across the three loops: 98 in loop 1, 151 in loop 2, 160 in loop 3; 124 villages) allows the relationship between orchard age and cocoa yield to be tested at the scale of the three production zones. Mean farm age is 24.2 years (SD 15.8; range 1-83 years), for a mean area of 3.6 ha and a mean yield of 307 kg/ha/year (median 225 kg/ha/year), calculated as the seasonal mean of the three annual harvest declarations recorded for each farm (Fig. 8). In the sub-sample of the first loop alone, the relationship is not detectable (*Kruskal-Wallis*, *H* = 0.31; *p* = 0.957; *n* = 98); at the national sample scale, however, a significant relationship emerges (*ANOVA* one-way by age class, *F_3,405_* = 3.81; *p* = 0.010; *Kruskal-Wallis*, *H* = 18.23; *p* < 0.001), following a bell-shaped profile: mean yield rises from 133 kg/ha/year for Type I orchards (0-5 years, *n* = 19) to 301 kg/ha/year for Type II (6-15 years, *n* = 145) and reaches its maximum at 354 kg/ha/year for Type III (16-30 years, *n* = 115), then falls back to 296 kg/ha/year for Type IV (over 30 years, *n* = 130). A quadratic regression on continuous age confirms this result (*R^2^* = 0.014; age coefficient: *p* = 0.021; age^2^ coefficient: *p* = 0.018; parabola peak around age 31.5), robust to log-transformation of yield (*p* = 0.005 and *p* = 0.015 respectively; peak at 37.1 years) and to the exclusion of extreme yield values (2.5th and 97.5th percentiles; *p* = 0.002 for both coefficients on the truncated sample, *n* = 387). A non-parametric generalized additive model (*GAM*) confirms the same bell shape, with a peak between 20 and 30 years, at the heart of the observed age distribution (11-35 years for the 25th-75th percentiles). The relationship remains significant after adding a random intercept by village (mixed model: age coefficient *p* = 0.042; age^2^ coefficient *p* = 0.042) and after adding loop as a covariate (*p* = 0.018 and *p* = 0.020). The *age×loop* interaction is not significant (type II ANOVA, *p* = 0.265). The categorical age-class test taken separately by loop reaches significance only in loops 2 and 3 (*Kruskal-Wallis H* = 11.11; *p* = 0.011 and *H* = 12.43; *p* = 0.006, respectively) and remains non-significant in loop 1 alone (*H* = 0.31; *p* = 0.957), whose age distribution is narrower and younger (range 1-53 years versus 1-66 and 1-83 years for loops 2 and 3). This relationship explains about 1.4 to 1.6% of the variance in yield, and 5-fold cross-validation shows no out-of-sample predictive gain over a null model (*R^2^* cross-validated close to zero, including negative in some folds).

**Fig. 8.**
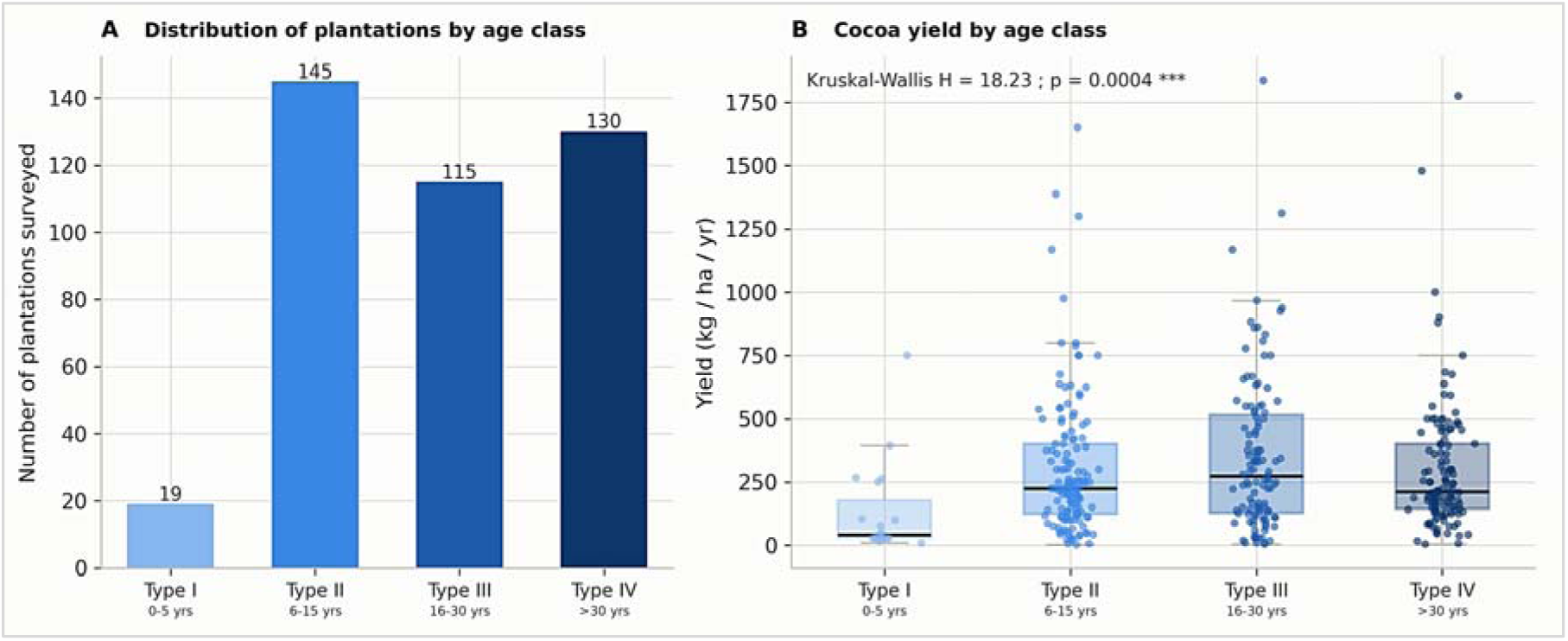
Agromorphological typology of cocoa AFS by orchard age (national sample with age and yield recorded, *n* = 409, spread across the three loops: 98 in loop 1, 151 in loop 2, 160 in loop 3).

### 3.9. Structure and diversity of associated trees by age class of cocoa orchards and loop

A complementary analysis, carried out separately for each of the three production loops and cross-tabulating cocoa orchard age class (four types, defined by the same thresholds as the national typology) with the agro-morphological characteristics of the system and the structure of associated trees, refines this national profile through a zone-by-zone reading (Table 5). The agrosilviculture system dominates very largely in the twelve *loop × age class* combinations (77 to 100% of the cocoa farms concerned), confirming the homogeneity of the underlying technical model across the three zones and the four age classes. The distribution of orchards across age classes, however, differs noticeably from one loop to another: loop 3 has the highest proportion of orchards over 30 years old (47.0%, versus 28.2% in loop 1 and 16.5% in loop 2); while loop 2 concentrates the highest proportion of young orchards (0-5 years, 13.6%). Mean yield by age class follows, in all three loops, a trajectory broadly consistent with the bell-shaped profile identified at the national scale above, with a maximum yield reached in the Type III class (16-30 years) in loops 2 and 3 (369.6 and 356.3 kg/ha/year respectively). Loop 1 is an exception, with a maximum in Type II (352.7 kg/ha/year); the Type III class instead showing the lowest yield of the four (274.5 kg/ha/year); this constitutes a singularity that should nonetheless be interpreted with caution given the small size of this sub-population.

**Table 5.**
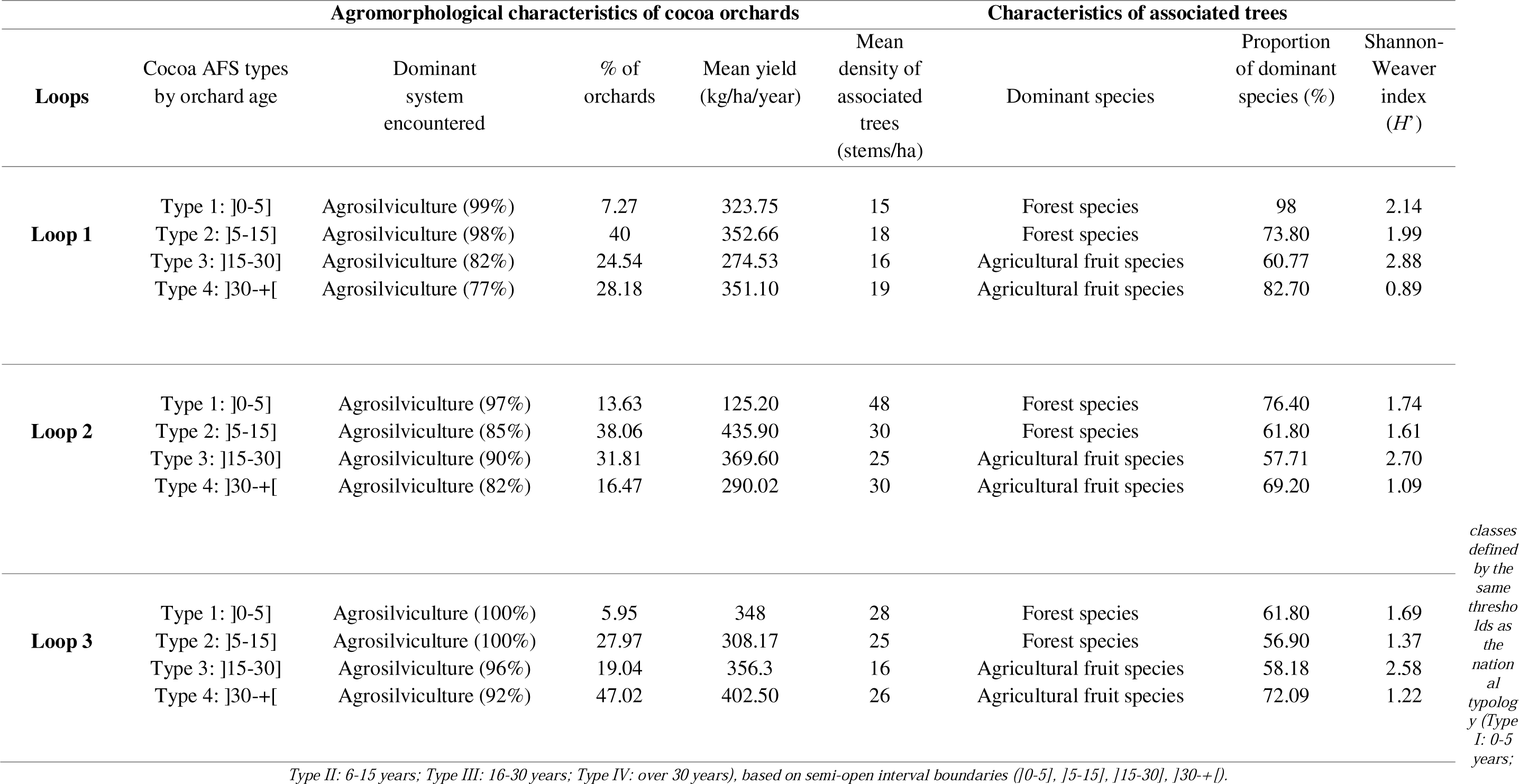
Agromorphological characteristics and structure of associated trees of cocoa agroforestry systems by orchard age class, per loop.

The composition of associated trees shifts markedly with orchard age, following a remarkably consistent pattern across the three loops: the dominant species belongs to the forest-species group in young and intermediate-age orchards (Type I and Type II, in all three loops without exception), then switches to an agricultural fruit species (avocado, mango, orange, or oil palm, depending on the zone) in mature and old orchards (Type III and Type IV, likewise without exception). This compositional shift is accompanied by a bell-shaped diversity profile, distinct from the yield profile alone: the Shannon-Weaver index rises from the Type I class to the Type III class in all three loops (for example, from 2.14 to 2.88 in loop 1; from 1.74 to 2.70 in loop 2; from 1.69 to 2.58 in loop 3), before falling sharply in the Type IV class (0.89 in loop 1; 1.09 in loop 2; 1.22 in loop 3). This drop coincides with a sharp increase in the proportion of stems belonging to the single dominant species (from 58-61% in Type III to 70-83% in Type IV, depending on the loop). The mean density of associated trees, meanwhile, generally increases with orchard age in each of the three loops (for example, from 15 to 19 stems/ha in loop 1), indicating that the decline in diversity at the end of the cycle results from an increasing concentration on a reduced number of species rather than a simple impoverishment in the abundance of the associated cover.

## 4. Discussion

### 4.1. A real regional differentiation, but more complex than a simple diversity gradient

The significance of the multivariate MANOVA on alpha diversity indices (Wilks’ λ = 0.350, *F_8,180_* = 15.54. *p* < 0.0001) is confirmed by three of the four univariate tests that compose it (the *Shannon* « *F* = 9.51; *p* < 0.001 » index, taxonomic richness « *F* = 54.93; *p* < 0.0001 », and the logarithm of total abundance « *F* = 13.68; *p* < 0.0001 »), as well as by the comparison of richness standardized by rarefaction (*F* = 8.72; *p* < 0.001), which rules out a simple sampling-effort artifact. Only evenness does not differ clearly between zones (*F* = 2.76; *p* = 0.069). Floristic composition itself also differs by zone, as shown by the *PERMANOVA* on Bray-Curtis dissimilarities (*pseudo-F* = 2.85; *p* = 0.001), but this result should be read with caution: the *PERMDISP* test indicates significant heterogeneity in within-group dispersion (*p* = 0.024), such that the significance of the *PERMANOVA* reflects both a shift in the floristic centroid between zones and a difference in dispersion, rather than an unambiguous compositional separation.

Taken together, these results converge on a robust conclusion: the three Ivorian cocoa loops host statistically distinct cocoa AFS in terms of diversity, a result that the original analysis at *n* = 3 (one observation per loop) could neither establish nor rule out with any inferential guarantee. This reanalysis thus provides a statistically grounded demonstration, at an inferentially valid scale, of a zone effect on cocoa agroforestry diversity in Côte d’Ivoire; an effect qualitatively anticipated but never formally tested at this scale in the source study, and which complements recent typological work conducted at the national scale (Adji et al., 2020; Konan et al., 2023) with explicit statistical inference based on replicated localities.

### 4.2. A regional differentiation of alpha diversity, robust to sampling effort

The alpha diversity contrast between the three loops (loop 1 > loop 3 > loop 2, confirmed by the multivariate MANOVA, *Wilks* λ = 0.350; *p* < 0.0001) could a priori be a simple artifact of the original sampling design, loop 1 being aggregated at the department scale with a markedly higher collection effort. The fact that this ranking remains unchanged after standardization by Hurlbert rarefaction at equal effort (15 individuals) rules out this explanation and confirms that this is a genuine ecological difference rather than a sampling-design artifact. This represents a methodological check rarely performed in existing West African agroforestry characterizations, which most often compare raw indices without correcting for effort (Boffa, 1999; Somarriba, 2007; Sonwa et al., 2003). This robustness reinforces the validity of the identified regional contrast, while also serving as a reminder that high richness at the department scale (loop 1) should not be interpreted at the same analytical level as richness measured at the village scale (loops 2 and 3) without this statistical precaution.

### 4.3. High beta diversity in the Centre/Centre-West zone: signature of a heterogeneous agricultural mosaic

The most original result of this reanalysis is undoubtedly the demonstration of markedly higher beta diversity in loop 2 (Centre/Centre-West, β = 16.4; versus 7.2 in loop 3 and 2.2 in loop 1), even though its mean alpha diversity per farm is the lowest of the three zones. It constitutes one of the most informative results of this new analysis, and could not have been detected from the aggregated indices of the source study alone. It reveals that the relative poverty of each individual farm in this zone coexists with very strong compositional heterogeneity from one farm to another, such that the cumulative regional richness (γ = 169 taxa) ultimately exceeds that of the other two loops. This configuration, well documented within the multiplicative diversity framework (Jost, 2007; Whittaker, 1972), is consistent with the ordination of floristic composition (Fig. 4), where loop 2 sites occupy the entire left and central half of the main plane, versus a more compact clustering of loops 1 and 3. This contrast can be interpreted in light of the region’s socio-historical context.

This mosaic is interpreted therefore as the result of uncoordinated individual agroforestry choices rather than as a homogeneous regional system. The socio-historical context of the region indeed offers a plausible line of explanation: the Ivorian Centre-West was colonized by cocoa farming later than the other loops (East/Southeast and South/Southwest loops), through particularly diverse (Dian Boni, 1978; Ruf, 1995) and successive migratory waves, characteristic of Ivorian cocoa « pioneer fronts », generating a mosaic of heterogeneous individual agroforestry practices rather than a homogeneous regional model (Ruf, 1995); which may have favored a lasting heterogeneity of individual agroforestry practices rather than convergence toward a single regional model. This link between farmer trajectories and the composition of associated woody flora nonetheless remains to be tested explicitly, for example by coupling beta diversity with individual socio-demographic covariates (farmer origin, length of settlement) in future work. This link between producer origin and the composition of the woody flora of cocoa farms has already been documented in this same region by Konan et al. (2011), who showed significant floristic differentiation of farms according to the farmer’s ethnic group and orchard age in the Ivorian Centre-West; a result consistent with the high beta diversity demonstrated here. High beta diversity can thus be interpreted as an indirect indicator of the heterogeneity of farmer trajectories rather than as a simple sampling artifact; a hypothesis that would merit explicit testing in future work coupling beta diversity with individual socio-demographic covariates. This finding also aligns with the broader tropical agroforestry literature, which shows that farm-to-farm variation in shade management is itself a major determinant of the biodiversity actually delivered by a cocoa landscape (Clough et al., 2011). An assessment of cocoa AFS « quality » at the landscape scale relying solely on a single regional average would thus have missed the most striking result of this reanalysis.

From a conservation standpoint, this heterogeneity calls for not reducing the assessment of the ecological value of cocoa AFS to the regional mean alpha diversity alone: a zone with low mean diversity per farm can nonetheless contribute significantly to regional diversity if compositional turnover between plots is high. A result that any landscape assessment based on a single indicator would miss, and which is directly relevant to the design of landscape-level rather than strictly plot-level monitoring indicators (Clough et al., 2011; Tscharntke et al., 2011).

### 4.4. Floristic composition and typology grounded in data: beyond simple administrative zoning

The *PERMANOVA* confirms a significant effect of zone on floristic composition, but the *PERMDISP* test reveals unequal within-group multivariate dispersion: the significance of the overall test therefore reflects both a genuine shift in the floristic centroid between zones and a difference in dispersion; loop 2 is intrinsically more heterogeneous than loops 1 and 3. This dual reading, rarely made explicit in West African agroforestry studies that use *PERMANOVA* without checking this condition (Anderson, 2006), calls for a cautious interpretation rather than simply concluding a uniform compositional differentiation between zones. The resulting data-driven typology (four floristic groups, ranging from « dense and diversified » to « impoverished, simplified »; Table 3) partially, but not entirely, overlaps with the administrative division by loop. Group 2 (« dense and diversified », *n* = 23) brings together sites from all three zones at once, demonstrating that a rich, structured agroforestry system is not the exclusive preserve of any one zone, but remains attainable wherever individual practices allow it. Conversely, groups 3 and 4 (« poor, regular » and « impoverished, simplified »), although very largely composed of loop 2 sites, are not entirely confined to it. These results demonstrate that contrasting agroforestry configurations coexist within a single zone, not captured by a typology based on orchard age alone.

This data-driven typology aligns with the shade-management continuum, ranging from highly diversified to highly simplified systems, widely documented in the international literature on cocoa agroforestry (Clough et al., 2011; Rolando et al., 2014; Somarriba, 2007; Tscharntke et al., 2011), and suggests that a characterization or certification policy based on administrative zoning alone would underestimate the within-zone variability that actually structures the associated tree resource. This data-driven typological approach, rather than one based on geographic description alone, fits within a broader methodological shift in the characterization of West African cocoa AFS, where recent work has also begun to distinguish associated tree stands by their origin and mode of establishment (planted, spontaneous, or remnant) rather than by geographic criteria alone (Konan et al., 2025).

### 4.5. A handful of multi-use taxa structure the entire system: an economic logic more than a spontaneous ecological one

The very strong dominance of five taxa (oil palm, avocado, mango, orange, kola tree accounting for 61.9% of recorded stems, Fig. 6) across all three zones (all with high market or subsistence use value) confirms, at an unprecedented quantitative scale, the hypothesis of a farmer income-diversification strategy already documented qualitatively in the source study (Adji et al., 2016). This result indeed confirms that the composition of trees associated with cocoa in Côte d’Ivoire results from a rational economic choice by producers much more than from simple spontaneous regeneration of forest species (Boffa, 1999; Sonwa et al., 2003). This finding aligns with the general result, drawn from syntheses on shade management in tropical cocoa- and coffee-growing regions, according to which farmer economic incentives (more than ecological optimality alone) are generally the dominant determinant of shade-species choice (Dago et al., 2025; Tscharntke et al., 2011). A recent study conducted among Ghanaian cocoa producers reaches a highly convergent conclusion: shade-species choice there is determined jointly by producers’ practical ecological knowledge and by tangible use values (food, income, pharmacopoeia), with socio-demographic factors modulating which species are actually retained on the farm (Asigbaase et al., 2025). Moreover, this result is consistent with the multiple-use definition of agroforestry (Nair, 1993) and also with recent West African work showing that farmer selection of shade species simultaneously combines income-diversification objectives with microclimatic regulation of the cocoa farm (Asigbaase et al., 2025; Konan et al., 2023; Konaté et al., 2026; Rolando et al., 2014). This is a lesson directly transposable to agroforestry extension and certification programs (deforestation-free cocoa, RA/UTZ, biodiversity cocoa), which would benefit from integrating these five taxa as default options with a high spontaneous adoption rate, rather than promoting and relying solely on native forest species valued for timber, which have historically been less well accepted by producers or whose retention by these producers has long been discouraged by a land-tenure regime granting rights over valuable trees growing in rural areas to the state rather than to the farmer (ClientEarth, 2020; Kouassi et al., 2021); a tree-tenure insecurity that the 2019 reform of the Ivorian forestry code aims precisely to correct (Côte d’Ivoire, 2019). The strong presence of oil palm (28.9% of stems, the leading associated taxon) is notable and would merit specific monitoring, as this species occupies, in Côte d’Ivoire, a position that is both complementary to and competing with cocoa depending on local land-tenure trajectories. The complementary scientific identification of 26 rarer vernacular taxa (Table 4), although marginal in individual abundance, nonetheless broadens the available knowledge base (verified taxonomic coverage of this inventory) to document the residual biodiversity actually hosted by these systems, beyond the dominant commercial core alone and recognized (mahogany: *Khaya grandifoliola*, sipo: *Entandrophragma utile*), a woody-diversification potential that remains underexploited given its low frequency.

### 4.6. Vertical stratification: a multi-layer structure consistent with the classic shaded-cocoa-based agroforestry model, subject to the caveat of an illustrative sample

The vertical stratification observed in the illustrative subsample (60% of stems in the upper stratum, dominated by forest emergents, 14 of 16 plots with a dominant or co-dominant upper stratum; Fig. 7) is consistent with the “cocoa under residual forest cover” model characteristic of West African cocoa agroforestry (Nair, 1993; Rolando et al., 2014). This clear dominance of the upper stratum is consistent with the vertical structure classically associated with the ecological and microclimatic benefits of shaded cocoa cultivation (Rolando et al., 2014; Somarriba, 2007; Tscharntke et al., 2011). This emergent-canopy organization above the cocoa trees promotes thermal and hydric regulation of the crop as well as hosting a broader associated biodiversity than the cocoa stratum alone would suggest. This result should, however, be interpreted as illustrative rather than representative at the scale of the three loops, since the available structural subsample (16 plots) is limited in size and spatial coverage relative to the complete floristic inventory; a dedicated, larger-scale structural inventory would be needed to confirm the generality of this vertical profile at the national scale. Moreover, this complex vertical architecture constitutes a structural indicator complementary to compositional diversity indices: an agroforestry system can display modest taxonomic diversity while still retaining an elaborate stratification; which has direct implications for the associated ecosystem services (wildlife habitat, aboveground carbon sequestration, microclimatic regulation). A recent synthesis based on the functional traits of shade trees highlights in this regard that, it is the traits underlying vertical structure (height, crown architecture, wood density), more than species richness alone, that determine the actual delivery of these services (Addo-Danso et al., 2024; Isaac et al., 2024). Recent work conducted on West African cocoa plots also confirms that forest-origin trees dominating this upper stratum, particularly those arising from natural regeneration rather than planting, can reach commercially exploitable diameters in as little as 14-15 years (K. Kouassi et al., 2025; Kouassi et al., 2023); an additional argument in favor of preserving this stratum, both for its ecosystem services and for its potential economic value as timber.

### 4.7. Age-yield relationship: a real but low-amplitude effect, sensitive to the scale of analysis

The analysis confirms, at the scale of the three loops combined, the existence of a significant relationship between orchard age and cocoa yield, following a bell-shaped profile peaking between 20 and 30 years (*ANOVA*, Kruskal-Wallis, quadratic regression, and *GAM* all converging; Figure 8), consistent with the qualitative suggestion of the source study and with the known biology of cocoa trees, whose productivity increases during the juvenile phase, reaches a plateau at maturity, then declines with tree senescence and the accumulation of disease pressure on older orchards. The shift from a subsample dominated by loop 1 alone to the national sample of 409 farms concretely illustrates a methodological risk inherent to any analysis restricted to a single geographic zone: taken in isolation, loop 1 shows no detectable relationship between age and yield, its age distribution being narrower and younger than that of the other two loops, whereas loops 2 and 3 each reveal a significant effect. A conclusion drawn from the loop 1 subsample alone would thus have wrongly concluded that no relationship exists, even though one is indeed real at the national scale (albeit of low amplitude).

This amplitude nonetheless remains modest: orchard age explains only about 1.4 to 1.6% of the variance in observed yield, an order of magnitude markedly lower than that generally reported for other agronomic determinants of cocoa yield (planting material, fertilization, phytosanitary management; Assiri et al., 2015, 2016). This finding converges with a very recent study on Ivorian cocoa farms, which likewise does not retain orchard age as a significant predictor of yield once a broader set of socio-demographic and biophysical determinants is accounted for (Yéo et al., 2026), suggesting that the age-yield relationship, while statistically real, remains secondary to other agronomic management levers. More decisively still, 5-fold cross-validation shows no out-of-sample predictive gain associated with this relationship over a null model: despite its robust statistical significance, it would not usefully predict the yield of an individual farm from its age alone. This dissociation between in-sample significance and out-of-sample predictive utility illustrates an important methodological point, too rarely emphasized in the descriptive agronomic literature, and underscores the value of systematically applying a dual reading (hypothesis testing and cross-validation) to any relationship identified in this type of survey data, rather than relying on the value of *p*. It suggests that orchard age, although genuinely associated with yield at the population scale, is far from being the main determinant of observed yield at the individual-farm scale, where management practices (pruning, fertilization, planting material, phytosanitary control) likely play a predominant role. This finding calls for caution in designing replanting programs based on age alone, which would benefit from being coupled with, rather than substituted for, targeted agronomic support.

### 4.8. Composition and diversity trajectory with orchard age: a dynamic consistent across loops, with the exception of loop 1

The cross-tabulated analysis by age class and loop (Table 5) reveals a remarkably consistent management trajectory across the three zones: dominance systematically shifts from a forest species in young and intermediate-age orchards (Types I-II) to an agricultural fruit species in mature and old orchards (Types III-IV), while the *Shannon-Weaver* index rises up to Type III before falling sharply in Type IV, this drop coinciding with an increasing concentration of stems on the single dominant species (from 58-61% to 70-83% depending on the loop). This diversity peak in the Type III class coincides, in all three loops without exception, with the age class in which national cocoa yield is also at its maximum, suggesting that the most productive orchards are also, structurally, the most diversified in associated species. This dynamic manifests not only as a phenomenon aggregated across farms, but also as a trajectory intrinsic to the life cycle of a single farm, occurring in two phases: a progressive enrichment of the associated cover during the orchard’s early decades, likely corresponding to increasing farmer reinvestment as the cocoa farm becomes productive, followed by an active simplification at the end of the cycle, consistent with the shade-reduction and intensification strategies documented elsewhere in the tropics as orchards age and the least productive associated trees are progressively eliminated (Clough et al., 2011; Tscharntke et al., 2011). Loop 1 is an exception to this pattern only in terms of yield (peak in Type II rather than Type III, minimum in Type III): a singularity that should be interpreted with caution given the small size of its subsample (*n* = 98) and its department-level aggregated sampling structure; a signal to be confirmed rather than a refutation of the national profile. It may also reflect this zone’s older trajectory, as the historical cradle of Ivorian cocoa farming, where some mature orchards may already be entering a renewal cycle not captured by declared age alone. This singularity illustrates, once again, the value of a zone-disaggregated analysis rather than a single national average.

### 4.9. Methodological contribution, limitations, and perspectives

From a methodological standpoint, this reanalysis illustrates the value of remobilizing, using an ecological and statistical framework now standard (Hill numbers, individual-based rarefaction, *alpha/beta/gamma* partitioning, ordination, and *PERMANOVA* on Bray-Curtis dissimilarities, data-driven typology, mixed models, and *GAM* validated by cross-validation), existing agroforestry survey data that were initially processed descriptively, without requiring new field data collection. This framework, already widespread in the international ecological literature on shaded perennial crops (Anderson, 2001; Chao et al., 2014; Clough et al., 2011; Isaac et al., 2024; Jost, 2007, 2006), remains still little used in the characterization of West African cocoa AFS, where most national inventories remain aggregated and descriptive.

This work nonetheless has limitations that should be transparently acknowledged. First, the data come from a single cross-sectional survey (2013-2016); inferring an age-related trajectory from a snapshot sampling design rests on the usual but not directly verifiable assumption that older orchards reflect the future trajectory of younger orchards rather than cohort differences linked to different past practices; a longitudinal follow-up or repeated resurveying of the same plots would help resolve this ambiguity. Second, the reported richness remains a richness in harmonized vernacular taxa recognized by producers, not a strict Linnaean floristic richness, except for the core of dominant taxa and the 26 rare taxa identified a posteriori, which benefit from complete scientific identification. Third, the heterogeneous spatial resolution (department for loop 1, village for loops 2 and 3) limits the precision of *beta* diversity estimates for the first zone; since it is represented by only three units in the diversity analyses, the statistical power and generalizability of comparisons involving this zone remain limited and should be interpreted with caution. Fourth, this study remains focused on the woody floristic component and draws on no direct faunal biodiversity data (avifauna, entomofauna); coupling it with faunal inventories, as has been done elsewhere in the tropics (Clough et al., 2011; Tscharntke et al., 2011), would constitute a natural extension for more fully assessing the conservation value of Ivorian cocoa AFS.

Finally, the agronomic sample available for the age-yield analysis is not linked to floristic diversity data at the individual-farm scale, which prevents testing whether the diversity of associated trees directly modulates yield at the plot level; a limitation that, together with the following avenue, defines the main research perspective arising from this work. A complementary approach, illustrated by very recent work conducted on Ivorian cocoa plots, would consist of distinguishing, within the floristic inventory itself, the origin of associated trees (planted, spontaneous, or relict from prior forest cover), these three cohorts contributing distinctly and complementarily to the stand’s total diversity (Konan et al., 2025); an additional resolution that could enrich the typology proposed here.

### 4.10. Implications for cocoa sustainability policy and « zero-deforestation » certification

The typology and *beta* diversity results presented here have practical resonance that extends beyond academic ecology, in light of the emerging traceability and due-diligence requirements of the European Union regulation on « deforestation-free » products (European Union, 2023), whose full application to cocoa-sector operators is scheduled for 30 December 2026 and which requires export supply chains to document, with increasing spatial granularity, the ecological status of the land from which the cocoa originates.

On the one hand, the demonstration of high *beta* diversity in loop 2, invisible in an aggregated reading based on a simple regional average, shows that credible documentation of the ecological value of cocoa landscapes cannot be limited to a single per-farm diversity indicator, but must incorporate a landscape dimension. On the other hand, a statistically grounded typology of cocoa AFS at the locality scale (as opposed to a simple descriptive average by historical loop) offers precisely the kind of spatially resolved evidence base that due-diligence mechanisms will increasingly require. The data-driven typology constitutes, in this regard, an objective, reproducible framework, less costly than an exhaustive inventory, for targeting agroforestry support interventions toward the most simplified configurations, rather than applying a uniform policy by administrative zone.

This documentary requirement, however, can only translate into genuine improvement in practices if producers themselves are able to meet it: recent empirical work shows that this degree of preparedness remains highly heterogeneous in Côte d’Ivoire and depends strongly on producers’ integration into collective support structures such as cooperatives (Moluh Njoya et al., 2025), which suggests that cocoa sustainability policies would benefit from linking documentary requirements with the strengthening of farmer support structures (by coupling organizational support with a spatially resolved ecological diagnostic such as the one proposed here) rather than treating these two levers separately. More broadly, the statistical framework used in this work, fully reproducible from already-collected survey data, could be applied at low marginal cost to other national agroforestry inventories in West and Central Africa facing similar methodological limitations, thereby helping to document, with the statistical rigor now expected by markets and regulators, the actual state of the tree component of cocoa landscapes.

## 5. Conclusion

This ecological and statistical reanalysis of survey data on cocoa-based agroforestry systems in Côte d’Ivoire indicates a marked regional differentiation in *alpha* diversity across the three production loops, but above all reveals, through the *alpha/beta/gamma* partitioning, an unprecedented contrast between lower local diversity and markedly higher *beta* diversity in the Centre/Centre-West zone; a sign of a mosaic of heterogeneous farmer practices rather than a homogeneous regional system. A data-driven floristic typology, independent of administrative divisions, distinguishes four cocoa AFS profiles and shows that within-zone variability often exceeds that observed between zones, calling for interventions to be targeted by system profile rather than by region. The relationship between orchard age and yield, confirmed at the national scale following a bell-shaped profile peaking around 20-30 years, remains, for its part, of low amplitude and without out-of-sample predictive power; a useful reminder that statistical significance and practical utility are not the same thing. This work proposes a reproducible and transferable methodological framework, still little used in the characterization of West African cocoa AFS, directly relevant to the ecological evidence requirements set by « zero-deforestation » policies; notably the European Union regulation on deforestation-free products. Its main limitations (cross-sectional data, heterogeneous spatial resolution, absence of faunal data) define the priorities for future work, particularly a longitudinal follow-up of the same plots and coupling with animal biodiversity inventories. As these regulations increasingly demand spatially resolved ecological evidence, statistically defensible typologies such as the one proposed here are poised to become a practical tool for cocoa sustainability policy, beyond their contribution to Ivorian agroforestry research.

## Supporting information

Survey questionnaire and species recorded in cocoa plots list in each loop or belt

## Author Contributions

B.I.A. Developed the methodology, collected survey data, analysed the data and wrote the paper. A.A.A. Developed the methodology, collected survey data and supervised the work. A.D.L.H. Produced the distribution map of the surveyed study sites. A.M.E., K.K.E. and D.S.A supervised the work.

## Fundind

This work was funded by the National Centre for Agricultural Research of Côte d’Ivoire (*CNRA*).

## Acknowledgements

The authors would like to thank the rural communities in the surveyed areas for their hospitality, kindness and cooperation, and for making their plantations available to ensure the smooth implementation and execution of the activities carried out on their land during this period. The authors would also like to thank ANADER Côte d’Ivoire for its support, for making the cooperatives and cocoa farmers in the surveyed areas available, and for facilitating access to them.

## Data availability

The datasets analysed as part of this study are available at: Adji, B. I., Assiri, A. A., Assi, M. E., & Kassin, K. E. (2026). Raw survey and floristic inventory data on cocoa-based agroforestry systems in Côte d’Ivoire (three cocoa-growing "loops", 2013-2016): anonymized version [Dataset]. Zenodo. https://doi.org/10.5281/zenodo.22016383

