## Supplementary material for "Tree diversity and composition of cocoa agroforests in Côte d’Ivoire: a regional ecological mosaic, implications for cocoa landscapes sustainable management": Survey questionnaire and species recorded in cocoa plots list in each loop or belt

**Suplementary materials**

#### S 1 : Survey questionnaire

Questionnaire code:/____/____/___/

Surveyor : …………………………

Date : /____/___/___/

##### I. General information

Producer's first and last name: ______________________________ Contact: ……………………………

Ethnic origin: ____________________________

Nationality: ____________________________

Age: ____________________________

Education level: ____________________________

Region: ____________________________

Department: ____________________________

Sub-prefecture: ____________________________

Village: __________________________

Camp: __________________________

Member of a farming cooperative :

☐ Yes ________________________1
Which one :……………………………..................................

☐ No ______________________2

#### II. Information on the cocoa plantation

| No. | Questions | Answers | Answer no. |
|---|---|---|---|
| 2.1 | How did you acquire your cocoa plantation ? | ☐ Created by the producer himself _____1<br>☐ Inheritance ____________________2<br>☐ Purchase _____________________3<br>☐ Rental ____________________4<br>☐ Other (specify) _________________5 | |
| 2.2 | Where is your plantation located ? | ☐ National Park and Reserves ______________1<br>☐ Classified Forest ______________________2<br>☐ Rural land__________________________3<br>☐ Other (specify) ______________________4 | |
| 2.3 | What was the previous land use ?<br>(*land use before the plantation was established; tick the correct answer*) | ☐ Forest ______________________ 1<br>☐ Old cocoa trees ______________________ 2<br>☐ Old coffee trees ______________________ 3<br>☐ Fallow of more than 10 years _____________ 4<br>☐ Fallow of less than 10 years _____________ 5<br>☐ Other (*specify*) ______________________ 6 | |
| 2.4 | Where did you obtain the cocoa seedlings or seeds used to establish your plantation | ☐ SATMACI____________1<br>☐ ANADER____________ 1<br>☐ CNRA______________1<br>☐ Other_______________2<br>NB : Satmaci, Anader and Cnra = selected plant material_1 | |
| 2.5 | What is the total area of your plantation ? | Total plot area: ......................ha | |

| | | |
|---|---|---|
| 2.6 | In what year did you establish this plantation ? | Year plot establishment began:………………<br>Year plot establishment was completed:…………………… |
| Planting density (1) ____________________ trees/ha (2) ___________________ trees/ha (3) _______________ trees/ha | | |
| 2.7 | What was the plantation's output over the last 3 seasons ? | 2009/2010 : ...........................kg or bags of ............kg<br>2010/2011 : ...........................kg or bags<br>2011/2012 : ...........................kg or bags |
| 2.8 | Does your plantation have associated trees ? | ☐ Yes _________________1<br>☐ No_________________2<br>Why :……………………………………………………… |
| 2.9 | Do you have other cocoa plantations elsewhere ? | ☐ Yes __________ 1 How many ?: ……………….<br>☐ No __________ 2 |
| 2.10 | Do all of these other plantations have associated trees ? | ☐ Yes _________________1<br>☐ No_________________2<br>Why :……………………………………………………… |

### III. Information on trees associated with cocoa

| | | |
|---|---|---|
| 3.1 | What are the origins of the trees present in your plantation ? | ☐ Residual<br>*(forest trees not felled during land preparation*<br>*of the land)* ________ 1<br>☐ Naturally regenerated<br>*(forest trees that grew naturally after the plantation*<br>*was established and that were retained*<br>*by the producer)* ________ 2<br>☐ Planted by the producer ________ 3<br>*Specify the forest trees planted* : ……………………………………………………………………………...<br>☐ Other origins *(specify) :* .......................................<br>.............................................................. _______ 4 |
| 3.2 | How are these trees arranged in the plantation ? | ☐ In rows ______________________________1<br>*(specify the spacing* <u>*in metres*</u> *between the trees)*<br>.............................................................................<br>☐ Not in rows ______________________ 2 |
| 3.3 | What type of seedlings or planting material did you use for the planted trees ? | ☐ Wildings ________________1<br>☐ Seeds _________________2<br>☐ Cuttings _________________3<br>☐ Layered plants _________________4<br>☐ Grafts _________________5<br>☐ Other (specify) _____________6 |

| | | |
|---|---|---|
| 3.4 | For the planted trees, did you produce the seedlings yourself ? | ☐ Yes<br>How : (specify techniques) :………………………………………<br>……………………………………………………………………………..<br>☐ No<br>Why :……………………………………………………………………………<br>……………………………………………………………………… |
| 3.5 | At what age of the cocoa plot did you plant the trees ?<br>(*for forest and fruit/cash trees introduced by the farmer)* | Young plot :<br>☐ ] 0-5] ____________________1<br>☐ ] 5-10] ____________________ 2<br>Why :…………………………………………………… |
| | | Adult plantation :<br>☐ ] 11-15] ______________________ 3<br>☐ ] 16-20] _______________________4<br>☐ ] 21-30] _______________________5<br>Why :…………………………………………………………………… |
| | | Senescent plantation :<br>[31-plus] ___________________________6<br>Why :…………………………………………………...……………… |
| 3.6 | Who advised you to plant or retain trees in the plantation ? | ☐ Family / Friends _______________1<br>☐ SATMACI __________________2<br>☐ CNRA __________________3<br>☐ ANADER _________________4<br>☐ Certification programme (specify) __5………………………..<br>☐ Other (specify) _________________6…............................. |
| 3.7 | Have you ever removed forest trees from your plantation ? (*over the lifetime of the cocoa plot)* | ☐ Yes _________________________1<br>When :…………………………………………………………<br>Which ones?....................................................................................<br>Why : ……………………………………………………………………<br>☐ No ________________________2 |

### IV. Tree management and ownership within cocoa AFS

| | | |
|---|---|---|
| 4.1 | Do you receive support from any organization for managing the forest trees on your plantation? | ☐ Yes _________________________1<br>Which ones?....................................................................................<br>☐ No ________________________2 |
| 4.2 | Do you think forest trees are useful in plantations ? | ☐ Yes ________________________1<br>☐ No ….______________________2 |
| 4.3 | What effects do these forest trees have on the plantation and on the cocoa trees ? | ***Benefits :***<br>☐ Reduced weed growth _____1<br>☐ Good yield_____________2<br>☐ Promotes soil moisture _______3 |

| | | |
|---|---|---|
| | | ☐ Good phytosanitary condition______________4<br>☐ Improved soil fertility__5<br>☐ Provides shade__________6<br>☐ Other (specify ) _____________7…………………… |
| | | ***Drawbacks***<br>☐ Water competition __________________________ 1<br>☐ Bring diseases and pests ____________ 2<br>☐ Lead to lower yields _______________ 3<br>☐ Cause damage by falling on the cocoa trees ________ 4<br>☐ Hinder the proper development of the cocoa tree ______ 5<br>☐ Depletes the soil __________________________ 6<br>☐ Other (specify) _____________________________7<br>………………………………………………………………………………… |
| 4.4 | Which forest trees do you prefer ? *(name the 3 most important)* | (1)……………………………………………………………………………<br>(2)……………………………………………………………………………<br>(3)…………………………………………………………………………… |
| 4.5 | Which forest trees would you like to remove from your farm ? *(name the 3 least preferred)* | (1)……………………………………………………………………………<br>(2)……………………………………………………………………………<br>(3)…………………………………………………………………………… |
| 4.6 | Do you encounter difficulties in managing the forest trees on your cocoa plantation ?<br>*(within the community or with the State)* | ☐ Yes _____________1<br>Which ones, and with whom:………………………………………………<br>…………………………………………………………………………………<br>…………………………………………………………………………………<br>…………………………………………………………………………………<br>☐ No ….._______2 |
| What solutions do you propose for the problems or difficulties you have listed?:<br>(1)……………………………………………………………………………………………………………<br>(2)……………………………………………………………………………………………………………<br>(3)…………………………………………………………………………………………………………… | | |
| 4.7 | Fill in the table below. | |

| No. | Tree names | Origin<br><br>- *Natural : 1*<br>- *Planted :..2*<br>- *Other :…3* | How many trees are there in the plantation ? | What roles does the tree play ?<br><br>- *Shade ..........1*<br>- *Fertilization------2*<br>- *Food........3*<br>- *Timber.....4*<br>- *Boundary marking.......5*<br>- *Medicinal.........6*<br>- *Belief/ritual...........7*<br>- *Other…………..8 (specify)* | Are any parts of these trees consumed or sold ? Which ones ?<br><br>- *Leaves........1*<br>- *Fruits............2*<br>- *Bark........3*<br>- *Roots........4*<br>- *Sap.............5*<br>- *Other ………6. (specify)* | What quantities of these parts are produced per year ?<br><br>*Specify bags, litres, kg, etc.* | How much does selling these parts bring in per year ?<br><br>*Give the value in FCFA* | Other information on the trees or the plantation |
|---|---|---|---|---|---|---|---|---|
| 1 | | | | | | | | |
| 2 | | | | | | | | |
| 3 | | | | | | | | |
| 4 | | | | | | | | |
| 5 | | | | | | | | |
| 6 | | | | | | | | |
| 7 | | | | | | | | |
| 8 | | | | | | | | |
| 9 | | | | | | | | |
| 10 | | | | | | | | |
| 11 | | | | | | | | |
| 12 | | | | | | | | |
| 13 | | | | | | | | |
| 14 | | | | | | | | |
| 15 | | | | | | | | |

#### V. Inventory sheet

| | |
|---|---|
| Date :___/___/___ | Farmer’s name :______________________________________ |
| Department : _____________________ | Contact :_________________________________________________ |
| Sub-prefecture : ___________________ | Farm area :________________________________ |
| Village :__________________________ | Year established :_________________________________________ |
| Camp :______________________ | |

| No. | Local name of the tree | Number of trees per stratum level | | | Observations |
|---|---|---|---|---|---|
| | | Lower stratum (below the cocoa tree canopy and >130cm) | Medium stratum (same level as the cocoa tree) | Upper stratum (clearly above the cocoa tree) | |

Cocoa tree density : Density 1 :______ trees ; Density 2 : ______ trees ; Density 3 :_________trees

#### S 2 : Residual and naturally regenerated forest species recorded in cocoa plots of Belt 1

| Scientific names | Botanical families | Number of individuals (n=6773) | Representativeness % | Number of plots (n=116) | % of plots | Shannon index (n=3,026) |
|---|---|---|---|---|---|---|
| *Elaeis guineensis* | Palmaceae | 1824 | 26,93 | 75 | 64,66 | 0,35330399 |
| *Terminalia superba* | Combretaceae | 354 | 5,15 | 57 | 48,28 | 0,15281323 |
| *Ceiba pentandra* | Bombacaceae | 196 | 2,76 | 55 | 43,97 | 0,09910726 |
| *Terminalia ivorensis* | Combretaceae | 161 | 2,38 | 41 | 35,34 | 0,08888624 |
| *Chlorophora excelsa/C, regia* | Moraceae | 135 | 1,99 | 35 | 30,17 | 0,07804257 |
| *Alstonia boonei* | Apocynaceae | 86 | 1,27 | 25 | 21,55 | 0,05544165 |
| *Entandrophragma angolense* | Meliaceae | 62 | 0,92 | 18 | 15,52 | 0,04296487 |
| *Ricinodendron heudelotii* | Euphorbiaceae | 23 | 0,34 | 13 | 11,21 | 0,01930603 |
| *Pygnanthus angolensis* | Myristicaceae | 111 | 1,64 | 13 | 11,21 | 0,06737631 |
| *Bombax buenopozense* | Bombacaceae | 58 | 0,86 | 12 | 10,34 | 0,04076404 |
| *Nesogordonia papaverifera* | Sterculiaceae | 60 | 0,89 | 12 | 10,34 | 0,04186938 |
| *Triplochiton scleroxylon* | Sterculiaceae | 63 | 0,93 | 10 | 8,62 | 0,04350902 |
| *Cola gigantea* | Sterculiaceae | 83 | 1,23 | 9 | 7,76 | 0,05394276 |
| *Musanga cecropioides* | Cecropiaceae | 45 | 0,66 | 6 | 5,17 | 0,0333134 |
| *Mansonia altissima* | Sterculiaceae | 10 | 0,15 | 5 | 4,31 | 0,00962367 |
| *Coffea canephora* | Rubiaceae | 486 | 7,18 | 5 | 4,31 | 0,1890392 |
| *Milicia regia* | Moraceae | 10 | 0,15 | 5 | 4,31 | 0,00962367 |
| *Rauvolfia vomitoria* | Apocynaceae | 70 | 1,03 | 5 | 4,31 | 0,04725444 |
| *Holarrhena floribunda* | Apocynaceae | 14 | 0,21 | 5 | 4,31 | 0,01277764 |
| *Antiaris africana* | Moraceae | 11 | 0,16 | 4 | 3,45 | 0,01043125 |
| *Ehouré(agni)* | X | 19 | 0,28 | 4 | 3,45 | 0,01648442 |
| *Margaritaria discoidea* | Euphorbiaceae | 12 | 0,18 | 4 | 3,45 | 0,01122538 |
| *Sterculia tragacantha* | Sterculiaceae | 19 | 0,28 | 4 | 3,45 | 0,01648442 |
| *Raphia hookeri* | Palmaceae | 39 | 0,58 | 4 | 3,45 | 0,02969561 |
| *Entandrophragma utile* | Meliaceae | 10 | 0,15 | 4 | 3,45 | 0,00962367 |
| *Raphia hookeri* | Arecaceae | 18 | 0,27 | 3 | 2,59 | 0,0157605 |
| *Di* | X | 11 | 0,16 | 3 | 2,59 | 0,01043125 |
| *Monodora sp.* | Annonaceae | 6 | 0,09 | 3 | 2,59 | 0,00622673 |
| Ema | X | 5 | 0,07 | 3 | 2,59 | 0,00532354 |
| *Napoleona leonensis* | Lechythidaceae | 103 | 1,52 | 3 | 2,59 | 0,0636579 |
| *Ficus exasperata* | Moraceae | 14 | 0,21 | 3 | 2,59 | 0,01277764 |
| *Ficus sp* | Moraceae | 5 | 0,07 | 3 | 2,59 | 0,00532354 |
| *Funtumia africana* | Apocynaceae | 14 | 0,21 | 3 | 2,59 | 0,01277764 |
| Houndje(abbey) | X | 32 | 0,47 | 3 | 2,59 | 0,02530029 |
| *Pterygota macrocarpa* | Sterculiaceae | 16 | 0,24 | 3 | 2,59 | 0,01428758 |
| *Cecropia peltata Linn.* | Cecropiaceae | 38 | 0,56 | 3 | 2,59 | 0,02907992 |
| *Albizia zygia* | Mimosaceae | 70 | 1,03 | 3 | 2,59 | 0,04725444 |
| *Kaya grandifolia* | Meliaceae | 4 | 0,06 | 2 | 1,72 | 0,00439061 |
| Bobolou | X | 5 | 0,07 | 2 | 1,72 | 0,00532354 |
| *Ficus goliath* | Moraceae | 2 | 0,03 | 2 | 1,72 | 0,00239999 |

| | | | | | | |
|---|---|---|---|---|---|---|
| Gnicherbier (attié) | X | 6 | 0,09 | 2 | 1,72 | 0,00622673 |
| Fêh | X | 11 | 0,16 | 2 | 1,72 | 0,01043125 |
| *Amphimas pteyocoipoides* | Cesalpiniaceae | 2 | 0,03 | 2 | 1,72 | 0,00239999 |
| Poe | X | 4 | 0,06 | 2 | 1,72 | 0,00439061 |
| Tin | X | 5 | 0,07 | 2 | 1,72 | 0,00532354 |
| Waha | X | 2 | 0,03 | 2 | 1,72 | 0,00239999 |
| Abôh | X | 1 | 0,01 | 1 | 0,86 | 0,00130233 |
| *Entandrophragma cylindricum* | Meliaceae | 10 | 0,15 | 1 | 0,86 | 0,00962367 |
| Hertinin(koyago) | X | 2 | 0,03 | 1 | 0,86 | 0,00239999 |
| *Mareya micrantha* | Euphorbiaceae | 1 | 0,01 | 1 | 0,86 | 0,00130233 |
| *Aningueria robusta* | Sapotaceae | 3 | 0,04 | 1 | 0,86 | 0,00342038 |
| *Napoleona leonensis* | Lechythidaceae | 3 | 0,04 | 1 | 0,86 | 0,00342038 |
| *Nauclea diderrichii* | Rubiaceae | 4 | 0,06 | 1 | 0,86 | 0,00439061 |
| *Bambusa vulgaris* | Poaceae (Gramineae) | 4 | 0,06 | 1 | 0,86 | 0,00439061 |
| Berdou | X | 3 | 0,04 | 1 | 0,86 | 0,00342038 |
| *Nesogordonia papaverifera* | Sterculiaceae | 1 | 0,01 | 1 | 0,86 | 0,00130233 |
| Ditinier | X | 1 | 0,01 | 1 | 0,86 | 0,00130233 |
| Diyinglario | X | 20 | 0,30 | 1 | 0,86 | 0,01720055 |
| *Albizia adianthifolia* | Mimosaceae | 30 | 0,44 | 1 | 0,86 | 0,02400488 |
| *Entandrophragma spp* | Sterculiaceae | 1 | 0,01 | 1 | 0,86 | 0,00130233 |
| *Olyra latifolia* | Poaceae (Gramineae) | 1 | 0,01 | 1 | 0,86 | 0,00130233 |
| Eflangui | X | 5 | 0,07 | 1 | 0,86 | 0,00532354 |
| Ehe | X | 1 | 0,01 | 1 | 0,86 | 0,00130233 |
| Hertinin(koyago) | X | 2 | 0,03 | 1 | 0,86 | 0,00239999 |
| *Ficus goliath* | Moraceae | 1 | 0,01 | 1 | 0,86 | 0,00130233 |
| *Delonix regia* | Cesalpiniaceae | 10 | 0,15 | 1 | 0,86 | 0,00962367 |
| Gbotopian | X | 1 | 0,01 | 1 | 0,86 | 0,00130233 |
| *Pycnanthus angolensis* | Myristicaceae | 4 | 0,06 | 1 | 0,86 | 0,00439061 |
| *Gmelina arborea.* | Verbenaceae | 2 | 0,03 | 1 | 0,86 | 0,00239999 |
| Heramba(agni) | X | 2 | 0,03 | 1 | 0,86 | 0,00239999 |
| *Irvingia gabonensis* | Irvingiraceae | 1 | 0,01 | 1 | 0,86 | 0,00130233 |
| *Terminalia scimperiana* | Combretaceae | 3 | 0,04 | 1 | 0,86 | 0,00342038 |
| *Morinda lucida* | Rubiaceae | 6 | 0,09 | 1 | 0,86 | 0,00622673 |
| Kpangba | X | 1 | 0,01 | 1 | 0,86 | 0,00130233 |
| Kparie | X | 1 | 0,01 | 1 | 0,86 | 0,00130233 |
| *Zanthoxylum gilletii* | Rutaceae | 2 | 0,03 | 1 | 0,86 | 0,00239999 |
| *Daniella oliveri* | Cesalpiniaceae | 1 | 0,01 | 1 | 0,86 | 0,00130233 |
| Micouzin | X | 2 | 0,03 | 1 | 0,86 | 0,00239999 |
| Negre | X | 1 | 0,01 | 1 | 0,86 | 0,00130233 |
| *Heritiera utilis* | X | 3 | 0,04 | 1 | 0,86 | 0,00342038 |
| *Discoglyprernna caloneura* | Euphorbiaceae | 6 | 0,09 | 1 | 0,86 | 0,00622673 |
| *Borassus aethiopum* | Palmaceae | 2 | 0,03 | 1 | 0,86 | 0,00239999 |
| *Xylopia aethiopica* | Annonaceae | 1 | 0,01 | 1 | 0,86 | 0,00130233 |

| | | | | | | |
|---|---|---|---|---|---|---|
| Tabayiri(malinke) | X | 1 | 0,01 | 1 | 0,86 | 0,00130233 |
| Tauro | X | 1 | 0,01 | 1 | 0,86 | 0,00130233 |
| *Phyllanthus discoideus* | Euphorbiaceae | 1 | 0,01 | 1 | 0,86 | 0,00130233 |
| Tene | X | 1 | 0,01 | 1 | 0,86 | 0,00130233 |

**S 3 : Residual and naturally regenerated forest species recorded in cocoa plots of Belt 2**

| Scientific names | Botanical families | Number of individuals (n=8947) | Representativeness % | Number of plots (n=178) | % of plots | Shannon index (n=2,8354) |
|---|---|---|---|---|---|---|
| *Elaeis guineensis* | Palmaceae | 2439 | 27,26 | 136 | 76,40 | 0,354313386 |
| *Ceiba pentandra* | Bombacaceae | 189 | 2,11 | 66 | 37,08 | 0,081483706 |
| *Ficus exasperata* | Moraceae | 479 | 5,35 | 54 | 30,34 | 0,156724226 |
| *Chlorophora excelsa/C, regia* | Moraceae | 145 | 1,62 | 53 | 29,78 | 0,066808905 |
| *Cola nitida* | Sterculiaceae | 489 | 5,47 | 43 | 24,16 | 0,158866851 |
| *Ricinodendron heudelotii* | Euphorbiaceae | 97 | 1,08 | 31 | 17,42 | 0,049051433 |
| *Triplochiton scleroxylon* | Sterculiaceae | 77 | 0,86 | 24 | 13,48 | 0,040924963 |
| *Antiaris africana* | Moraceae | 55 | 0,61 | 22 | 12,36 | 0,031300516 |
| *Bombax buenopozense* | Bombacaceae | 45 | 0,50 | 22 | 12,36 | 0,026618811 |
| *Nesogordonia papaverifera* | Sterculiaceae | 29 | 0,32 | 12 | 6,74 | 0,018578468 |
| *Cola gigantea* | Sterculiaceae | 25 | 0,28 | 10 | 5,62 | 0,016430641 |
| *Terminalia superba* | Combretaceae | 16 | 0,18 | 10 | 5,62 | 0,011313709 |
| *Cordia platythrysa* | Boraginaceae | 23 | 0,26 | 8 | 4,49 | 0,015330538 |
| *Albizzia spp* | Mimosaceae | 78 | 0,87 | 7 | 3,93 | 0,041343964 |
| *Alstonia boonei* | Apocynaceae | 11 | 0,12 | 7 | 3,93 | 0,008238847 |
| *Spathodea campanulata* | Bignoniaceae | 41 | 0,46 | 6 | 3,37 | 0,024679285 |
| *Funtumia africana* | Apocynaceae | 23 | 0,26 | 6 | 3,37 | 0,015330538 |
| *Morinda lucida.* | Rubiaceae | 28 | 0,31 | 6 | 3,37 | 0,018047651 |
| *Sterculia tragacantha* | Sterculiaceae | 5 | 0,06 | 6 | 3,37 | 0,004185557 |
| *Mansonia altissima* | Sterculiaceae | 36 | 0,40 | 5 | 2,81 | 0,02219291 |
| *Terminalia ivorensis* | Combretaceae | 6 | 0,07 | 5 | 2,81 | 0,004900401 |
| *Ficus capensis* | Moraceae | 30 | 0,34 | 4 | 2,25 | 0,01910543 |
| *Tieghemella heckelii* | Sapotaceae | 9 | 0,10 | 3 | 1,69 | 0,006942734 |
| *Celtis zenkeri* | Ulmaceae | 5 | 0,06 | 3 | 1,69 | 0,004185557 |
| *Coffea canephora* | Rubiaceae | 161 | 1,80 | 3 | 1,69 | 0,072297389 |
| *Entandrophragma utile* | Meliaceae | 3 | 0,03 | 3 | 1,69 | 0,002682618 |
| *Irvingia gabonensis* | Irvingiraceae | 13 | 0,15 | 3 | 1,69 | 0,009494089 |
| *Lannea acida* | Anacardiaceae | 16 | 0,18 | 3 | 1,69 | 0,011313709 |
| *Napoleona leonensis* | Lechythidaceae | 10 | 0,11 | 3 | 1,69 | 0,007596388 |
| *Newbouldia laevis* | Bignoniaceae | 22 | 0,25 | 3 | 1,69 | 0,014773297 |
| *Pilper guineesis* | Moraceae | 111 | 1,24 | 3 | 1,69 | 0,054458401 |
| *Piptadeniastrum africanum* | Mimosaceae | 3 | 0,03 | 3 | 1,69 | 0,002682618 |

| | | | | | | |
|---|---|---|---|---|---|---|
| *Pygnanthus angolensis* | Myristicaceae | 5 | 0,06 | 3 | 1,69 | 0,004185557 |
| *Antiaris toxicaria* | Moraceae | 7 | 0,08 | 2 | 1,12 | 0,005596529 |
| *Blighia welwitschii* | Sapindaceae | 6 | 0,07 | 2 | 1,12 | 0,004900401 |
| *kaya grandifolia* | Meliaceae | 5 | 0,06 | 2 | 1,12 | 0,004185557 |
| *Pycnanthus angolensis* | Myristicaceae | 13 | 0,15 | 2 | 1,12 | 0,009494089 |
| *Xylopia aethiopica* | Annonaceae | 8 | 0,09 | 2 | 1,12 | 0,006276635 |
| *Albizia ferruginea* | Mimosaceae | 12 | 0,13 | 1 | 0,56 | 0,00887113 |
| *Anthocleista nobilis* | Loganiaceae | 7 | 0,08 | 1 | 0,56 | 0,005596529 |
| *Blighia sapida* | Sapindaceae | 1 | 0,01 | 1 | 0,56 | 0,001016997 |
| *Borassus flabellifer* | Palmaceae | 1 | 0,01 | 1 | 0,56 | 0,001016997 |
| *Carapa procera* | Meliaceae | 10 | 0,11 | 1 | 0,56 | 0,007596388 |
| *Cola grandifolia* | Sterculiaceae | 1 | 0,01 | 1 | 0,56 | 0,001016997 |
| *Cola grandis* | Sterculiaceae | 20 | 0,22 | 1 | 0,56 | 0,013643325 |
| *Cola heterophylla* | Sterculiaceae | 15 | 0,17 | 1 | 0,56 | 0,010714804 |
| *Distemonanthus benthamianus* | Caesalpiniaceae | 1 | 0,01 | 1 | 0,56 | 0,001016997 |
| *Erythrima senegalensis* | Papillonnaceae | 1 | 0,01 | 1 | 0,56 | 0,001016997 |
| *Fagara xanthoxyloides* | Rutaceae | 2 | 0,02 | 1 | 0,56 | 0,001879049 |
| *Ficus dicranostyla* | Moraceae | 1 | 0,01 | 1 | 0,56 | 0,001016997 |
| *Ficus mucuso* | Moraceae | 3 | 0,03 | 1 | 0,56 | 0,002682618 |
| *Ficus vogeii* | Moraceae | 1 | 0,01 | 1 | 0,56 | 0,001016997 |
| *Gmelina arborea* | Verbenaceae | 10 | 0,11 | 1 | 0,56 | 0,007596388 |
| *Hevea brasiliensis* | Euphorbiaceae | 20 | 0,22 | 1 | 0,56 | 0,013643325 |
| *Luffa sp* | Cucurbitaceae | 10 | 0,11 | 1 | 0,56 | 0,007596388 |
| *Milicia regia* | Moraceae | 1 | 0,01 | 1 | 0,56 | 0,001016997 |
| *Myrianthus arboreus* | Cecropiaceae | 1 | 0,01 | 1 | 0,56 | 0,001016997 |
| *Napoleona leonensis* | Lechythidaceae | 1 | 0,01 | 1 | 0,56 | 0,001016997 |
| *Parkia bicolore* | Mimosaceae | 5 | 0,06 | 1 | 0,56 | 0,004185557 |
| Parkia biglobosa | Mimosaceae | 4 | 0,04 | 1 | 0,56 | 0,003448208 |
| *Pentaclethia macrophila* | Mimosaceae | 2 | 0,02 | 1 | 0,56 | 0,001879049 |
| *Peteysanthus macrocarpus* | Lechythidaceae | 1 | 0,01 | 1 | 0,56 | 0,001016997 |
| *Raphia hookeri.* | Palmaceae | 1 | 0,01 | 1 | 0,56 | 0,001016997 |
| *Solanum rugosum* | Solanaceae | 12 | 0,13 | 1 | 0,56 | 0,00887113 |
| *Sterculia africana* | Sterculiaceae | 4 | 0,04 | 1 | 0,56 | 0,003448208 |
| *Treculia africana* | Moraceae | 6 | 0,07 | 1 | 0,56 | 0,004900401 |
| *Xylopia elliotii* | Annonaceae | 4 | 0,04 | 1 | 0,56 | 0,003448208 |

#### S 4 : Residual and naturally regenerated forest species recorded in cocoa plots of Belt 3.

| Scientific names | Botanical families | Number of individuals (n=10492) | Representa tiveness % | Number of plots (n=175) | % of plots | Shannon index (n=2,9144) |
|---|---|---|---|---|---|---|
| *Elaeis guineensis* | Palmaceae | 3003 | 28,62 | 64 | 36,57 | 0,358059 |
| *Ficus exasperata* | Moraceae | 130 | 1,24 | 50 | 28,57 | 0,054404 |
| *Alstonia boonei* | Apocynaceae | 57 | 0,54 | 27 | 15,43 | 0,028333 |

| | | | | | | |
|---|---|---|---|---|---|---|
| *Coffea canephora* | Rubiaceae | 79 | 0,75 | 21 | 12,00 | 0,036811 |
| *Discoglyprernna caloneura* | Euphorbiaceae | 32 | 0,30 | 20 | 11,43 | 0,017667 |
| *Nesogordonia papaverifera* | Sterculiaceae | 59 | 0,56 | 19 | 10,86 | 0,029134 |
| *Albizzia spp* | Mimosaceae | 26 | 0,25 | 17 | 9,71 | 0,014869 |
| *Mansonia altissima* | Sterculiaceae | 137 | 1,31 | 17 | 9,71 | 0,056649 |
| *Cola grandis* | Sterculiaceae | 14 | 0,13 | 13 | 7,43 | 0,008832 |
| *Napoleona leonensis* | Lechythidaceae | 116 | 1,11 | 8 | 4,57 | 0,049805 |
| *Morinda lucida Benth.* | Rubiaceae | 9 | 0,09 | 6 | 3,43 | 0,006057 |
| *Bombax buenopozense.* | Bombacaceae | 10 | 0,10 | 5 | 2,86 | 0,006630 |
| *Cedrela odorata* | Meliaceae | 4 | 0,04 | 5 | 2,86 | 0,003001 |
| *Albizia ferruginea* | Mimosaceae | 8 | 0,08 | 4 | 2,29 | 0,005474 |
| *Holarrhena floribunda* | Apocynaceae | 12 | 0,11 | 4 | 2,29 | 0,007747 |
| *Irvingia gabonensis* | Irvingiraceae | 11 | 0,10 | 4 | 2,29 | 0,007193 |
| *Myrianthus arboreus* | Cecropiaceae | 122 | 1,16 | 4 | 2,29 | 0,051795 |
| *Peteysanthus macrocarpus* | Lechythidaceae | 18 | 0,17 | 3 | 1,71 | 0,010925 |
| *Ceiba pentandra* | Bombacaceae | 228 | 2,17 | 3 | 1,71 | 0,083208 |
| *Chlorophora excelsa/C, regia* | Moraceae | 105 | 1,00 | 3 | 1,71 | 0,046079 |
| *Cola cordifolia* | Sterculiaceae | 4 | 0,04 | 3 | 1,71 | 0,003001 |
| *Fagara xanthoxyloides* | Rutaceae | 5 | 0,05 | 3 | 1,71 | 0,003645 |
| *Musanga cecropioides* | Cecropiaceae | 8 | 0,08 | 3 | 1,71 | 0,005474 |
| *Triplochiton scleroxylon* | Sterculiaceae | 62 | 0,59 | 3 | 1,71 | 0,030322 |
| *Aningueria robusta* | Sapotaceae | 20 | 0,19 | 2 | 1,14 | 0,011938 |
| *Antiaris africana* | Moraceae | 74 | 0,71 | 2 | 1,14 | 0,034943 |
| *Daniella oliveri* | Cesalpiniaceae | 14 | 0,13 | 2 | 1,14 | 0,008832 |
| *Ficus goliath* | Moraceae | 3 | 0,03 | 2 | 1,14 | 0,002333 |
| *Ficus mucuso* | Moraceae | 39 | 0,37 | 2 | 1,14 | 0,020797 |
| *Gaycinia kola* | Guttiferaceae | 6 | 0,06 | 2 | 1,14 | 0,004270 |
| Gboudihi(dida) | X | 8 | 0,08 | 2 | 1,14 | 0,005474 |
| Tanvanwaka | X | 6 | 0,06 | 2 | 1,14 | 0,004270 |
| *Raphia hookeri* | Arecaceae | 10 | 0,10 | 2 | 1,14 | 0,006630 |
| *Albizia adianthifolia* | Mimosaceae | 8 | 0,08 | 1 | 0,57 | 0,005474 |
| *Albizzia sp* | Mimosaceae | 7 | 0,07 | 1 | 0,57 | 0,004879 |
| *Amphimas pteyocoipoides* | Cesalpiniaceae | 3 | 0,03 | 1 | 0,57 | 0,002333 |
| *Anthocleista nobilis* | Loganiaceae | 3 | 0,03 | 1 | 0,57 | 0,002333 |
| *Antiaris toxicaria* | Moraceae | 15 | 0,14 | 1 | 0,57 | 0,009365 |
| *Artoccarpus communis* | Moraceae | 10 | 0,10 | 1 | 0,57 | 0,006630 |
| *Borassus aethiopum* | Palmaceae | 1 | 0,01 | 1 | 0,57 | 0,000882 |
| *Borassus flabellifer* | Palmaceae | 3 | 0,03 | 1 | 0,57 | 0,000882 |
| *Celtis zenkeri* | Ulmaceae | 31 | 0,30 | 1 | 0,57 | 0,002333 |
| *Cola heterophylla.* | Sterculiaceae | 7 | 0,07 | 1 | 0,57 | 0,017209 |
| *Cola lateritia* | Sterculiaceae | 1 | 0,01 | 1 | 0,57 | 0,004879 |
| *Cola nitida* | Sterculiaceae | 384 | 3,66 | 1 | 0,57 | 0,000882 |
| *Cordia platythrysa* | Boraginaceae | 1 | 0,01 | 1 | 0,57 | 0,121060 |

| | | | | | | |
|---|---|---|---|---|---|---|
| *Entandrophragma angolense* | Meliaceae | 16 | 0,15 | 1 | 0,57 | 0,000882 |
| *Entandrophragma spp* | Sterculiaceae | 5 | 0,05 | 1 | 0,57 | 0,009891 |
| *Entandrophragma utile* | Meliaceae | 3 | 0,03 | 1 | 0,57 | 0,003645 |
| *Erythrima senegalensis* | Papillonnaceae | 3 | 0,03 | 1 | 0,57 | 0,002333 |
| *Ficus capensis* | Moraceae | 14 | 0,13 | 1 | 0,57 | 0,002333 |
| *Ficus sp* | Moraceae | 4 | 0,04 | 1 | 0,57 | 0,008832 |
| *Jatropha curcas* | Euphorbiaceae | 9 | 0,09 | 1 | 0,57 | 0,003001 |
| *Kaya anthothéca* | Meliaceae | 2 | 0,02 | 1 | 0,57 | 0,006057 |
| *kaya grandifolia* | Meliaceae | 41 | 0,39 | 1 | 0,57 | 0,001633 |
| *Margaritaria discoidea* | Euphorbiaceae | 1 | 0,01 | 1 | 0,57 | 0,021668 |
| *Nauclea diderrichii.* | Rubiaceae | 1 | 0,01 | 1 | 0,57 | 0,000882 |
| *Newbouldia laevis* | Bignoniaceae | 3 | 0,03 | 1 | 0,57 | 0,000882 |
| *Parkia bicolore* | Mimosaceae | 1 | 0,01 | 1 | 0,57 | 0,002333 |
| *Piptadeniastrum africanum* | Mimosaceae | 3 | 0,03 | 1 | 0,57 | 0,000882 |
| *Pycnanthus angolensis* | Myristicaceae | 27 | 0,26 | 1 | 0,57 | 0,002333 |
| *Pygnanthus angolensis* | Myristicaceae | 3 | 0,03 | 1 | 0,57 | 0,015344 |
| *Ricinodendron heudelotii* | Euphorbiaceae | 55 | 0,52 | 1 | 0,57 | 0,002333 |
| *Sperogordia nazogarpas* | Rubiaceae | 17 | 0,16 | 1 | 0,57 | 0,027526 |
| *Sterculia tragacantha* | Sterculiaceae | 1 | 0,01 | 1 | 0,57 | 0,010411 |
| *Terminalia ivorensis* | Combretaceae | 14 | 0,13 | 1 | 0,57 | 0,000882 |
| *Terminalia superba* | Combretaceae | 67 | 0,64 | 1 | 0,57 | 0,032272 |
| *Tieghemella heckelii .* | Sapotaceae | 5 | 0,05 | 1 | 0,57 | 0,003645 |
| *Treculia africana* | Moraceae | 5 | 0,05 | 1 | 0,57 | 0,003645 |
| *Zanthoxylum Zanthoxyloides* | Rutaceae | 2 | 0,02 | 1 | 0,57 | 0,001633 |

**S 5: Species richness and abundance by family in Belt 1**.

| Botanical families | Number of species | Number of individuals (n=6773) | Representa tiveness (%) | Number of plots or repetitions (n=116) | % of plots |
|---|---|---|---|---|---|
| Anacardiaceae | 3 | 571 | 8,43 | 74 | 8,60 |
| Annonaceae | 3 | 11 | 0,16 | 6 | 0,70 |
| Apocynaceae | 4 | 184 | 2,72 | 38 | 4,42 |
| Arecaceae | 1 | 18 | 0,27 | 3 | 0,35 |
| Bombacaceae | 4 | 257 | 3,79 | 69 | 8,02 |
| Cecropiaceae | 2 | 83 | 1,23 | 9 | 1,05 |
| Cesalpiniaceae | 5 | 160 | 2,36 | 19 | 2,21 |
| Combretaceae | 3 | 513 | 7,57 | 98 | 11,40 |
| Euphorbiaceae | 6 | 77 | 1,14 | 26 | 3,02 |
| Inconnu | 29 | 154 | 2,27 | 44 | 5,12 |
| Irvingiraceae | 1 | 1 | 0,01 | 1 | 0,12 |
| Lauraceae | 1 | 670 | 9,89 | 91 | 10,58 |
| Lechythidaceae | 2 | 106 | 1,57 | 4 | 0,47 |

| | | | | | |
|---|---|---|---|---|---|
| Meliaceae | 4 | 86 | 1,27 | 25 | 2,91 |
| Mimosaceae | 3 | 107 | 1,58 | 8 | 0,93 |
| Moraceae | 7 | 178 | 2,63 | 53 | 6,16 |
| Musaceae | 1 | 20 | 0,30 | 2 | 0,23 |
| Myristicaceae | 2 | 115 | 1,70 | 14 | 1,63 |
| Myrtaceae | 2 | 49 | 0,72 | 13 | 1,51 |
| Palmaceae | 4 | 2259 | 33,35 | 134 | 15,58 |
| Poaceae | 2 | 5 | 0,07 | 2 | 0,23 |
| Rubiaceae | 3 | 496 | 7,32 | 7 | 0,81 |
| Rutaceae | 5 | 362 | 5,34 | 71 | 8,26 |
| Sapindaceae | 1 | 10 | 0,15 | 1 | 0,12 |
| Sapotaceae | 1 | 3 | 0,04 | 1 | 0,12 |
| Sterculiaceae | 8 | 253 | 3,74 | 45 | 5,23 |
| Verbenaceae | 2 | 25 | 0,37 | 2 | 0,23 |
| Total | 80 + 29 | 6773 | 100 | 860 | 100,00 |

**S 6 : Species richness and abundance by family in Belt 2**

| Families botaniques | Number of species | Number of individuals (n=8947) | Representa tiveness (%) | Number of plots or repetitions (n=178) | % of plots |
|---|---|---|---|---|---|
| Rutaceae | 5 | 1197 | 13,38 | 213 | 17,78 |
| Palmaceae | 4 | 2527 | 28,25 | 151 | 12,60 |
| Lauraceae | 1 | 1077 | 12,04 | 150 | 12,52 |
| Moraceae | 12 | 840 | 9,39 | 144 | 12,02 |
| Sterculiaceae | 10 | 701 | 7,84 | 104 | 8,68 |
| Bombacaceae | 3 | 239 | 2,67 | 90 | 7,51 |
| Anacardiaceae | 4 | 1093 | 12,22 | 80 | 6,68 |
| Euphorbiaceae | 3 | 318 | 3,56 | 60 | 5,01 |
| Myrtaceae | 1 | 115 | 1,29 | 46 | 3,84 |
| Mimosaceae | 7 | 134 | 1,50 | 16 | 1,34 |
| Combretaceae | 2 | 22 | 0,25 | 15 | 1,25 |
| Apocynaceae | 2 | 34 | 0,38 | 13 | 1,09 |
| Annonaceae | 3 | 26 | 0,29 | 11 | 0,92 |
| Bignoniaceae | 2 | 63 | 0,70 | 9 | 0,75 |
| Rubiaceae | 2 | 189 | 2,11 | 9 | 0,75 |
| Boraginaceae | 1 | 23 | 0,26 | 8 | 0,67 |
| Sapindaceae | 4 | 28 | 0,31 | 7 | 0,58 |
| Verbenaceae | 2 | 128 | 1,43 | 7 | 0,58 |
| Meliaceae | 3 | 18 | 0,20 | 6 | 0,50 |
| Lechythidaceae | 3 | 12 | 0,13 | 5 | 0,42 |
| Myristicaceae | 2 | 18 | 0,20 | 5 | 0,42 |
| Cesalpiniaceae | 2 | 13 | 0,15 | 3 | 0,25 |
| Irvingiraceae | 1 | 13 | 0,15 | 3 | 0,25 |
| Ulmaceae | 1 | 5 | 0,06 | 3 | 0,25 |

| | | | | | |
|---|---|---|---|---|---|
| Caesalpiniaceae | 1 | 1 | 0,01 | 1 | 0,08 |
| Cecropiaceae | 1 | 1 | 0,01 | 1 | 0,08 |
| Cucurbitaceae | 1 | 10 | 0,11 | 1 | 0,08 |
| Loganiaceae | 1 | 7 | 0,08 | 1 | 0,08 |
| Moringaceae | 1 | 1 | 0,01 | 1 | 0,08 |
| Papillonnaceae | 1 | 1 | 0,01 | 1 | 0,08 |
| Total | 86 | 8947 | 100 | 1164 | 100 |

#### S 7 : Species richness and abundance by family in Belt 3

| Botanical families | Number of species | Number of individuals (n=10493) | Representa tiveness (%) | Number of plots or repetitions (n=175) | % of Parcelle |
|---|---|---|---|---|---|
| Mimosaceae | 8 | 3086 | 29,41 | 95 | 16,21 |
| Moraceae | 10 | 1061 | 10,11 | 67 | 11,43 |
| Myrtaceae | 2 | 1185 | 11,29 | 65 | 11,09 |
| Irvingiraceae | 1 | 152 | 1,45 | 64 | 10,92 |
| Inconnu | 2 | 174 | 1,66 | 44 | 7,51 |
| Bombacaceae | 3 | 68 | 0,65 | 29 | 4,95 |
| Rutaceae | 6 | 909 | 8,66 | 27 | 4,61 |
| Palmaceae | 4 | 268 | 2,55 | 25 | 4,27 |
| Lechythidaceae | 2 | 283 | 2,70 | 22 | 3,75 |
| Meliaceae | 6 | 421 | 4,01 | 19 | 3,24 |
| Bignoniaceae | 1 | 26 | 0,25 | 17 | 2,90 |
| Sterculiaceae | 10 | 444 | 4,23 | 17 | 2,90 |
| Rubiaceae | 4 | 247 | 2,35 | 16 | 2,73 |
| Euphorbiaceae | 6 | 468 | 4,46 | 13 | 2,22 |
| Anacardiaceae | 3 | 33 | 0,31 | 11 | 1,88 |
| Cesalpiniaceae | 5 | 237 | 2,26 | 8 | 1,37 |
| Combretaceae | 2 | 11 | 0,10 | 6 | 1,02 |
| Apocynaceae | 2 | 16 | 0,15 | 5 | 0,85 |
| Musaceae | 2 | 20 | 0,19 | 5 | 0,85 |
| Boraginaceae | 1 | 10 | 0,10 | 4 | 0,68 |
| Cecropiaceae | 2 | 30 | 0,29 | 3 | 0,51 |
| Guttiferaceae | 1 | 105 | 1,00 | 3 | 0,51 |
| Lauraceae | 1 | 980 | 9,34 | 3 | 0,51 |
| Loganiaceae | 1 | 4 | 0,04 | 3 | 0,51 |
| Sapotaceae | 2 | 13 | 0,12 | 3 | 0,51 |
| Ulmaceae | 1 | 62 | 0,59 | 3 | 0,51 |
| Bromeliaceae | 1 | 20 | 0,19 | 2 | 0,34 |
| Myristicaceae | 2 | 43 | 0,41 | 2 | 0,34 |
| Annonaceae | 1 | 1 | 0,01 | 1 | 0,17 |
| Arecaceae | 1 | 7 | 0,07 | 1 | 0,17 |
| Papillonnaceae | 1 | 80 | 0,76 | 1 | 0,17 |
| Sapindaceae | 1 | 27 | 0,26 | 1 | 0,17 |

| | | | | | |
|---|---|---|---|---|---|
| Verbenaceae | 1 | 2 | 0,02 | 1 | 0,17 |
| Total | 94 | 10493 | 100 | 586 | 100 |

#### S 8 : List of trees preferred by cocoa farmers for belt 1

| Common names or Other names | Scientific names | Botanical families | Number of plots or repetitions (n=116) | % total of citations |
|---|---|---|---|---|
| Ekoué(agni) | *Monodora sp.* | Annonaceae | 75 | 64,66 |
| Ilomba | *Pygnanthus angolensis* | Myristicaceae | 55 | 47,41 |
| Fraké (fla, yabi, plat) | *Terminalia superba* | Combretaceae | 47 | 40,52 |
| Sipo | *Entandrophragma utile* | Meliaceae | 18 | 15,52 |
| Akpi | *Ricinodendron heudelotii* | Euphorbiaceae | 16 | 13,79 |
| Framiré (tin) | *Terminalia ivorensis* | Combretaceae | 9 | 7,76 |
| Amien/ermien/corkpê | *Alstonia boonei* | Apocynaceae | 8 | 6,90 |
| Moringa | *Moringa oleifera* | Moringaceae | 8 | 6,90 |
| Fromager(gnin/gounga/djo/gôh) | *Ceiba pentandra* | Bombacaceae | 7 | 6,03 |
| Glyricidia | *Gliricidia sepium* | Cesalpiniaceae | 5 | 4,31 |
| Samba/kpatayobouet/coffa | *Triplochiton scleroxylon* | Sterculiaceae | 5 | 4,31 |
| Kplé/kakrou/diamon/boborou | *Irvingia gabonensis* | Irvingiraceae | 4 | 3,45 |
| Ako(offi/offouin) | *Antiaris africana* | Moraceae | 3 | 2,59 |
| Doukouman | *Entandrophragma spp* | Sterculiaceae | 3 | 2,59 |
| Heramba | *Heramba* | X | 3 | 2,59 |
| Acajou | *Kaya grandifolia* | Meliaceae | 3 | 2,59 |
| Acacia mangium | *Acacia mangium* | Mimosaceae | 2 | 1,72 |
| Walê | *Cola gigantea* | Sterculiaceae | 2 | 1,72 |
| Yabi | inconnu | X | 2 | 1,72 |
| Bois bété(glonhi) | *Mansonia altissima* | Sterculiaceae | 2 | 1,72 |
| Aya/cotibé/bois rouge | *Nesogordonia papaverifera* | Sterculiaceae | 2 | 1,72 |
| Karité | *Vitellaria paradoxa* | Sapotaceae | 2 | 1,72 |
| Aniégré(ahinclin) | *Aningueria robusta* | Sapotaceae | 1 | 0,86 |
| Bofouin | *Antiaris toxicaria* | Moraceae | 1 | 0,86 |
| Iroko(alla/ago/golyiri) | *Milicia excelsa/C, regia* | Moraceae | 1 | 0,86 |
| Tiama | *Entandrophragma angolense* | Meliaceae | 1 | 0,86 |
| Django/djangoe/ficus etrangleur | *Ficus goliath* | Moraceae | 1 | 0,86 |
| Niangon | *Heritiera utilis* | X | 1 | 0,86 |
| Aonialé | Aonialé | X | 1 | 0,86 |
| Bossoman | Bossoman | X | 1 | 0,86 |
| Ehouré | Ehouré | X | 1 | 0,86 |
| Gni | *Milicia regia* | Moraceae | 1 | 0,86 |
| N'deffou/diffou | *Morus mesozygia* | Moraceae | 1 | 0,86 |
| Ayêclê/ayinglê/ahinclin | *Napoleona leonensis* | Lechythidaceae | 1 | 0,86 |
| Mirabelle(tronma/ouinyiri) | *Spondias mombin* | Anacardiaceae | 1 | 0,86 |

| | | | | |
|---|---|---|---|---|
| Total | 35 | 16 | 116 | 100 |

**S 9 : List of trees preferred by cocoa farmers in Belt 2**.

| Common names or Other names | Scientific names | Botanical families | Number of plots or repetitions (n=178) | % of total citations |
|---|---|---|---|---|
| Iroko(alla/ago/golyiri) | *Milicia excelsa/M, regia* | Moraceae | 57 | 32,02 |
| Fraké(fla/plat/yabi) | *Terminalia superba* | Combretaceae | 35 | 19,66 |
| Akpi | *Ricinodendron heudelotii* | Euphorbiaceae | 28 | 15,73 |
| Avocatier | *Percea americana* | Lauraceae | 27 | 15,17 |
| Fromager(gnin/gounga/djo/gôh) | *Ceiba pentandra* | Bombacaceae | 26 | 14,61 |
| Manguier(amango) | *Mangifera indica* | Anacardiaceae | 24 | 13,48 |
| Oranger(n'dédé) | *Citrus sinensis* | Rutaceae | 16 | 8,99 |
| Bois bété(glonhi) | *Mansonia altissima* | Sterculiaceae | 16 | 8,99 |
| Samba(kpatayobouet/coffa) | *Triplochiton scleroxylon* | Sterculiaceae | 8 | 4,49 |
| Anacardier | *Anacardium occidentale* | Anacardiaceae | 6 | 3,37 |
| Ploussou(tamtam) | *Cordia platythrysa* | Boraginaceae | 6 | 3,37 |
| Aloma | *Ficus capensis* | Moraceae | 5 | 2,81 |
| Ako(offi/offouin) | *Antiaris africana* | Moraceae | 4 | 2,25 |
| Gnangnoui | Gnangnoui | X | 3 | 1,69 |
| Sipo | *Entandrophragma utile* | Meliaceae | 3 | 1,69 |
| Yêclê/yêglê/yinglè/gnêglê | *Ficus exasperata* | Moraceae | 3 | 1,69 |
| Mirabelle(tronma/ouinyiri) | *Spondias mombin* | Anacardiaceae | 2 | 1,12 |
| Amien/ermien/corkpê | *Alstonia boonei* | Apocynaceae | 2 | 1,12 |
| Kapokier (kpouka/gbouka/kapoté) | *Bombax buenopozense* | Bombacaceae | 2 | 1,12 |
| Glyricidia | *Gliricidia sepium* | Cesalpiniaceae | 2 | 1,12 |
| Acajou | *Kaya grandifolia* | Meliaceae | 2 | 1,12 |
| Agnissanssan/ahissian | *Pilper guineesis* | Moraceae | 2 | 1,12 |
| Ilomba | *Pygnanthus angolensis* | Myristicaceae | 2 | 1,12 |
| Koya | *Morinda lucida* | Rubiaceae | 2 | 1,12 |
| Mandarinier | *Citrus reculata* | Rutaceae | 2 | 1,12 |
| Colatier(leu/colayiri) | *Cola nitida* | Sterculiaceae | 2 | 1,12 |
| Walê | *Cola cordifolia* | Sterculiaceae | 2 | 1,12 |
| Biébiésrélé | *Spathodea campanulata* | Bignoniaceae | 1 | 0,56 |
| Lingué (lagrê) | *Afzelia africana* | Caesalpiniaceae | 1 | 0,56 |
| Trikpor | *Myrianthus arboreus* | Cecropiaceae | 1 | 0,56 |
| Wodjé/akêdê/angama/kakahikpé/odjé | *Mareya micrantha.* | Euphorbiaceae | 1 | 0,56 |
| Kouêmankouê | inconnu | X | 1 | 0,56 |
| Kplé/kakrou/diamon/boborou | *Irvingia gabonensis* | Irvingiraceae | 1 | 0,56 |
| Abale/tanvanwaka | *Peteysanthus macrocarpus* | Lechythidaceae | 1 | 0,56 |
| Acacia | *Auriculo formis* | Mimosaceae | 1 | 0,56 |
| Dabeman(gblégblé/gbloungbloun) | *Piptadeniastrum africanum* | Mimosaceae | 1 | 0,56 |
| Kpatakpla | *Pentaclethia macrophila* | Mimosaceae | 1 | 0,56 |

| | | | | |
|---|---|---|---|---|
| Yatanza | *Albizia ferruginea* | Mimosaceae | 1 | 0,56 |
| Ficus spp | *Ficus spp* | Moraceae | 1 | 0,56 |
| Lokpro/lokplo/loungblo/avissanssan | *Ficus mucuso* | Moraceae | 1 | 0,56 |
| Yêclê/yêglê/yinglè/gnêglê | *Ficus exasperata* | Moraceae | 1 | 0,56 |
| Aya/cotibé/bois rouge | *Nesogordonia papaverifera* | Sterculiaceae | 1 | 0,56 |
| Mlémlédou/blebleadou | *Sterculia africana* | Sterculiaceae | 1 | 0,56 |
| Teck | *Tecktona grandis* | Verbenaceae | 1 | 0,56 |
| Total | 46 | 20 | 178 | 100 |

#### S 10 : List of trees preferred by cocoa farmers in Belt 3.

| Common names or Other names | Scientific names | Botanical families | Number of plots or repetitions (n=175) | % total of Citations |
|---|---|---|---|---|
| Iroko | *Chlorophora excelsa/C. regia* | Moraceae | 64 | 36,46 |
| Akpi | *Ricinodendron heudelotii* | Euphorbiaceae | 37 | 20,99 |
| Fromager | *Ceiba pentadra* | Bombacaceae | 37 | 20,99 |
| Fraké | *Terminalia superba* | Combretaceae | 36 | 20,44 |
| Samba | *Triplochiton scleroxylon* | Sterculiaceae | 23 | 13,26 |
| Glyricidia | *Gliricidia sepium* | Cesalpiniaceae | 14 | 7,73 |
| Kotibé | *Nesogordina papaverifera* | Sterculiaceae | 14 | 7,73 |
| Amien | *Alstonia boonei* | Apocynaceae | 12 | 6,63 |
| Acajou | *Kaya anthotheca/K grandifolia* | Meliaceae | 9 | 4,97 |
| Ako | *Antiaris africana* | Moraceae | 8 | 4,42 |
| Bété | *Mansonia altisima* | Sterculiaceae | 8 | 4,42 |
| Tiama | *Endophragma angolense* | Meliaceae | 8 | 4,42 |
| Avocatier | *Persea Americana* | Lauraceae | 7 | 3,87 |
| Parasolier (Agboui/Egbouin) | *Musenga cecropioides* | Moraceae | 6 | 3,31 |
| Framiré | *Terminalia ivorensis* | Combretaceae | 6 | 3,31 |
| Oranger | *Citrus sinensis* | Rutaceae | 6 | 3,31 |
| Teck | *Tecktona grandis* | Verbenaceae | 6 | 3,31 |
| Koto | *Pterygota macrocarpa* | Sterculiaceae | 5 | 2,76 |
| Manguier | *Magifera indica* | Anacardiaceae | 5 | 2,76 |
| Mirabellier | *Spondias mombin* | Anacardiaceae | 4 | 2,21 |
| Koya | *Morida lucida* | Rubiaceae | 3 | 1,66 |
| Drédré | inconnu | X | 2 | 1,1 |
| Kosipo | *Entandophragma candollei* | Meliaceae | 2 | 1,1 |
| Kplé | *Irvingia gabonensis* | Irvingiraceae | 2 | 1,1 |
| Muti | *Ficus vogeii* | Moraceae | 2 | 1,1 |
| Ayinglè | *Napoleona leonensis* | Lechythidaceae | 2 | 1,1 |
| Palmier | *Elaeis guineensis* | Palmaceae | 2 | 1,1 |
| Sipo | *Endophragma utile* | Meliaceae | 2 | 1,1 |
| Agnissanssan | *Ficus mucuso* | Moraceae | 1 | 0,55 |
| Aloma | *Ficus capensis* | Moraceae | 1 | 0,55 |
| Anacardier | *Anacardium occidentale* | Anacardiaceae | 1 | 0,55 |

| | | | | |
|---|---|---|---|---|
| Asan | *Celtis zenkeri* | Ulmaceae | 1 | 0,55 |
| Baobab | *Andansonia digitata* | Bombacaceae | 1 | 0,55 |
| Bobla | inconnu | X | 1 | 0,55 |
| Bossé | *Guarea* cedrata/G.thompsoneii | Meliaceae | 1 | 0,55 |
| Gnopoui | inconnu | X | 1 | 0,55 |
| Mandarine | *Citrus reculata* | Rutaceae | 1 | 0,55 |
| Neem | *Azadirachta indica* | Meliaceae | 1 | 0,55 |
| Walê | *Cola cordifolia/ C. lateritia* | Sterculiaceae | 1 | 0,55 |
| Total | 39 | 17 | 175 | 100 |

#### S 11 : List of species deemed incompatible by producers in Belt 1

| Common names or Other names | Scientific names | Botanical families | Number of plots or repetitions (n=116) | % of total citations |
|---|---|---|---|---|
| Fromager(gnin/gounga/djo) | *Ceiba pentandra* | Bombacaceae | 61 | 52,59 |
| Amien/ermien/corkpê | *Alstonia boonei* | Apocynaceae | 20 | 17,24 |
| Walê | *Cola lateritia* | Sterculiaceae | 18 | 15,52 |
| Colatier(leu/colayiri) | *Cola nitida* | Sterculiaceae | 17 | 14,66 |
| Kapokier (kpouka/gbouka/kapoté) | *Bombax buenopozense* | Bombacaceae | 14 | 12,07 |
| Parassolier(aigré/agboui/egbouin | *Musanga cecropioides* | Cecropiaceae | 9 | 7,76 |
| Samba(kpatayobouet/coffa) | *Triplochiton scleroxylon* | Sterculiaceae | 9 | 7,76 |
| Iroko(alla/ago/golyiri) | *Chlorophora excelsa/C, regia* | Moraceae | 8 | 6,90 |
| Fraké(fla/plat/yabi) | *Terminalia superba* | Combretaceae | 7 | 6,03 |
| Mouin/moin | *Cecropia peltata* | Cecropiaceae | 6 | 5,17 |
| Sessien/ahisiansian/cidian/poivre long | *Xylopia aethiopica* | Annonaceae | 5 | 4,31 |
| Pepecia/kpekpecia/avassoua/tchon | *Phyllanthus discoideus* | Euphorbiaceae | 5 | 4,31 |
| Ilomba | *Pygnanthus angolensis* | Myristicaceae | 4 | 3,45 |
| Ekoué | *Monodora sp.* | Annonaceae | 3 | 2,59 |
| Acacia | *Cassia siamea* | Cesalpiniaceae | 3 | 2,59 |
| Ako(offi/offouin) | *Antiaris africana* | Moraceae | 3 | 2,59 |
| Django/djangoe/ficus etrangleur | *Ficus goliath* | Moraceae | 3 | 2,59 |
| Manguier(amango) | *Mangifera indica* | Anacardiaceae | 2 | 1,72 |
| Baobab | *Adansonia digitata* | Bombacaceae | 2 | 1,72 |
| Framiré/tin | *Terminalia ivorensis* | Combretaceae | 2 | 1,72 |
| Dabeman/gblégblé/gbloungbloun | *Piptadeniastrum africanum* | Mimosaceae | 2 | 1,72 |
| Pamban | *Albizia zygia* | Mimosaceae | 2 | 1,72 |
| Adrain/atrin/gilo/irombo | *Pycnanthus angolensis* | Myristicaceae | 2 | 1,72 |
| Palmier | *Elaeis guineensis* | Palmaceae | 2 | 1,72 |
| Koto (baoulé) | *Sperogordia nazogarpas* | Rubiaceae | 2 | 1,72 |
| Bois bété(glonhi) | *Mansonia altissima* | Sterculiaceae | 2 | 1,72 |
| Ananyler/gborgbor/gbobo | *Voacanga africana* | Apocynaceae | 1 | 0,86 |
| Flamboyant/flambaquet | *Delonix regia* | Cesalpiniaceae | 1 | 0,86 |
| Akolê | *Discoglyprernna caloneura* | Euphorbiaceae | 1 | 0,86 |

| | | | | |
|---|---|---|---|---|
| Pê | *Discoglyprernna caloneura* | Euphorbiaceae | 1 | 0,86 |
| Dignibli | inconnu | X | 1 | 0,86 |
| Divon | inconnu | X | 1 | 0,86 |
| Ehouré | inconnu | X | 1 | 0,86 |
| Hertinin/koyago | inconnu | X | 1 | 0,86 |
| Gnicherbier | inconnu | X | 1 | 0,86 |
| Houndje | inconnu | X | 1 | 0,86 |
| Patahouhou | inconnu | X | 1 | 0,86 |
| Sitinin | inconnu | X | 1 | 0,86 |
| Tabayiri | inconnu | X | 1 | 0,86 |
| Laurenthus | *Tapinanthus bangwensis* | Loranthaceae | 1 | 0,86 |
| Tiama | *Entandrophragma angolense* | Meliaceae | 1 | 0,86 |
| Djomolohaka(baoulé) | *Albizia adianthifolia* | Mimosaceae | 1 | 0,86 |
| Ficus | *Ficus sp* | Moraceae | 1 | 0,86 |
| N'deffou/diffou | *Morus mesozygia* | Moraceae | 1 | 0,86 |
| Aniégré/ahinclin | *Aningueria robusta* | Sapotaceae | 1 | 0,86 |
| Walê | *Cola cordifolia* | Sterculiaceae | 1 | 0,86 |
| Total | 38 | 17 | 116 | 100 |

#### S 12 : List of species deemed incompatible by producers in Belt 2

| Common names or Other names | Scientific names | Botanical families | Number of plots or repetitions (n=178) | % of total citations |
|---|---|---|---|---|
| Fromager(gnin/gounga/djo/gôh) | *Ceiba pentandra* | Bombacaceae | 62 | 34,83 |
| Samba(kpatayobouet/coffa) | *Triplochiton scleroxylon* | Sterculiaceae | 56 | 31,46 |
| Mirabelle(tronma/ouinyiri) | *Spondias mombin* | Anacardiaceae | 21 | 11,80 |
| Iroko(alla/ago/golyiri) | *Chlorophora excelsa/c, regia* | Moraceae | 20 | 11,24 |
| Kapokier (kpouka/gbouka/kapoté | *Bombax buenopozense* | Bombacaceae | 15 | 8,43 |
| Ayêclê/ayinglê/ahinclin | *Napoleona leonensis* | Lechythidaceae | 12 | 6,74 |
| Walê | *Cola cordifolia* | Sterculiaceae | 10 | 5,62 |
| Ofidjedje/ | *Antiaris toxicaria* | Moraceae | 9 | 5,06 |
| Walè | *Cola gigantea* | Sterculiaceae | 9 | 5,06 |
| Amien/ermien/corkpê | *Alstonia boonei* | Apocynaceae | 8 | 4,49 |
| Dabeman(gblégblé/gbloungblou) | *Piptadeniastrum africanum* | Mimosaceae | 8 | 4,49 |
| Palmier | *Elaeis guineensis* | Palmaceae | 7 | 3,93 |
| Aya/cotibé/bois rouge | *Nesogordonia papaverifera* | Sterculiaceae | 7 | 3,93 |
| Ako/offi/offouin | *Antiaris africana* | Moraceae | 6 | 3,37 |
| Yopoui | inconnu | X | 6 | 3,37 |
| Fraké(fla/plat/yabi) | *Terminalia superba* | Combretaceae | 5 | 2,81 |
| Avocatier | *Percea americana* | Lauraceae | 5 | 2,81 |
| Goli/golikpakpa | *Albizzia spp* | Mimosaceae | 5 | 2,81 |
| Kotibé(ayiaha/ayiala/aya) | *Nesogordonia papaverifera* | Sterculiaceae | 5 | 2,81 |

| | | | | |
|---|---|---|---|---|
| Akpi | *Ricinodendron heudelotii* | Euphorbiaceae | 4 | 2,25 |
| Yêclê/yêglê/yinglè/gnêglê | *Ficus exasperata* | Moraceae | 4 | 2,25 |
| Assan | *Celtis zenkeri* | Ulmaceae | 4 | 2,25 |
| Biébiésrélé | *Spathodea campanulata* | Bignoniaceae | 3 | 1,69 |
| Ploussou(tamtam) | *Cordia platythrysa* | Boraginaceae | 3 | 1,69 |
| Lingué (lagrê ) | *Afzelia africana* | Caesalpiniaceae | 3 | 1,69 |
| Acacia | *Cassia siamea* | Cesalpiniaceae | 3 | 1,69 |
| Framiré(tin) | *Terminalia ivorensis* | Combretaceae | 3 | 1,69 |
| Néré sauvage | *Parkia bicolore* | Mimosaceae | 3 | 1,69 |
| Oranger(n'dédé) | *Citrus sinensis* | Rutaceae | 3 | 1,69 |
| Bois bété(glonhi) | *Mansonia altissima* | Sterculiaceae | 3 | 1,69 |
| Kpekpecia/tchon | *Margaritaria discoidea* | Euphorbiaceae | 2 | 1,12 |
| Kplé/kakrou/diamon/boborou | *Irvingia gabonensis* | Irvingiraceae | 2 | 1,12 |
| Wowoliwo | *Anthocleista nobilis* | Loganiaceae | 2 | 1,12 |
| Acajou | *Kaya grandifolia* | Meliaceae | 2 | 1,12 |
| Kondro | *Lannea acida* | Anacardiaceae | 1 | 0,56 |
| Hévéa sauvage/poyê | *Funtumia africana* | Apocynaceae | 1 | 0,56 |
| Serber/sêsê/sêwê/sohouê | *Holarrhena floribunda* | Apocynaceae | 1 | 0,56 |
| Blimo | Kigelia africana | Bignoniaceae | 1 | 0,56 |
| Baobab | *Adansonia digitata* | Bombacaceae | 1 | 0,56 |
| Kplé/kakrou/diamon/boborou | *Irvingia gabonensis* | Irvingiraceae | 1 | 0,56 |
| Django/djangoe/ficus etrangleur | *Ficus goliath* | Moraceae | 1 | 0,56 |
| Kpon | *Zanthoxylum gilletii* | Rutaceae | 1 | 0,56 |
| Boliwa/marcoré | *Tieghemella heckelii* | Sapotaceae | 1 | 0,56 |
| Cola grandis | *Cola grandis* | Sterculiaceae | 1 | 0,56 |
| Colatier(leu/colayiri) | *Cola nitida* | Sterculiaceae | 1 | 0,56 |
| Kototchê/porépore/lotopha | *Sterculia tragacantha* | Sterculiaceae | 1 | 0,56 |
| Total | 49 | 20 | 178 | 100 |

**S 13 : List of species deemed incompatible by producers in Belt 3.**

| Common names or Other names | Scientific names | Botanical families | Number of plots or of repetition (n=175) | % total of citations |
|---|---|---|---|---|
| Fromager | *Ceiba pentadra* | Bombacaceae | 73 | 41,44 |
| Samba | *Triplochiton scleroxylon* | Sterculiaceae | 46 | 26,52 |
| Walè | *Cola cordifolia/C.lateritia* | Sterculiaceae | 22 | 12,71 |
| Ako | *Antiaris africana* | Moraceae | 17 | 9,94 |
| Fraké | *Terminalia superba* | Combretaceae | 13 | 7,18 |
| Iroko | *Chrophora exelsa/ C. régia* | Moraceae | 12 | 6,63 |
| Koto(ouffouéwalè) | *Pterygota macrocarpa* | Sterculiaceae | 10 | 5,52 |
| Asan | *Celtis zenkeri* | Ulmaceae | 9 | 4,97 |
| Golikpakpa | *Albizzia spp* | Mimosaceae | 7 | 3,87 |

| | | | | |
|---|---|---|---|---|
| Kotibé | *Nesogordonia papaverifera* | Sterculiaceae | 7 | 3,87 |
| Bété | *Mansonia altisima* | Sterculiaceae | 7 | 3,86 |
| Amien | *Alstonia boonei* | Apocynaceae | 5 | 2,76 |
| Colatier | *Cola nitida* | Sterculiaceae | 4 | 2,21 |
| Dabema | *Piptadeniastrum africanum* | Mimosaceae | 4 | 2,21 |
| Ficus étrangleur | *Ficus goliath* | Moraceae | 4 | 2,21 |
| Lohô/logro | *Daniella oliveri* | Cesalpiniaceae | 4 | 2,21 |
| Aloma | *Ficus capensis* | Moraceae | 3 | 1,66 |
| Manguier | *Mangifera indica* | Anacardiaceae | 3 | 1,66 |
| Acacia magium | *Acacia mangium* | Mimosaceae | 2 | 1,1 |
| Acajou | *Kaya anthotheca/K grandifolia* | Meliaceae | 2 | 1,1 |
| Agboui | *Musenga cecropioides* | Moraceae | 2 | 1,1 |
| Aniégré | *Pouteria aningri* | Sapotaceae | 2 | 1,1 |
| Ficus exasperata | *Ficus exasperata* | Moraceae | 2 | 1,1 |
| Teck | *Tectona grandis* | Verbenceae | 2 | 1,1 |
| Akpi | *Ricinodendron heudelotii* | Euphorbiaceae | 1 | 0,55 |
| Flamboyant | *Dolenix regia* | Ceselpiniaceae | 1 | 0,55 |
| Gmelina | *Gmelina arborea* | Verbenaceae | 1 | 0,55 |
| Irègué | *Ficus dicranostyla* | Moraceae | 1 | 0,55 |
| Konaapi | *Canarium schweinfürthii* | Burseraceae | 1 | 0,55 |
| Kpatakpla | *Pentaclethia macrophila* | Mimosaceae | 1 | 0,55 |
| Muti | *Ficus vogeii* | Moraceae | 1 | 0,55 |
| Neem(djèkouadjo) | *Azadirachta indica* | Meliaceae | 1 | 0,55 |
| Sroka | *Erythrima senegalensis* | Papillonnaceae | 1 | 0,55 |
| Total | 33 | 15 | 175 | 100 |

**S 14 : Number of plantations/producers and surveyed areas by zone and department.**

| Cocoa production zones | Departments covered | Number of plantations / producers surveyed | Total cocoa plot area (ha) | Minimum area (ha) | Maximum area (ha) |
|---|---|---|---|---|---|
| | Abengourou | 76 | 342,5 | 0,5 | 40 |
| **1st Belt** | Adzopé | 24 | 65 | 0,5 | 8 |
| | Agboville | 16 | 53,25 | 0,5 | 9 |
| Zone total | 3 | 116 | 461,15 | 0,5 | 40 |
| | Bouaflé | 33 | 84,25 | 0,5 | 10 |
| | Daloa | 35 | 94,25 | 0,5 | 10 |
| | Djekanou | 14 | 26,5 | 0,5 | 4 |
| **2nd Belt** | Issia | 32 | 77,5 | 1 | 6 |
| | Toumodi | 23 | 75,5 | 1 | 15 |
| | Vavoua | 11 | 25 | 1 | 7 |
| | Yamoussoukro | 15 | 81,5 | 1 | 30 |
| | Zoukougbeu | 17 | 40,25 | 0,75 | 10 |
| Zone total | 8 | 180 | 504,75 | 0,5 | 30 |
| | Divo | 115 | 484 | 1 | 46 |
| **3rd Belt** | Guitry | 4 | 7 | 7 | 7 |
| | Soubré | 35 | 84,25 | 0,25 | 7 |
| | Tiassalé | 24 | 45,57 | 0,32 | 7 |
| Zone total | 4 | 178 | 620,82 | 0,25 | 46 |

**S 15 : Distribution of producers as a percentage (p.c) by ethnic origin, education level and age, per cocoa production zone**

| | 1st Belt | 2nd Belt | 3rd Belt | Moyennes |
|---|---|---|---|---|
| **Ethnic origin :** | | | | |
| Native | 76,72 | 63,13 | 33,71 | 55,53 |
| Domestic migrant | 16,38 | 20,11 | 48,57 | 29,79 |
| Foreign migrant | 6,90 | 16,76 | 17,71 | 14,68 |
| **Education level :** | | | | |
| Illiterate | 32,76 | 49,72 | 36,57 | 40,64 |
| Primary | 31,90 | 24,58 | 30,86 | 28,11 |
| Secondary | 30,17 | 24,02 | 29,14 | 28,06 |
| Tertiary | 5,17 | 1,68 | 3,43 | 3,19 |
| **Age group :** | | | | |
| ]0-20] | 0,86 | 0 | 0,57 | 0,45 |
| ]20-35] | 14,66 | 15,09 | 22,41 | 17,82 |
| ]35-50] | 52,59 | 40,25 | 39,08 | 42,98 |
| ]50-65] | 18,97 | 32,7 | 25,86 | 26,5 |
| ]65-80] | 11,21 | 10,06 | 10,34 | 10,47 |
| ]80-95] | 1,72 | 1,89 | 1,72 | 1,78 |

**S 16 : Field data collection methods for the survey forms (474) administered by the surveyors in 2013, 2014 and 2016.**

In order to properly carry out the data-collection process across the three main cocoa production zones (belts), the surveyors proceeded as follows :

**A. Diagnostic survey**

The survey was carried out in three main phases:

- Designing a questionnaire;
- Selecting the villages and producers to be surveyed;
- Administering the questionnaire by interview.

**A.1. Designing the questionnaire**

To collect data from cocoa producers, a questionnaire was designed in collaboration with CNRA's "Forest and Environment" research programme. This questionnaire combines closed, semi-closed and multiple-choice questions. It is structured in four parts:

- General information;
- Information on the cocoa plantation;
- Information on tree species associated with cocoa;
- Management and ownership of trees within cocoa AFS.

**A.2. Selection of the villages and producers surveyed**

**A.2.1. Selection of villages**

In each sub-prefecture of the cocoa-growing zone, villages were selected within the main cocoa production areas based on information gathered from ANADER. In some cases, this information came from earlier work carried out by CNRA, in particular the sites where on-farm experimental plots had been established.

**A.2.2. Selection of producers**

Surveyed producers were selected at random within the villages covered. They came from:

- Branches of certain cooperatives active in the departments;
- Participants in farmer field schools set up by ANADER.

In addition, other individual producers were selected directly in villages where information from ANADER and the cooperatives was not available.

**A.3. Conduct of the survey**

The survey was preceded by a test phase to validate the questionnaire. The test was carried out on samples of producers in villages across the three belts. The survey itself was conducted through interviews of the target population. It was carried out with technicians from CNRA Divo's Cocoa Agronomy Laboratory. Cocoa farmers were interviewed individually, either at home or on their cocoa plantations.

**A.3.1. Data collected**

Several types of information, grouped into four categories, were collected during the surveys. These were:

- General information: name, ethnic origin, education level, age of the producer;
- Agronomic characteristics of the cocoa plantations: year established, location, origin of planting material, area, output;
- Information on tree species associated with cocoa: origin, number of species, role, plant organs used, fate of the products… ;
- Management and ownership of trees within cocoa AFS.

**B. Inventory of tree species associated with cocoa**

In this study, the notion of "tree" refers to all species, other than herbaceous plants and food crops, present in the cocoa plot at the time of the inventory. It therefore includes palms as well as papaya trees.

**B.1. Choice of inventory type and plantations inventoried**

Given the definition above and the size of the cocoa plots surveyed, a full-count inventory method was used. Also known as a "tree-by-tree inventory", this type of floristic inventory covers the entire extent of the area concerned. It was assumed, a fortiori, that cocoa plots generally belong to the same environment. Plantations with an area of 5 ha or less, based on the producers' own declarations, were therefore selected. According to Kakou (2010), these declarations do not differ significantly from measured areas or yields.

**B.2. Conducting the inventory**

Conducting the inventory involved walking through the cocoa plots and collecting the following information on the trees associated with cocoa. This included:

- The common name of the species;
- The number of individuals per species;
- The stratum occupied by the tree relative to the cocoa canopy; three stratum classes were considered:
  - ✓ Lower stratum: trees below the cocoa canopy;
  - ✓ Medium stratum: trees whose crown is at the same level as the cocoa canopy
  - ✓ Upper stratum: trees whose crown is clearly above the cocoa canopy

These strata were recorded in the field by observation.

In addition, in the tree inventory plots, cocoa tree density was determined from three 10 m x 10 m quadrats. The number of cocoa trees was counted in each quadrat. The resulting mean was multiplied by 100 to give the mean density per hectare.

**B.3. Identification of the recorded species**

An initial identification of the species recorded during the interviews and inventories was carried out in the field using the field guide to the main plants listed in the flora of Taï National Park (Anonyme, 2000). This identification was then supplemented and finalized with the support of the botany laboratory of the Université Jean Lorougnon Guédé, the Centre National de Floristique of Université Félix Houphouët-Boigny, and the Département Eaux, Forêts et Environnement of INP-HB. To this end, the Flore de Côte d'Ivoire (Aubréville, 1959) and the Flore du Sénégal (Berhaut, 1967) were consulted, along with exchanges with various resource persons.

**S 17 : Useful tree species to conserve**

| Common names | Families | Scientific names |
|---|---|---|
| Amien | Apocynaceae | *Alstonia boonei* |
| Hévéa sauvage | Apocynaceae | *Funtuma africana* |
| Sohouè | Apocynaceae | *Holarrhena floribunda* |
| Fraké | Combretaceae | *Terminalia superba* |
| Framiré | Combretaceae | *Terminalia ivorensis* |
| Akpi | Euphorbiaceae | *Ricinodendron heudelotii* |
| Gbègbècia yassoua | Euphorbiaceae | *Phyllanthus discodeus* |
| Kplé | Irvingiraceae | *Irvingia gabonensis* |
| Acajou | Meliaceae | *Kaya anthotheca/K grandifolia* |

| | | |
|---|---|---|
| Tiama | Meliaceae | *Endophragma angolense* |
| Sipo | Meliaceae | *Endophragma utile* |
| Iroko | Moraceae | *Chlorophora excelsa/ regia* |
| Ficus capensis | Moraceae | *Ficus capensis* |
| Agboui | Moraceae | *Musenga cecropioides* |
| Ilomba | Myristicaceae | *Pygnanthus angolensis* |
| Tchindjè | Rutaceae | *Fagara xanthoxyloïdes* |

**S 18 : Useful and harmful trees recorded in cocoa plots**

***shade trees that host or act as reservoirs for cocoa swollen shoot virus**

| Common names | Scientific names | Families |
|---|---|---|
| Mirabellier* | *Spondias mombin* | Anacardiaceae |
| Fromager/kapokier | *Ceiba pentadra/ B. buonopozense* | Bombacaceae |
| Baobab | *Andansonia digitata* | Bombacaceae |
| Rovia | *Distemonanthus bentamianus* | Cesalpiniaceae |
| Bléblendou | *Treculia africana* | Moraceae |
| Iroko* | *Chlophora exelsa/ C. regia* | Moraceae |
| Walè | *Cola codifolia/C.lateritia* | Sterculiaceae |
| Koto | *Pterygota macrocarpa* | Sterculiaceae |
| Bété | *Mansonia altisima* | Sterculiaceae |
| Kotodjé | *Sterculia tragacantha* | Sterculiaceae |
| Samba | *Triplochiton scleroxylon* | Sterculiaceae |
| Colatier | *Cola nitada* | Sterculiceae |
| Kotibé(ayiaha) | *Nesogordina papaverifera* | Streculiaceae |

#### S 19 : Tree species, their uses and the type of plant organ used

| Common names | Genus and species | Families | Other names | Functions | | | | | | | | Plant organs used | | | | | |
|---|---|---|---|---|---|---|---|---|---|---|---|---|---|---|---|---|---|
| | | | | Shade | Fertilization | Food | Timber | Boundary marking | Medicinal | Belief/ritual | Other | Leaves | Fruits | Bark | Roots | Sap | Other |
| Anacardier | *Anacardium occidentale* | Anacardiaceae | | | | x | | x | | | | | x | | | | |
| Manguier | *Magifera indica* | Anacardiaceae | Amago | x | x | x | | | x | | x | | x | x | X | | x |
| Mirabellier | *Spondias mombin* | Anacardiaceae | | x | | x | | x | x | | | | x | | X | | |
| Corossolier | *Annona muricata* | Annonaceae | | | | x | | | | | | | x | | | | |
| Amien | *Alstonia boonei* | Apocynaceae | Amien/emien | x | x | | | | x | | | x | | x | X | | x |
| Hévéa svge | *Funtuma africana* | Apocynaceae | | x | | | | | | | | | | | | x | |
| Sohouè(#palu) | *Holarrhena floribunda* | Apocynaceae | | | | | | | x | | x | x | | x | X | | |
| Baobab | *Andansonia digitata* | Bombacaceae | | | | x | | | | | | | x | | | | |
| Fromager/kapokier | *Ceiba pentadra* | Bombacaceae | Gnin | x | x | x | x | x | x | | x | x | | x | x | | x |
| Cordia platy.(ploussou) | *Cordia platythrysa* | Boraginaceae | Plousou | | | | | x | | x | x | | | | | | x |
| Cassia siaméa | *Cassia siamea* | Cesalpiniaceae | Acacia | | | | | | x | | | x | | | | | |
| Gliricidia sepium | *Gliricidia sepium* | Cesalpiniaceae | Gliricidia | x | x | | x | x | | | | x | | | | | x |
| Logro | *Daniella oliveri* | Cesalpiniaceae | | x | | | | | | | | | | | | | |
| Tamarinier | *Tamarindus indica* | Cesalpiniaceae | | | | x | | | x | | | x | x | | | | |
| Lati | *Amphimas pterocoïpoïdes* | Cesalpiniaceae | | | | | | | | | | | | | | | |
| Rovia | *Distemonanthus bentamianus* | Cesalpiniaceae | Rovia | x | | | | | x | | | **x** | | **x** | **x** | | x |
| Fraké | *Terminalia superba* | Combretaceae | Fla | x | x | | x | | x | | x | | | x | x | | x |
| Framiré | *Terminalia ivorensis* | Combretaceae | | x | | | x | | | | | | | | | | x |

| | | | | | | | | | | | | | | | | | |
|---|---|---|---|---|---|---|---|---|---|---|---|---|---|---|---|---|---|
| Akpi | *Ricinodendron heudelotii* | Euphorbiaceae | Akpi | x | x | x | x | | x | | | | x | x | | | |
| Gbègbècia yassoua | *Phyllanthus discodeus* | Euphorbiaceae | Kpèkpècia yasoua | x | | | | | | | | | | | | | x |
| Jatropha | *Jatropha curcas* | Euphorbiaceae | | | | | | x | | x | x | | x | | | | |
| Papayer | *Carica papaya* | Euphorbiaceae | | | | x | | | x | | | x | x | | x | | x |
| Petit Cola | *Garcinia kola* | Guttiferaceae | | | | x | | x | | | x | | x | | | | |
| Kplé(kakrou) | *Irvingia gabonensis* | Irvingiraceae | | | | x | | x | | | | | x | | | | |
| Avocatier | *Persea Americana* | Lauraceae | Avocatier | x | x | x | | | x | | x | | x | | | | x |
| Abalé(tanvanwaka) | *Petersanthus macrocarpus* | Lechythidaceae | Tanvawaka | | | | x | | | | | | | | | | x |
| Napoléona(ayinglè) | *Napoleona leonensis* | Lechythidaceae | | x | | | x | | | | | | | | | | x |
| Wowoliwo | *Anthocleista nobilis/ A. procera* | Loganiaceae | | | | | | | x | | | x | | x | x | | |
| Acajou(louclou) | *Kaya anthotheca/K grandifolia* | Meliaceae | Louclou | x | | | x | | x | | | | | x | x | | x |
| Cedrela | *Cedrela spp* | Meliaceae | | x | | | | | | | x | | | | | | x |
| Neem | *Azadirachta indica* | Meliaceae | | | | | | | x | | | x | | | | | |
| Sipo | *Endophragma utile* | Meliaceae | | x | | | | | | | | | | | | | |
| Tiama | *Endophragma angolense* | Meliaceae | | x | | | x | | x | | | | | x | | | x |
| Acacia Auriculoformis | *Acacia auriculoformis* | Mimosaceae | | | x | | | | | | x | | | | | | x |
| Acacia Magium | *Acacia mangium* | Mimosaceae | | | x | | | | | | x | | | | | | x |
| Albizzia spp(goli/golikpakpa) | *Albizzia spp* | Mimosaceae | Gloikpakpa | x | | | | | x | | x | | | x | x | | x |
| Dabema | *Piptadeniastrum africanum* | Mimosaceae | Gblègblè, gbloungbloun | | | | | | | | x | | | | | | x |
| Agboui | *Musenga cecropioides* | Moraceae | Agboui/egbouin | x | | x | | | | | | | x | | | | |
| Ako | *Antiaris africana* | Moraceae | Offi/offoin | | x | x | x | x | x | | x | | | x | | | x |
| Arbre à pain | *Artoccarpus communis* | Moraceae | | | | x | | | | | | | x | | | | |
| Bléblendou | *Treculia africana* | Moraceae | Bleblendou | | | | x | x | | | | | | | | | x |
| Djomlo Djandji | *Ficus sp* | Moraceae | | x | | | | | | | | | | | | | x |
| Ficus capensis (alloma) | *Ficus capensis* | Moraceae | Aloma | | | | | | x | | | | x | x | | | |

| | | | | | | | | | | | | | | | | | |
|---|---|---|---|---|---|---|---|---|---|---|---|---|---|---|---|---|---|
| Ficus exasp (yinglè) | *Ficus exasperata* | Moraceae | Yinglè/yèglè | x | x | | | | x | | x | x | | x | x | | x |
| Ficus goliath (django) | *Ficus goliath* | Moraceae | Django | | | | | | x | | | | | | | | x |
| Ficus mucuso | *Ficus mucuso* | Moraceae | ayissassan | | | | | | | | | | | | | | |
| Iroko(Alla, Ago) | *Chlorophora excelsa/ regia* | Moraceae | | x | | | x | x | x | x | x | x | | x | x | x | x |
| Ilomba | *Pygnanthus angolensis* | Myristicaceae | | x | | | x | | x | | | | | | | x | x |
| Goyavier | *Psidium goyava* | Myrtaceae | | | | x | | | x | | | x | x | x | x | | x |
| Pomme rose | *Eugenia jambos* | Myrtaceae | | | | x | | | | | | | x | | | | |
| Cocotier | *Cocos nucifera* | Palmaceae | | | | x | | x | | | x | | x | x | x | | |
| Palmier | *Elaeis guineensis* | Palmaceae | | | x | x | | | | | x | x | x | | | x | x |
| Raphia | *Raphia hookeri* | Palmaceae | | x | | | | | | | | | | | | | x |
| Rônier | *Borassus flabellifer/ B. aethiopum* | Palmaceae | | | | | | | | | | | | | | | |
| Soka | *Erythrima senegalensis* | Papillonnaceae | | | | | | | x | | | | | x | | | |
| Badi | *Nauclea diderrichii* | Rubiaceae | Badi | | | | | | x | | | | | x | | | |
| Caféier | *Coffea canephora* | Rubiaceae | | x | | | | | | | x | | x | | | | x |
| koya | *Morida lucida* | Rubiaceae | | | | | | | x | | x | x | | x | x | | |
| Citronnier | *Citrus limon* | Rutaceae | | | | x | | | x | x | | x | x | x | x | | x |
| Mandarinier | *Citrus reculata* | Rutaceae | | | | x | | x | x | | x | x | x | x | | | x |
| Oranger | *Citrus sinensis* | Rutaceae | | x | | x | | x | x | | x | | x | x | | | x |
| Pamplemoussier | *Citrus grandis/C. paradisi* | Rutaceae | | x | | x | | x | x | | x | | x | | | | |
| Tchindjè | *Fagara xanthoxyloïdes* | Rutaceae | | | | | | | | | | | | | | | |
| Aniégré | *Pouteria aningri* | Sapotaceae | Aniégri | | | | x | | x | | | | | | | | x |
| Bété | *Mansonia altisima* | Sterculiaceae | Bété | x | | | x | | | | x | | | | | | x |
| Cola Cord(Walè) | *Cola codifolia* | Sterculiaceae | Walè | | | | | | | | | | | | | | |
| Colatier | *Cola nitada* | Sterculiaceae | | | | x | | x | x | x | x | | x | x | | | |
| Cotodjé | *Sterculia tragacantha* | Sterculiaceae | Poré poré | | | | | | x | | | | | x | x | | |

| | | | | | | | | | | | | | | | | | |
|---|---|---|---|---|---|---|---|---|---|---|---|---|---|---|---|---|---|
| Koto | *Pterygota macrocarpa* | Sterculiaceae | | x | | | x | | | | x | | | | | | x |
| Samba | *Triplochiton scleroxylon* | Sterculiaceae | | x | | | x | x | | | | | | | | | x |
| Kotibé(ayiaha) | *Nesogordina papaverifera* | Sterculiaceae | | x | | | x | | | | x | | | x | | | x |
| Asan | *Celtis zenkeri* | Ulmaceae | Asan | | | | x | x | | | x | | | | | | x |
| Teck | *Tecktona grandis* | Verbenaceae | | | | | x | x | | | x | | | | | | x |
| Tohozoué | *Newbouldia laevis* | Bignoniaceae | Tohozoué | | | | | | x | | x | | | **x** | | | |
| Djamon | | X | | | | | | | | | x | | | | | | x |
| Gboudihi | | X | | | | | x | | | | | | | | | | x |
| Néréoulosso | | X | | | | | | | | | x | | | | | | |
| Odjé | | X | | | | | | | | | | | | | | | |

#### S 20 : Full list of associated tree species in Ivorian cocoa farming.

| Vernacular names or local-language names | Scientific names (binomial) | Families |
|---|---|---|
| acacia mangium | *Acacia mangium* | Mimosaceae |
| baobab | *Adansonia digitata Linn.* | Bombacaceae |
| lingué (lagrê en baoulé) | *Afzelia africana Sm.* | Caesalpiniaceae |
| djomolohaka(baoulé) | *Albizia adianthifolia (Schumach.) W.F. Wright* | Mimosaceae |
| yatanza(baoulé) | *Albizia ferruginea (Guill. & Perr.) Benth.* | Mimosaceae |
| pamban(agni) | *Albizia zygia (DC.) J.F. Macbr.* | Mimosaceae |
| goli/golikpakpa | *Albizzia spp* | Mimosaceae |
| amien/ermien (agni,baoulé)/corkpê(attié) | *Alstonia boonei De Wild.* | Apocynaceae |
| lati | *Amphimas pteyocoipoides* | cesalpiniaceae |
| anacardier | *Anacardium occidentale* | anacardiaceae |
| ananas | *Ananas comosus* | bromeliaceae |
| aniégré(baoulé)/ahinclin(baoule) | *Aningueria robusta (A. Chev.) Aubrév* | Sapotaceae |
| corossolier | *Annona muricata* | Annonaceae |

| | | |
|---|---|---|
| wowoliwo (baoulé) | *Anthocleista nobilis* | loganiaceae |
| wowoliwo (baoulé) | *Anthocleista procera* | loganiaceae |
| ako(offi/offouin en baoulé) | *Antiaris africana* | moraceae |
| ofidjedje/(baoulé) | *Antiaris toxicaria var. africana (Engl.) C.C. Berg* | Moraceae |
| bofouin(baoule) | *Antiaris toxicaria var. welwitschii (Engl.) Corner* | Moraceae |
| arbre a pain | *Artoccarpus communis* | moraceae |
| acacia | *Auriculo formis* | mimosaceae |
| neem(djekouadjo en baoule) | *Azadirachta indica* | meliaceae |
| bambou de chine | *Bambusa vulgaris Schrad. ex J. C. Wendel.* | Poaceae (Gramineae) |
| kaha/kaya(pomme rose ou d'eau en baoulé et agni) | *Blighia sapida K. D. Koenig* | Sapindaceae |
| cligan(baoule) | *Blighia welwitschii (Hiern) Radlk.* | Sapindaceae |
| gbouka(baoule,gouro)/kapoté(attié)/kapokier (kpouka en baoulé) | *Bombax buenopozense P. Beauv.* | Bombacaceae |
| ronier | *Borassus aethiopum* | palmaceae |
| ronier | *Borassus flabellifer* | palmaceae |
| karite | *Butyrospermum paradoxum subsp. parkii (G. Don) Hepper* | Sapotaceae |
| kodou | *Carapa procera DC. De Wilde* | Meliaceae |
| papayer | *Carica papaya* | Euphorbiaceae |
| acacia | *Cassia siamea* | cesalpiniaceae |
| mouin/moin (attié) | *Cecropia peltata Linn.* | Cecropiaceae |
| cedrela | *Cedrela odorata* | meliaceae |
| fromager(gnin en baoule,gounga en more, djo en yacouba,gôh en bété,sokya) | *Ceiba pentandra (Linn.) Gaerth.* | Bombacaceae |
| assan(dida, baoule,agni,attie) | *Celtis zenkeri* | ulmaceae |
| iroko(alla,ago en baoule,golyiri en moré) | *Chlorophora excelsa/C, regia* | moraceae |
| pamplemoussier | *Citrus grandis* | rutaceae |
| citronier | *citrus limon* | rutaceae |
| pamplemoussier | *Citrus paradisi* | rutaceae |
| mandarinier | *Citrus reculata* | rutaceae |
| oranger(n'dédé en attié) | *Citrus sinensis* | rutaceae |

| | | |
|---|---|---|
| cocotier | *cocos nucifera* | palmaceae |
| cafeier | *Coffea canephora* | Rubiaceae |
| walê (baoulé) | *Cola cordifolia* | sterculiaceae |
| walè(baoule,agni) | *cola gigantea* | sterculiaceae |
| cola grandis | *Cola grandis* | Sterculiaceae |
| cola sauvage(toutou eb bété) | *Cola heterophylla (P. Beauv.) Schott & Endl.* | Sterculiaceae |
| walê (agni) | *Cola lateritia* | sterculiaceae |
| colatier(leu en attié,colayiri en malinke) | *Cola nitida* | sterculiaceae |
| ploussou(tamtam en baoule, agni) | *Cordia platythrysa* | boraginaceae |
| loho/logro (abbey) | *Daniella oliveri* | cesalpiniaceae |
| flamboyant/flambaquet(agni) | *Delonix regia* | cesalpiniaceae |
| pê (attie) | *Discoglyprernna caloneura (Pax) Prain* | Euphorbiaceae |
| akolê (attie) | *Discoglyprernna caloneura (Pax) Prain* | Euphorbiaceae |
| movindji | *Distemonanthus benthamianus Baill* | Caesalpiniaceae |
| palmier | *Elaeis guineensis* | palmaceae |
| kosipo(gouro) | *Entandophragma candollei* | meliaceae |
| tiama(agni,baoule,attié) | *Entandrophragma angolense* | meliaceae |
| aboudikro(baoulé) | *Entandrophragma cylindricum (Sprague) Sprague* | Meliaceae |
| doukouman (baoulé) | *Entandrophragma spp* | sterculiaceae |
| sipo | *Entandrophragma utile* | meliaceae |
| soka/sroka | *Erythrima senegalensis* | papillonnaceae |
| pomme citerne | *Eugenia jambos* | myrtaceae |
| tchindjè( agni, baoule, bete, dida) | *Fagara xanthoxyloides* | rutaceae |
| aloma (baoule) | *Ficus capensis* | moraceae |
| prêguè | *Ficus dicranostyla* | moraceae |
| yêclê/yêglê/yinglè/gnêglê(baoulé, agni) | *Ficus exasperata* | moraceae |
| django/djangoe (baoulé, agni, attié)/ficus etrangleur | *Ficus goliath* | moraceae |
| lokpro/lokplo/loungblo/avissanssan(baoulé) | *Ficus mucuso Welw. ex Ficalho* | Moraceae |
| djombo djandji | *Ficus sp* | moraceae |

| | | |
|---|---|---|
| muti(abbey) | *Ficus vogeii* | moraceae |
| hevea sauvage/poyê (baoulé) | *Funtumia africana (Benth.) Stapf* | Apocynaceae |
| petit cola | *Gaycinia kola* | guttiferaceae |
| glyricidia | *Gliricidia sepium* | cesalpiniaceae |
| gmelina(mena en baoulé) | *Gmelina arborea Roxb.* | Verbenaceae |
| bossé | *Guarea cedrata* | meliaceae |
| bossé | *Guarea thompsoneii* | meliaceae |
| niangon (attié) | *Heritiera utilis* | inconnu |
| hevea | *Hevea brasiliensis (Kunth) Müll.Arg* | Euphorbiaceae |
| serber/sêsê/sêwê/sohouê(baoulé) | *Holarrhena floribunda (G. Don) Dur. & Schinz var. floribunda* | apocynaceae |
| kplé/kakrou/diamon/boborou (abbey) | *Irvingia gabonensis* | irvingiraceae |
| jatropha | *Jatropha curcas* | Euphorbiaceae |
| acajou(louclou) | *Kaya anthothéca* | meliaceae |
| acajou | *kaya grandifolia* | meliaceae |
| kondro (baoule) | *Lannea acida A. Rich.* | Anacardiaceae |
| pistache(n'katé en attié) | *Luffa cylindrica (L.) M. Roem.* | Cucurbitaceae |
| calebassier | *luffa sp* | Cucurbitaceae |
| manguier(amango en attié) | *Mangifera indica* | anacardiaceae |
| manioc | *Manihot esculenta Crantz* | Euphorbiaceae |
| bois bété(glonhi en bété) | *Mansonia altissima* | sterculiaceae |
| wodjé (baoulé)/akêdê/angama(baoulé,dida)/kakahikpé(baoule)/odjé (baoule) | *Mareya micrantha (Benth.) Müll. Arg.* | Euphorbiaceae |
| kpekpecia (baoulé)/tchon(attie) | *Margaritaria discoidea ( Baill . ) Webster* | Euphorbiaceae |
| gni (attié) | *Milicia regia* | moraceae |
| ekoué(agni) | *Monodora sp.* | Annonaceae |
| koya(baoulé) | *Morinda lucida Benth.* | Rubiaceae |
| moringa | *Moringa oleifera Lam* | Moringaceae |
| n'deffou( en abbey)/diffou(baoulé) | *Morus mesozygia Stapf ex A. Chev.* | Moraceae |
| banane plantain | *Musa paradisiaca Linn.* | Musaceae |
| banane douce | *Musa sapientum L.* | Musaceae |

| | | |
|---|---|---|
| parassolier(aigré en baoulé,agboui/egbouin)/mouin/moin (attié) | *Musanga cecropioides R. Br.* | Cecropiaceae |
| trikpor | *Myrianthus arboreus* | Cecropiaceae |
| ayêclê/ayinglê/ahinclin(baoulé) | *Napoleona leonensis* | lechythidaceae |
| badi (dida) | *Nauclea diderrichii (De Wild.& T. Durand) Merr.* | Rubiaceae |
| kotibé(ayiaha/ayiala en baoulé,attié,agni/aya (agni) | *Nesogordonia papaverifera (A. Chev.) R. Capuron* | Sterculiaceae |
| aya/cotibé/bois rouge | *Nesogordonia papaverifera (A. Chev.) R. Capuron* | Sterculiaceae |
| tohozoué (baoule)/bodjé(baoulé) | *Newbouldia laevis* | Bignoniaceae |
| éffé(baoulé) | *Olyra latifolia Linn.* | Poaceae (Gramineae) |
| néré sauvage | *Parkia bicolore* | mimosaceae |
| kpatakpla(baoule) | *Pentaclethia macrophila* | Mimosaceae |
| avocatier | *Percea americana* | lauraceae |
| abale(moré)/tanvanwaka | *Peteysanthus macrocarpus* | lechythidaceae |
| pepecia/kpekpecia/gbegbeciavassoua(baoule,agni)/tchon | *Phyllanthus discoideus* | Euphorbiaceae |
| agnissanssan(agni) | *Pilper guineesis* | Moraceae |
| dabeman(gblégblé,gbloungbloun en baoulé) | *Piptadeniastrum africanum* | mimosaceae |
| aniégré(baoulé)/ahinclin(baoule) | *Pouteria aningeri Baehni* | Sapotaceae |
| goyavier | *Psidium guajava Linn.* | Myrtaceae |
| koto (ouffoué en agni) | *Pterygota macrocarpa* | sterculiaceae |
| adrain/atrin (baoulé)/gilo(attie)/irombo(agni,baoule) | *Pycnanthus angolensis (Welw.) Warb* | Myristicaceae |
| ilomba | *Pygnanthus angolensis* | myristicaceae |
| dédé/bambou | *Raphia hookeri G. Mann & H. Wendl.* | Arecaceae |
| raphia | *Raphia hookeri G. Mann & H. Wendl.* | palmaceae |
| n'décherie(attie,abbey) | *Rauvolfia vomitoria Afzel.* | Apocynaceae |
| akpi(baoulé,agni,attié,abbey) | *ricinodendron heudelotii* | euphorbiaceae |
| koumossi (baoulé) | *Solanum rugosum Dun.* | Solanaceae |
| biébiésrélé (baoulé) | *Spathodea campanulata P. Beauv.* | Bignoniaceae |
| koto (baoulé) | *Sperogordia nazogarpas* | Rubiaceae |
| mirabelle(tronma en baoulé, ouinyiri en gouro) | *Spondias mombin* | anacardiaceae |

| | | |
|---|---|---|
| mlémlédou/blebleadou(baoulé) | *Sterculia africana* | Sterculiaceae |
| kototchê/poréporé(baoule)/lotopha(moré) | *Sterculia tragacantha* | sterculiaceae |
| bois de tamtam(pili en bété,lilituê en yacouba) | *Stylopigma lilogeii* | inconnu |
| tamarinier/tomy | *Tamarindus indica* | cesalpiniaceae |
| laurenthus | *Tapinanthus bangwensis (Engl. & K. Krause) Danser* | Loranthaceae |
| teck | *Tecktona grandis* | verbenaceae |
| koman | *Terminalia glaucescens* | Combretaceae |
| framiré/tin(attie) | *Terminalia ivorensis A. Chev.* | Combretaceae |
| koman | *Terminalia scimperiana Hochst.* | Combretaceae |
| fraké(fla en baoulé,plat en gouro)/yabi(attié) | *Terminalia superba* | combretaceae |
| boliwa(baoule)/marcoré | *Tieghemella heckelii Pierre ex A. Chev.* | Sapotaceae |
| bleblendou | *Treculia africana* | moraceae |
| samba(kpatayobouet en baoulé)/coffa(attié) | *Triplochiton scleroxylon* | sterculiaceae |
| karite | *Vitellaria paradoxa C. F. Gaertn.* | Sapotaceae |
| ananyler/gborgbor/gbobo(attie) | *Voacanga africana Stapf* | Apocynaceae |
| sessien/ciciana/ahisiansian/cidian (baoulé)/poivre long | *Xylopia aethiopica (Dunal) A. Rich.* | Annonaceae |
| kanin finh(attié,abbey) | *Xylopia elliotii Engl. & Diels* | Annonaceae |
| kpon (attie) | *Zanthoxylum gilletii (De Wild.) P. G. Waterman* | Rutaceae |
| tchindjé(agni) | *Zanthoxylum Zanthoxyloides (Lam.) Zepern. & Timler* | Rutaceae |
| afia/afian(baoule) | *inconnu* | inconnu |
| ago(baoule,dida,bete) | *inconnu* | inconnu |
| akoi(attie) | *inconnu* | inconnu |
| aonialé(agni) | *inconnu* | inconnu |
| bossoman(agni) | *inconnu* | inconnu |
| botchôliê(baoulé) | *inconnu* | inconnu |
| dignibli(agni) | *inconnu* | inconnu |
| divon(baoulé) | *inconnu* | inconnu |
| djanon | *inconnu* | inconnu |

| | | |
|---|---|---|
| doman(baoule) | *inconnu* | inconnu |
| ehouré(agni) | *inconnu* | inconnu |
| flabougnan(agni) | *inconnu* | inconnu |
| gboudihi(dida) | *inconnu* | inconnu |
| gnangnoui | *inconnu* | inconnu |
| gnicherbier (attié) | *inconnu* | inconnu |
| gnizazui | *inconnu* | inconnu |
| gnopoui(dida) | *inconnu* | inconnu |
| guigui(bété) | *inconnu* | inconnu |
| heramba(agni) | *inconnu* | inconnu |
| houndje(abbey) | *inconnu* | inconnu |
| irêguê(baoule) | *inconnu* | inconnu |
| kondré | *inconnu* | inconnu |
| kouêmankouê(wan) | *inconnu* | inconnu |
| kpamouhi(bete) | *inconnu* | inconnu |
| kpiniman(baoule) | *inconnu* | inconnu |
| monin(attié) | *inconnu* | inconnu |
| nêrêhiri(worodougou) | *inconnu* | inconnu |
| néréoulosso | *inconnu* | inconnu |
| patahouhou(agni) | *inconnu* | inconnu |
| sama(bété) | *inconnu* | inconnu |
| sitinin (agni) | *inconnu* | inconnu |
| tabayiri(malinke) | *inconnu* | inconnu |
| tarri(dida) | *inconnu* | inconnu |
| tiklitissou | *inconnu* | inconnu |
| tongatiga(more) | *inconnu* | inconnu |
| yapa(bete) | *inconnu* | inconnu |
| hertinin(koyago) | *inconnu* | inconnu |
